# Spatial and epigenetic programs underlie heterogeneous cellular senescence in the aging liver

**DOI:** 10.64898/2026.11.18.767706

**Authors:** Sainath Mamde, Kelly Yichen Li, Marcos G. Teneche, Jacqueline Olness, Elizabeth Smoot, Colin Kern, Karl N. Miller, Jesus Flores, Zoey Zhao, Pik Ki Lau, Bing Yang, Adarsh Rajesh, Aaron P. Havas, Jacinta Lucero, Jovina Djulamsah, Qian Yang, Charlene Miciano, Jessica Proulx, Rouven Arnold, Michael Alcaraz, Andrew Davis, Armin Gandhi, Laurence Haddadin, Seoyeon Lee, Xiaomeng Gao, Christina Huan Shi, Irene Huang, Carolina Cano Macip, Shanshan Yin, Hillina Berhane, Sha Li, Zong Ming Chua, Bin Li, Amy Li, Kenneth Kim, Diana Jurk, João F. Passos, Quan Zhu, Allen Wang, Bing Ren, Peter D. Adams, Kevin Y. Yip, Nathan R. Zemke

## Abstract

Aging is a complex biological process leading to tissue dysfunction, and driven in part by progressive epigenetic drift, widespread transcriptional dysregulation and cellular senescence. Senescent cells (SnCs) accumulate across tissues in a heterogeneous and cell type-specific manner, yet their molecular signatures and spatial organization within native tissue have not been systematically mapped. Here we combined single-nucleus transcriptomic, epigenomic, three-dimensional (3D) genome and spatial profiling to systematically characterize aging of the mouse liver and associated cellular senescence. Aging was marked by hepatocyte loss, immune cell expansion, erosion of repressive H3K9me3 heterochromatin, alteration of 3D genome organization, and activation of stress-response pathways known to promote senescence. Age-associated SnCs were spatially colocalized within discrete inflammatory microenvironments termed SenSpots. Different SenSpots were dominated by distinct hepatocyte senescence states (senotypes), driven by unique transcription factors, and exhibited distinct lobular zonation. Single-cell DNA methylation profiling revealed a subset of immune cells in aged livers with loss of DNA methylation at constitutive heterochromatin and elevated senescence signatures. AAV-mediated, p21-promoter driven *Stat1* deletion in the aged liver selectively suppressed interferon-responsive senescence programs while pharmacological inhibition of MDM2 attenuated senescence-associated inflammatory signatures. Together, these findings reveal that hepatic senescence is not a uniform state but a set of diverse epigenetically reprogrammed, spatially organized, and pharmacologically modifiable inflammatory niches.

## Introduction

Aging is a complex biological process accompanied by chromatin remodeling^1,2^ and widespread alterations in gene expression that lead to tissue dysfunction.^3^ Together with other hallmarks of aging, cellular senescence is linked to this age-related tissue dysfunction.^4^ Cellular senescence is a stress response in which cells enter a stable proliferation arrest limiting their expansion.^5^ In vitro, this cell-cycle arrest is typically enforced by the accumulation of cyclin-dependent kinase inhibitors (CDKIs), including CDKN1A (p21) and CDKN2A (p16^INK4a^).^5–7^ Senescent cells (SnCs) remain metabolically active and can release a mix of inflammatory cytokines, chemokines and other bioactive molecules, collectively referred to as the senescence-associated secretory phenotype (SASP).^8–10^ Although senescence can suppress tumor development via proliferation arrest and immune surveillance,^11^ accumulation of SnCs with age can impair tissue function and promote chronic inflammation.^12–14^ Genetic clearance of p16^Ink4a^-positive cells extends median lifespan in naturally aged mice,^12^ while pharmacological senolysis can improve stem-cell function^15^ and physical function and remaining survival in old mice.^16^

The liver plays a central metabolic role, and its hepatocytes, together with other liver cell types, are spatially organized into distinct functional zones.^17^ With age, the liver develops steatosis, low-grade fibrosis and chronic inflammation,^18,19^ and single-cell atlases have linked these changes to shifts in hepatocyte zonation and cellular identity.^20–23^ Several studies have implicated senescence in these processes: senescence of hepatic stellate cells (HSCs) has been proposed to limit fibrosis after injury,^24^ whereas SnCs also contribute to age-dependent hepatic steatosis.^18^ In addition, depletion of p16^High^ liver sinusoidal endothelial cells (LSECs) has been reported to disrupt blood-tissue barriers, leading to perivascular fibrosis.^25^ These findings show that the effects of senescence depend on cell type and context, but how different SnC types accumulate in the liver during natural aging and contribute to its functional decline remains poorly understood.

SnCs are heterogeneous, and no single marker can reliably identify them across organs.^5,26,27^ Commonly used markers, such as CDKN1A, CDKN2A, decreased perinuclear lamin B1, γH2AX foci and SASP factors, are not individually reliable across contexts because not all SnCs express a given marker, and non-senescent cells can express it in response to other stresses.^26,28,29^ Spatial context also matters: CDKN1A-positive hepatocytes are enriched periportally in younger human livers, with broader distributions in older or more fibrotic livers.^22,30^ As a result, estimates of SnC abundance depend on the markers and methods used, and senescence has to be defined by combinations of features rather than any single one.

Beyond marker expression, senescence is accompanied by changes in chromatin organization and 3D genome architecture.^31^ Heterochromatin regions marked by H3K9me3 and H3K27me3 reorganize within the nucleus,^32,33^ while senescence-associated genes can undergo changes in compartment organization and long-range chromatin contacts.^34^ However, these epigenetic features of senescence have been largely studied in cultured fibroblasts^31,34^ and remain poorly profiled at the single-cell level during liver aging. A major open question is whether these epigenetic changes are linked to the identity, molecular state and tissue location of individual senescent cells in the aging liver. Integrating measurements of transcription, chromatin state, DNA methylation and genome architecture with spatial mapping in the same aging tissue can help connect senescent cell states to their regulatory programs and local environments, and thereby identify candidate regulators that establish or sustain cellular senescence in vivo.

Here we combine single-nucleus transcriptomic, epigenomic and 3D genome profiling with spatial single-cell technologies to systematically characterize liver aging and SnCs in young (4 months), old (22 months) and very old (30 months) mice. Our data show that liver aging is marked by loss of hepatocytes and expansion of the immune cell compartment. To identify SnCs, we develop SenLearner, a machine-learning classifier initialized by combining senescence marker gene sets^35–37^ with curated p53 target genes^38^ and CDKI genes to identify additional senescence-related genes. Among the gene regulatory networks associated with senescence in hepatocytes, knockout of STAT1, a predicted regulator of one of these networks, in p21-Cre targeted cells reduces senescence features and improves liver metabolism. Hepatocyte SnCs further resolve into distinct senotypes that colocalize spatially to form SenSpots, and stabilization of p53 with the MDM2 inhibitor HDM201 suppresses specific senotypes and attenuates senescence-associated signatures.

## Results

### Cellular and gene regulatory dynamics in the aging liver

To characterize how aging and senescence reshape the liver, we combined single-cell profiling of gene expression, chromatin accessibility, histone modifications, DNA methylation and 3D genome architecture with spatial transcriptomics, in young (∼4 months), old (∼22 months) and very old (∼30 months) mice, balanced for sex within each age group (Fig. 1A, Supplementary Table 1, Supplementary Table 2). Each assay captured a distinct layer of regulation: 10x Multiome measured RNA and chromatin accessibility in the same nucleus (n = 18 mice), snm3C-seq measured DNA methylation together with chromosome conformation capture (3C) (n = 14 mice), Paired-Tag measured histone modifications (H3K27ac, H3K27me3 and H3K9me3), each alongside RNA in separate libraries (n = 12 mice), and MERFISH imaged individual transcripts within intact tissue sections (n = 19 mice) (Fig. 1B).

**Fig. 1:**
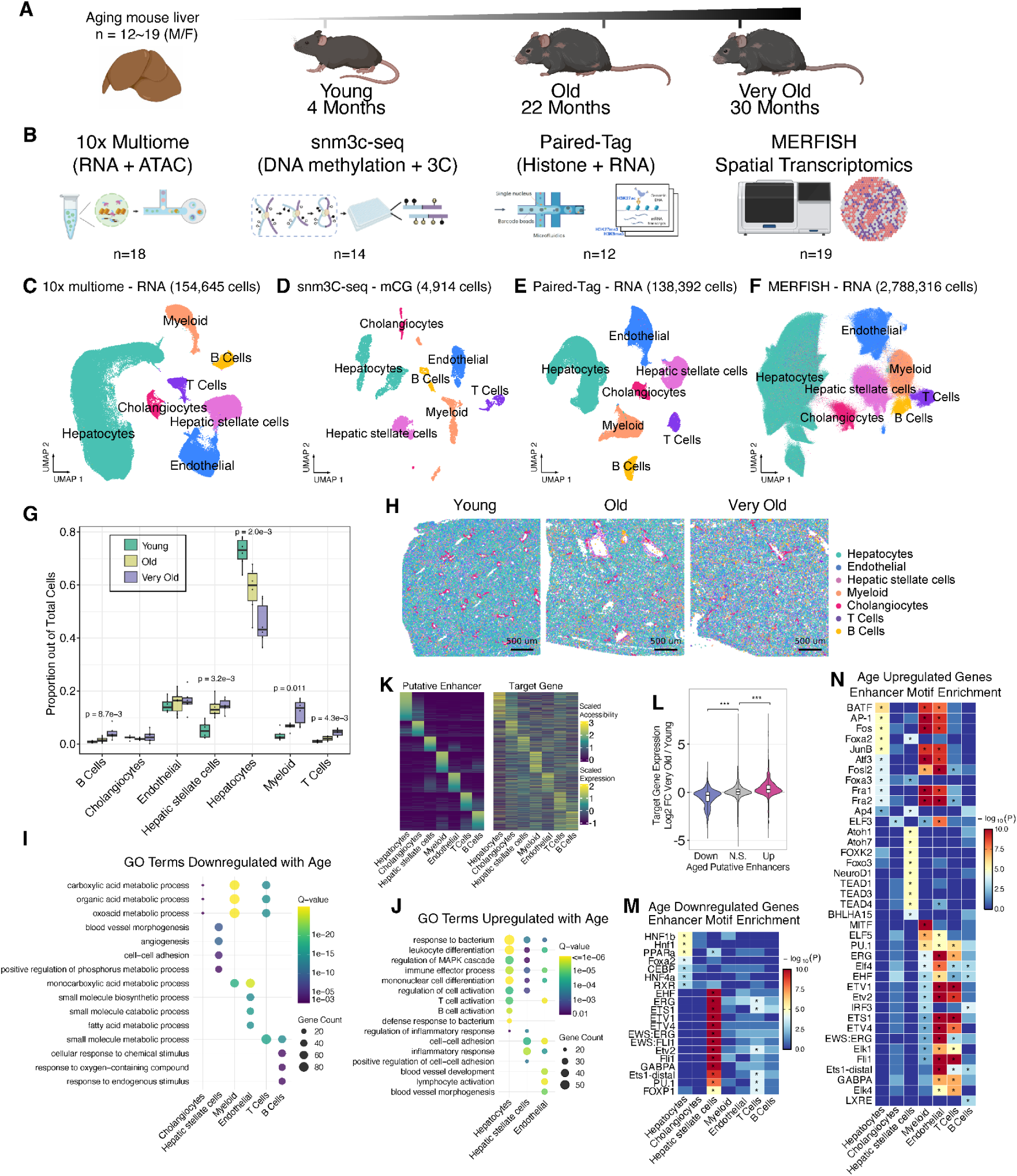
A single-cell multimodal atlas of the aging mouse liver. **A**, Experimental design (BioRender). Liver was collected at around 4 months (young), 22 months (old) and 30 months (very old); n = 12∼19 mice, sex-balanced. **B**, The four assays used (BioRender), with the number of mice profiled by each: 10x Multiome (RNA and chromatin accessibility from the same nucleus), snm3C-seq (DNA methylation and chromosome conformation(3C)), Paired-Tag (one histone modification, H3K27ac, H3K27me3 or H3K9me3, together with RNA) and MERFISH. **C** to **F**, Uniform manifold approximation and projection (UMAP) of 10x Multiome RNA (**C**), snm3C-seq mCG (**D**), Paired-Tag RNA (**E**) and MERFISH (**F**), colored by cell type. Number of nuclei or cells after quality control is shown above each panel. **G**, Cell type proportions in the 10x Multiome data by age group. Box, median and IQR; whiskers, 1.5 × IQR; points, individual mice (n = 6 per age group). P values are shown for young versus very old where P < 0.05 (Wilcoxon rank-sum test). **H**, MERFISH sections from one mouse per age group, colored by cell type. Scale bars, 500 µm. **I**, **J**, Gene ontology biological processes enriched among genes downregulated (**I**) or upregulated (**J**) with age in each cell type (edgeR, very old versus young, FDR < 0.05). Dot size, number of genes; color, Benjamini-Hochberg adjusted q value. Top terms in each cell type are shown. **K**, Chromatin accessibility of ABC-predicted putative enhancers (left) and expression of their top-scoring target genes (right), scaled by row. Rows are ordered by the accessibility within each cell type. **L**, Target gene expression change with age (log2 fold change, very old versus young) for genes linked to putative enhancers that gained (Up), lost (Down) or did not change (N.S.) accessibility. ***P < 0.001, Wilcoxon rank-sum test. **M**, **N**, Transcription factor (TF) motifs enriched in putative enhancers of age-downregulated (**M**) and age-upregulated (**N**) genes, by cell type. Color, −log10(P); asterisks, q < 0.05.

After quality filtering (Methods), the dataset contained 154,645 10x Multiome nuclei, 4,914 snm3C-seq nuclei, 138,392 Paired-Tag nuclei and 2,788,316 MERFISH cells (Fig. 1C–F, Supplementary Fig. 1A–E). Clusters were assigned to cell types on the basis of canonical marker expression, identifying hepatocytes, cholangiocytes, hepatic stellate cells, endothelial cells, myeloid cells, T cells and B cells in every modality (Fig. 1C–F and Supplementary Fig. 2A–G). This atlas therefore links the transcriptional, epigenetic, genome folding, and spatial organization of each liver cell type across the lifespan.

Aging markedly remodeled the cellular composition of the liver. The proportion of hepatocytes declined from young to very old mice (P = 2.0 × 10⁻³), accompanied by expansion of hepatic stellate cells (P = 3.2 × 10⁻³), T cells (P = 4.3 × 10⁻³), B cells (P = 8.7 × 10⁻³) and myeloid cells (P = 0.011) (Wilcoxon rank-sum test; Fig. 1G). MERFISH confirmed an absolute increase in immune cell density in very old mice (Fig. 1H and Supplementary Fig. 2H), with immune and stromal cells increasingly interspersed throughout the parenchyma.

To define the transcriptional programs that accompany these changes, we performed pseudobulk differential expression analysis between very old and young mice within each cell type (edgeR,^39^ FDR < 0.05; Supplementary Table 3). Age-associated changes were identified in each cell type: downregulated genes were largely cell type-specific, whereas upregulated genes converged on shared inflammatory programs (Fig. 1I,J). In endothelial cells, genes downregulated with age were enriched for fatty acid and small-molecule metabolism, processes central to the role of liver sinusoidal endothelium in lipid transport between blood and hepatocytes,^40,41^ whereas upregulated genes were enriched for cell–cell adhesion and inflammatory response. Hepatic stellate cells, the liver’s resident pericytes,^42^ showed reduced expression of genes involved in angiogenesis and blood vessel morphogenesis. Because these cells communicate with endothelial cells to maintain liver homeostasis,^43^ this loss may contribute to the sinusoidal dysfunction reported in the aged liver.^44^ Myeloid cells downregulated genes of carboxylic and organic acid metabolism, and B cells downregulated genes involved in responses to chemical and endogenous stimuli (Fig. 1I, Supplementary Table 4). Genes upregulated with age were instead enriched for immune and inflammatory processes, including leukocyte differentiation, T and B cell activation, MAPK signalling, responses to bacteria and regulation of the inflammatory response, in hepatocytes, hepatic stellate cells and endothelial cells (Fig. 1J, Supplementary Table 4). Together with the expansion of immune cells, these findings indicate that aging is accompanied by increased inflammation across the liver, extending to parenchymal and stromal cells, consistent with chronic inflammation as a hallmark of aging.^4,45^

To identify the regulatory elements underlying these programs, we used the activity-by-contact (ABC)^46^ model with accessible chromatin regions to identify putative enhancers and predict their target genes in each cell type (Supplementary Table 5). The resulting putative enhancers showed largely cell type-specific accessibility, with matching expression of their target genes (Fig. 1K). Age-associated changes were concordant between putative enhancers and their target genes (Fig. 1L). Motif analysis of these putative enhancers revealed cell type-specific regulators. Among enhancers of age-downregulated genes, those in hepatocytes were enriched for motifs of hepatocyte identity and metabolic regulators, including HNF4α, HNF1, FOXA2, C/EBP and PPARα, whereas those in hepatic stellate cells were dominated by ETS family motifs (ERG, FLI1, ETV2, ETS1) (Fig. 1M). Enhancers of age-upregulated genes were instead enriched for AP-1 motifs (BATF, Fos, JunB, Atf3, Fra1, Fra2) in hepatocytes, myeloid cells and endothelial cells, alongside motifs restricted to particular cell types, including IRF3 in myeloid and B cells, MiT/TFE (MITF) in myeloid cells, and TEAD and FOXO3 in hepatic stellate cells (Fig. 1N), the former implicating YAP/TAZ signalling, a driver of stellate cell activation.^43,47^ The recurrence of AP-1, which integrates stress and inflammatory signalling, across otherwise distinct enhancer landscapes points to a shared regulatory route to the age-associated inflammatory shift.^48^ Together, these results show that liver aging is accompanied by hepatocyte decline, expansion of immune and stellate cells, and increased inflammation that extends to parenchymal and stromal cells, alongside a cell type-specific loss of metabolic programs.

### Global Loss of epigenetic information and 3D genome organization in the aging liver

Loss of transcriptional fidelity is a feature of aging that may erode cell type-specific gene expression programs and cellular homeostasis.^49^ To assess transcriptional drift, we quantified cell-to-cell variability in gene expression using manifold drift^50^ and observed a significant age-related increase across cell types (Fig. 2A). We then asked whether this increased transcriptional drift reflects a broader deterioration of the gene regulatory architecture.^51,52^ Examining genome organization in hepatocytes from snm3C-seq data, we found an age-associated increase in trans-chromosomal contacts (Fig. 2B–C, Supplementary Fig. 3A) and a corresponding decrease in intra-TAD contacts (Fig. 2D–F, Supplementary Fig. 3B). Together, these findings suggest that the structural compartmentalization of the genome, which normally constrains enhancer–promoter communication,^53^ is progressively disrupted during aging.

**Fig. 2:**
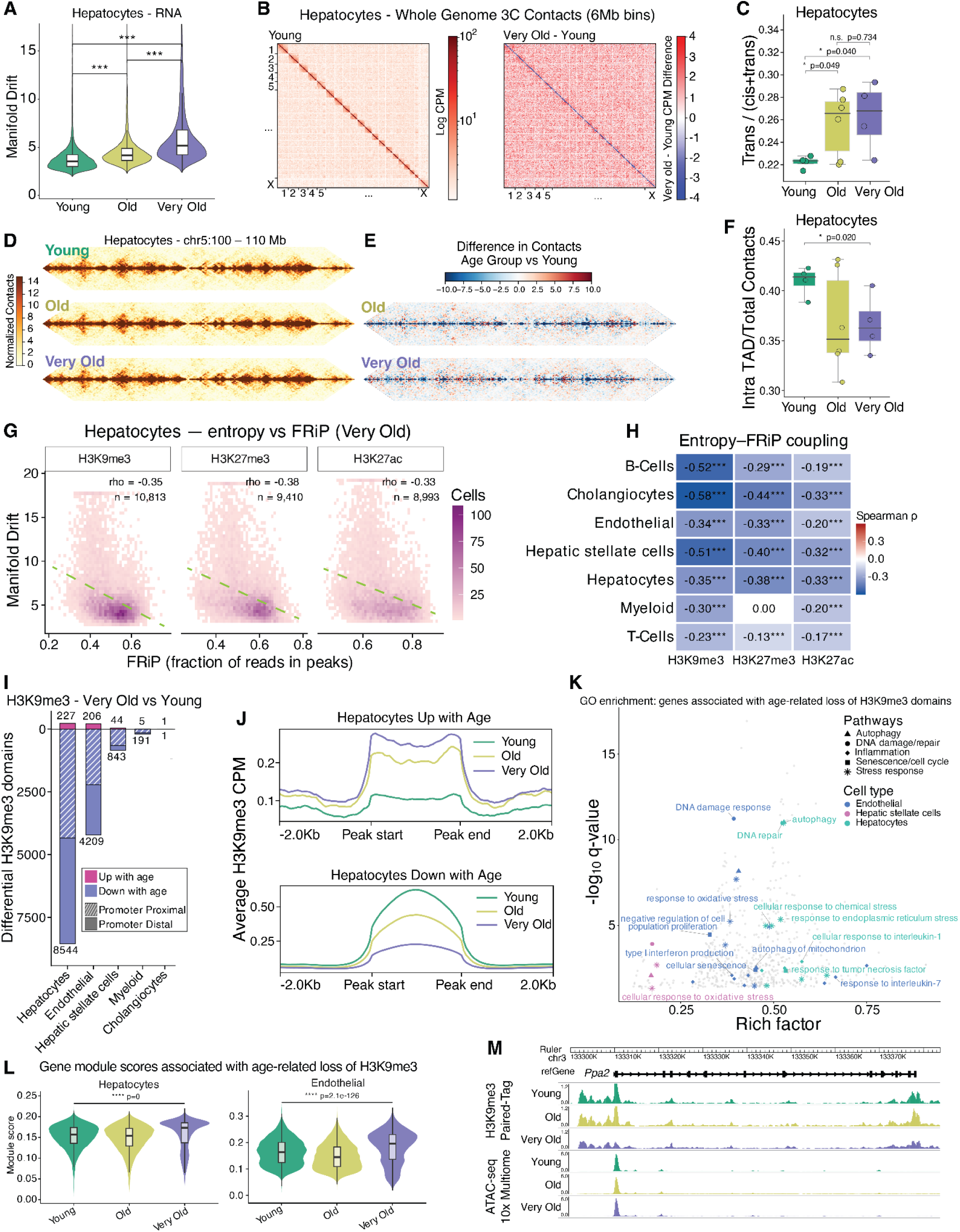
Age-associated erosion of epigenetic and three-dimensional genome organization. **A**, Manifold drift of hepatocyte nuclei by age group, defined as the mean distance to the 20 nearest young nuclei in Harmony space from 10X Multiome. Violin, distribution across nuclei; box, median and IQR. ***P < 0.001, Wilcoxon Rank Sum-Test. **B**, Genome-wide 3C contacts in 6-Mb bins in hepatocytes: young mice (left, log CPM) and the very old minus young (right). **C**, Fraction of hepatocyte contacts that were trans-chromosomal, by age group. Points, individual mice; box, median and IQR; whiskers, 1.5 × IQR. T-Test; n.s., not significant. **D**, **E**, Hepatocyte contact matrices at chr5:100–110 Mb for each age group (**D**) and the difference relative to young (**E**). **F**, Fraction of hepatocyte contacts falling within topologically associating domains (TADs), by age group; plotted and tested as in **C**. **G**, Density of manifold drift against the fraction of reads in peaks (FRiP) across individual very old hepatocyte nuclei, one panel per histone modification. Color, nuclei per bin; dashed line, linear fit; rho, Spearman correlation. **H**, Spearman correlation between manifold drift and FRiP for each cell type and histone modification. ***P < 0.001. **I**, H3K9me3 domains gained and lost in very old relative to young mice in each cell type (edgeR, FDR < 0.05), split by promoter overlap (transcription start site ± 1 kb). **J**, Mean H3K9me3 signal over domains gained (top) and lost (bottom) with age in hepatocytes, extended 2 kb either side, for each age group. **K**, Gene ontology biological processes enriched among genes near age-lost H3K9me3 domains. Colored points, terms with q < 0.05, by pathway category and cell type; gray points, all other tested terms. Rich factor, the fraction of a term’s genes present in the input set. **L**, Module scores for the genes in **K** in 10x Multiome hepatocytes and endothelial cells. Violin, distribution across cells; box, median and IQR. Wilcoxon rank-sum test, young versus very old. **M**, H3K9me3 Paired-Tag and 10x Multiome ATAC-seq signal at *Ppa2* (chr3:133,300,000–133,375,000) by age group.

Epigenetic erosion has been proposed as a mechanistic driver of transcriptional drift.^54^ To quantify epigenetic erosion at single-cell resolution, we measured the fraction of reads in peaks (FRiP) from H3K9me3, H3K27me3, and H3K27ac Paired-Tag data. For all three marks and across cell types, single-cell FRiP values were negatively correlated with manifold drift scores (Fig. 2G–H), linking the erosion of histone modifications at the single-cell level to increased transcriptional noise.

H3K9me3 marks constitutive heterochromatin^55^ and is a critical repressor of transposable elements.^56^ In the aging liver, we identified widespread domains of reduced H3K9me3 signal, most prominently in hepatocytes, endothelial cells, and hepatic stellate cells (Fig. 2I–J, Supplementary Fig. 3C, Supplementary Table 6). A substantial fraction of these age-associated H3K9me3 losses occurred within ±1 kb of a transcription start site (TSS). Genes associated with these promoter-proximal H3K9me3 losses were enriched for pathways including autophagy, DNA damage repair, and cellular senescence (Fig. 2K), consistent with reports that heterochromatin loss derepressed stress-response programs^57^ and were significantly upregulated with age (Fig. 2L). For example, *Ppa2*, which promotes mitophagy under cellular stress, exhibited promoter-proximal loss of H3K9me3 and increased expression in aged hepatocytes (Fig. 2M).

### Characterization of senescence transcriptional signatures and spatial organization

We next sought to identify senescent cells in the aging mouse liver. This is challenging because senescence is a heterogeneous cell state with no single defining marker.^5,26^ Cells with detectable RNA from common senescence marker genes, such as *Cdkn2a* and *Cdkn1a*, did not form discrete, age-associated clusters in UMAP space when using standard snRNA-seq clustering methods (Supplementary Fig. 4A). Moreover, given the inherent sparsity of snRNA-seq detection and the lack of established senescence signatures for individual liver cell types, we could not reliably classify cells as senescent based on the presence of specific marker genes alone. To overcome these limitations, we developed SenLearner, an interpretable machine-learning framework that learns cell type-specific senescence classifiers and assigns a senescence probability to every cell (Supplementary Table 7). We first designed a custom panel of 147 senescence-associated genes, termed the San Diego Tissue Mapping Center (SDTMC) gene set, from five sources: four senescence marker gene sets (SenMayo,^35^ Fridman,^36^ SenSig^37^ and an in-house Canonical IFN & SASP set (Methods)) and a curated set of p53 target genes.^38^ Genes present in at least two of these sources were retained, together with all seven mammalian cyclin-dependent kinase inhibitor genes (Supplementary Fig. 4B; Methods). SenLearner was trained on cells at the extremes of the SDTMC module score. Cells in the top 10% of scores within each cell type that were also positive for at least one cyclin-dependent kinase inhibitor gene transcript were labelled senescent, and cells in the bottom 30% of each sample were labelled non-senescent (Fig. 3A; Methods). L1-regularized logistic regression models trained on these labels achieved high cross-validation performance across cell types (Supplementary Fig. 4C) and were used to assign senescence probabilities to all cells, including the majority that were initially unlabelled. SenLearner probabilities increased with age in a cell type-specific manner (Supplementary Fig. 4D). Cells with a probability above a cell type-specific threshold, defined as the lowest probability among senescent training cells, were classified as senescent. Using this threshold, the fraction of senescent cells increased with age in T cells (Spearman ρ = 0.66, P = 0.0040), hepatocytes (ρ = 0.64, P = 0.0040), cholangiocytes (ρ = 0.60, P = 0.011) and myeloid cells (ρ = 0.47, P = 0.048) (Fig. 3B, Supplementary Table 8). In hepatocytes, the parenchymal cells that make up most of the liver mass, the mean senescent fraction rose from 4.7% in young to 13.4% in very old mice (Wilcoxon rank-sum test, P = 0.0087), with no significant difference between young and old mice (P = 0.48; Fig. 3C, Supplementary Table 9). Cells classified as senescent showed expected senescence signatures, including higher interferon-associated SASP (Methods) and SenMayo^35^ scores than non-senescent cells in all four cell types (Wilcoxon rank-sum test, P < 0.0001; Fig. 3D).

**Fig. 3:**
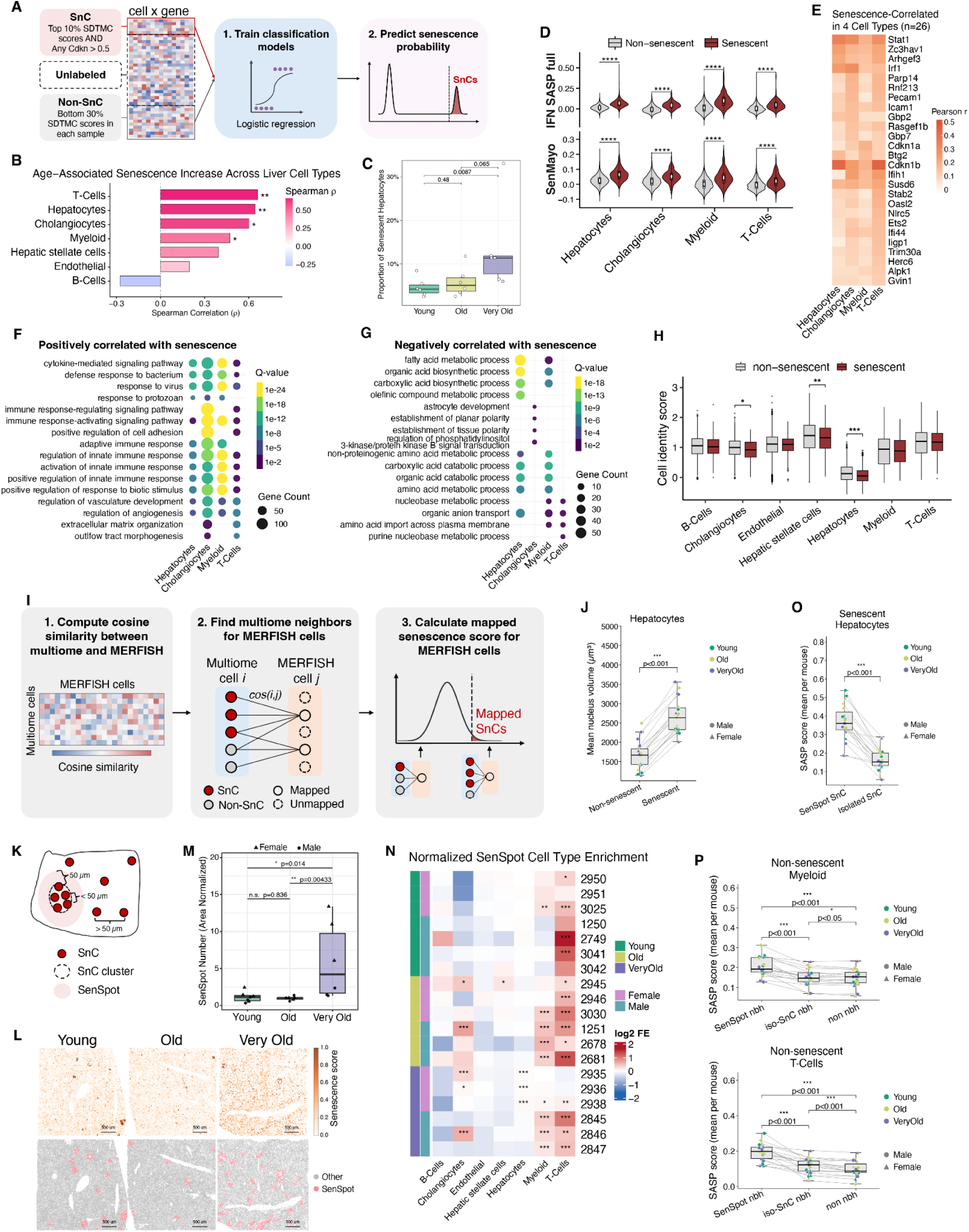
SenLearner identifies senescent cells across liver cell types and maps them in situ. **A**, The SenLearner framework. Within each cell type, cells in the top 10% of SDTMC module scores that were also positive for at least one cyclin-dependent kinase inhibitor (*Cdkn*) transcript were labelled as initial senescent cells (SnCs); within each sample, cells in the bottom 30% of scores were labelled as initial non-senescent cells (non-SnCs); the remainder were left unlabelled. An L1-regularized logistic regression model was trained on these labels to estimate a senescence probability for every cell (Methods). **B**, Spearman correlation between age and the proportion of senescent cells in each cell type (n = 18 mice). *P < 0.05, **P < 0.01. **C**, Proportion of senescent hepatocytes by age group. Points, individual mice (n = 6 per group); box, median and IQR; whiskers, 1.5 × IQR. P values, pairwise Wilcoxon rank-sum tests. **D**, Interferon-associated SASP (top) and SenMayo (bottom) module scores in senescent and non-senescent cells of each cell type. Violin, distribution across cells; box, median and IQR. ****P < 0.0001, Wilcoxon rank-sum test. **E**, Pearson correlation between gene expression and SenLearner probability for the 26 genes positively correlated with senescence in all four cell types (|r| > 0.05, adjusted P < 0.01). **F**, **G**, Gene ontology biological processes enriched among genes positively (**F**) and negatively (**G**) correlated with senescence probability in each cell type. Dot size, number of genes; color, adjusted q value. **H**, Cell identity scores in senescent and non-senescent cells of each cell type. Box, median and IQR; whiskers, 1.5 × IQR; points, outliers. *P < 0.05, **P < 0.01, ***P < 0.001, Wilcoxon rank-sum test, senescent versus non-senescent cells. I, Transfer of senescence labels from 10x Multiome to MERFISH. Each MERFISH cell was matched to its most similar Multiome cells by RNA cosine similarity and assigned a senescence score equal to the similarity-weighted fraction of those neighbours classified as senescent; cells in the upper tail were called senescent (Methods). **J**, Mean nuclear volume of senescent and non-senescent hepatocytes. Points, individual mice (n = 19 mice); lines connect paired measurements in each mouse. P value, paired Wilcoxon rank-sum test. **K**, Definition of SenSpots. Senescent cells lying within 50 µm of one another were grouped into clusters and SenSpots included these clustered SnCs and all surrounding cells within 50 µm; senescent cells with no senescent neighbours within 50 µm were classed as isolated (Methods). **L**, MERFISH sections from one mouse per age group, colored by mapped senescence score (top) and with SenSpots highlighted (bottom). Scale bars, 500 µm. **M**, Area-normalized SenSpot number by age group. Plotted as in **C**; point shape, sex. P values, pairwise Wilcoxon rank-sum tests; n.s., not significant. **N**, Enrichment of each cell type within SenSpots relative to the rest of the section, for 19 mice ordered by age group and sex. Color, log2 fold enrichment. *q < 0.05, **q < 0.01, ***q < 0.001, one-sided Fisher’s exact test with Benjamini-Hochberg FDR correction. **O**, SASP score of senescent hepatocytes inside SenSpots versus isolated senescent hepatocytes. Points, per-mouse means; lines connect paired measurements. Paired Wilcoxon rank-sum test. **P**, SASP score of non-senescent myeloid cells (top) and T cells (bottom) within a SenSpot neighbourhood (SenSpot nbh), adjacent to an isolated senescent cell (iso-SnC nbh), or neither (non nbh). Plotted and tested as in **O**.

To identify the gene programs underlying senescence in each cell type, we correlated gene expression with SenLearner probability. SenLearner probabilities correlated with 423–2,137 genes per cell type (Pearson |r| > 0.05, adjusted P < 0.01; positive and negative correlations combined), 89–97% of which were not in the SDTMC gene list (Supplementary Fig. 4E–F; Supplementary Table 10), indicating that SenLearner captures a broader senescence-associated program than the signature used to train it. Although the association with senescence was specific to individual cell types for most of these genes, 26 genes positively correlated with SenLearner probability in all four cell types (Fig. 3E). Besides the cell-cycle regulators *Cdkn1a, Cdkn1b and Btg2*, half of this shared set comprised interferon-response genes, including *Stat1, Irf1, Gbp2, Gbp7, Parp14*, *Ifih1, Oasl2, Nlrc5, Ifi44, Iigp1, Trim30a, Herc6 and Gvin1*. This suggests that interferon signalling is a common feature of senescence across hepatocytes, cholangiocytes, myeloid cells and T cells in the aging liver.

We next performed gene ontology enrichment analysis of senescence-correlated genes. Positively correlated genes were enriched for cytokine-mediated signalling, response to virus and innate and adaptive immune activation (Fig. 3F, Supplementary Table 11), whereas negatively correlated genes were enriched for metabolic processes, including fatty acid, organic acid and amino acid metabolism, most prominently in hepatocytes (Fig. 3G, Supplementary Table 11). Because metabolism is a principal function of hepatocytes, these results suggest that hepatocyte senescence is accompanied by a partial loss of metabolic programs and a gain of inflammatory ones. More broadly, senescent cells showed reduced expression of cell type identity genes in hepatocytes (P < 0.001), hepatic stellate cells (P < 0.01) and cholangiocytes (P < 0.05), but not in endothelial or immune cells (Wilcoxon rank-sum test; Fig. 3H).

Beyond these shared programs, each cell type showed distinct senescence-associated genes and pathways (Supplementary Fig. 4G, Supplementary Table 11). In senescent hepatocytes, positively correlated genes included the secreted SASP factors *Cxcl1, Igfbp2* and *Serpine1* (PAI-1) and were enriched for regulation of blood coagulation, haemostasis and wound healing, whereas negatively correlated genes included the metabolic and detoxification genes *Cyp2e1, Cyp2c38, Slco1b2* and *Gsta3* and were enriched for arachidonate, icosanoid and unsaturated fatty acid metabolism. Senescent cholangiocytes upregulated antigen presentation and pathways regulating natural killer cell-mediated cytotoxicity, consistent with increased immune recognition, while losing pathways that establish epithelial and planar polarity, consistent with disruption of biliary epithelial architecture. In myeloid cells, senescence was associated with expression of *Cdkn2a*, *Glb1*, *Il1b* and *Nfkbiz*, and with Toll-like receptor 2 signalling, production of CCL2 (MCP-1) and interleukin-18, and response to ionizing radiation, indicating an inflammatory, DNA damage-associated state. Senescent T cells were instead enriched for SMAD and BMP signalling and showed reduced organic acid transport and adenosine metabolism. TGF-β superfamily signalling has been linked to both cellular senescence^58^ and T cell dysfunction with age,^59^ although the role of BMP signalling in senescent T cells is less well characterized. Together, these results show that senescent cells in the aging liver share an inflammatory, interferon-associated program, while parenchymal and stromal cells also lose part of their cell type-specific functions.

We next examined the relationship between hepatocyte zonation and cellular senescence. Hepatocytes were assigned to zonation bins based on the expression of pericentral (PC) and periportal (PP) marker genes (Methods, Supplementary Fig. 5A–B, Supplementary Table 12). Although zonation scores showed no consistent age-associated changes (Supplementary Fig. 5C–D), both senescence probability and the proportion of SnCs decreased from PP to PC zones (Supplementary Fig. 5E–F). Consistently, the center of mass of the SnC zonal distribution was shifted toward PP zones relative to that of all hepatocytes, further supporting the enrichment of SnCs in periportal regions (Supplementary Fig. 5G).

To locate senescent cells in situ, we performed single-cell label transfer from the 10x Multiome to MERFISH spatial transcriptomics data (Fig. 3I). Each MERFISH cell was matched to its most similar 10x Multiome cells by RNA cosine similarity and assigned a senescence score equal to the similarity-weighted fraction of those neighbours classified as senescent; cells in the upper tail of this distribution were designated senescent (Methods). Mapped senescent cells showed higher senescence signatures such as SDTMC scores than non-senescent cells (Supplementary Fig. 6A). Senescent cells had larger nuclei than non-senescent cells within the same cell type, with senescent hepatocytes showing a 1.58-fold higher median mouse-level mean nuclear volume (2,633 versus 1,664 µm³; P < 0.001; Fig. 3J, Supplementary Fig. 6B, Supplementary Table 13), providing orthogonal support for their senescence labels, as nuclear enlargement is a feature of senescence.^22,26^ We next asked whether senescent cells were randomly distributed or clustered within the tissue. Neighborhood enrichment analysis revealed that senescent cells tended to colocalize (Supplementary Fig. 6C). We defined SenSpots as clusters of senescent cells lying within 50 µm of one another, together with the surrounding 50 µm, and senescent cells with no senescent neighbour within this distance as isolated (Fig. 3K, Methods). SenSpots were rare in young and old liver but more frequent in very old liver, with high variability between mice (Fig. 3L,M). Area-normalized SenSpot number was higher in very old mice than in young (P = 0.014) and old mice (P = 0.0043), with no difference between young and old mice (P = 0.84; Fig. 3M). SenSpots were enriched for myeloid cells and T cells in most mice (Fig. 3N), suggesting that immune cells accumulate around senescent cell niches.

Senescent hepatocytes within SenSpots had higher SASP scores than isolated senescent hepatocytes (P < 0.001; Fig. 3O), suggesting that their inflammatory output might be amplified by the senescent microenvironment. This extended to neighboring non-senescent cells: myeloid cells and T cells near SenSpots had the highest SASP scores, those near isolated senescent cells intermediate scores and those in senescence-free neighborhoods the lowest (P < 0.05 for all pairwise comparisons; Fig. 3P). Neighborhood inflammation thus increases with local senescent cell density, with SenSpots marking inflammatory niches that extend to surrounding immune cells. Together, these analyses identify age-associated senescence in hepatocytes, cholangiocytes, myeloid cells and T cells, defined by a shared interferon-response program and, in hepatocytes, loss of metabolic identity. In situ, senescent cells form immune-enriched SenSpots in very old liver, which are associated with heightened inflammation in neighboring cells.

### Gene regulatory programs associated with cellular senescence

To uncover the regulatory programs underlying these senescence-associated gene signatures, we identified putative enhancers that met two criteria: accessibility correlated with SenLearner probability across cells, and prediction as an enhancer of a target gene by the ABC model in the same cell type (Methods). This identified 5,102 senescence-associated putative enhancers, of which 4,567 gained and 535 lost accessibility with increasing SenLearner probability (|ρ| > 0.02, adjusted P < 0.05; Fig. 4A, Supplementary Table 14). These enhancers were largely cell type-specific, with 95% senescence-associated in only one cell type, indicating that senescence heterogeneity extends to the regulatory landscape. Across cell types, target genes of enhancers that gained accessibility (SnC-Up) showed higher expression correlations with senescence probability than all predicted target genes, whereas targets of enhancers that lost accessibility (SnC-Down) showed lower correlations (Wilcoxon rank-sum test, Benjamini–Hochberg-adjusted P < 0.01; Fig. 4B, Supplementary Table 15). For example, two interferon-induced guanylate-binding protein genes were predicted targets of SnC-Up enhancers: *Gbp3* in hepatocytes (r = 0.092, adjusted P = 9.5 × 10⁻¹⁶⁸) and *Gbp2* in myeloid cells (r = 0.186, adjusted P = 4.4 × 10⁻⁸⁷), consistent with the shared interferon program identified above (Fig. 4C). *Gbp3* expression was 4-fold higher in senescent than non-senescent hepatocytes (adjusted P = 1.3 × 10⁻⁹⁵). This consistent direction between enhancer accessibility and target gene expression supports a role for these putative enhancers in the senescence transcriptional program.

**Fig. 4:**
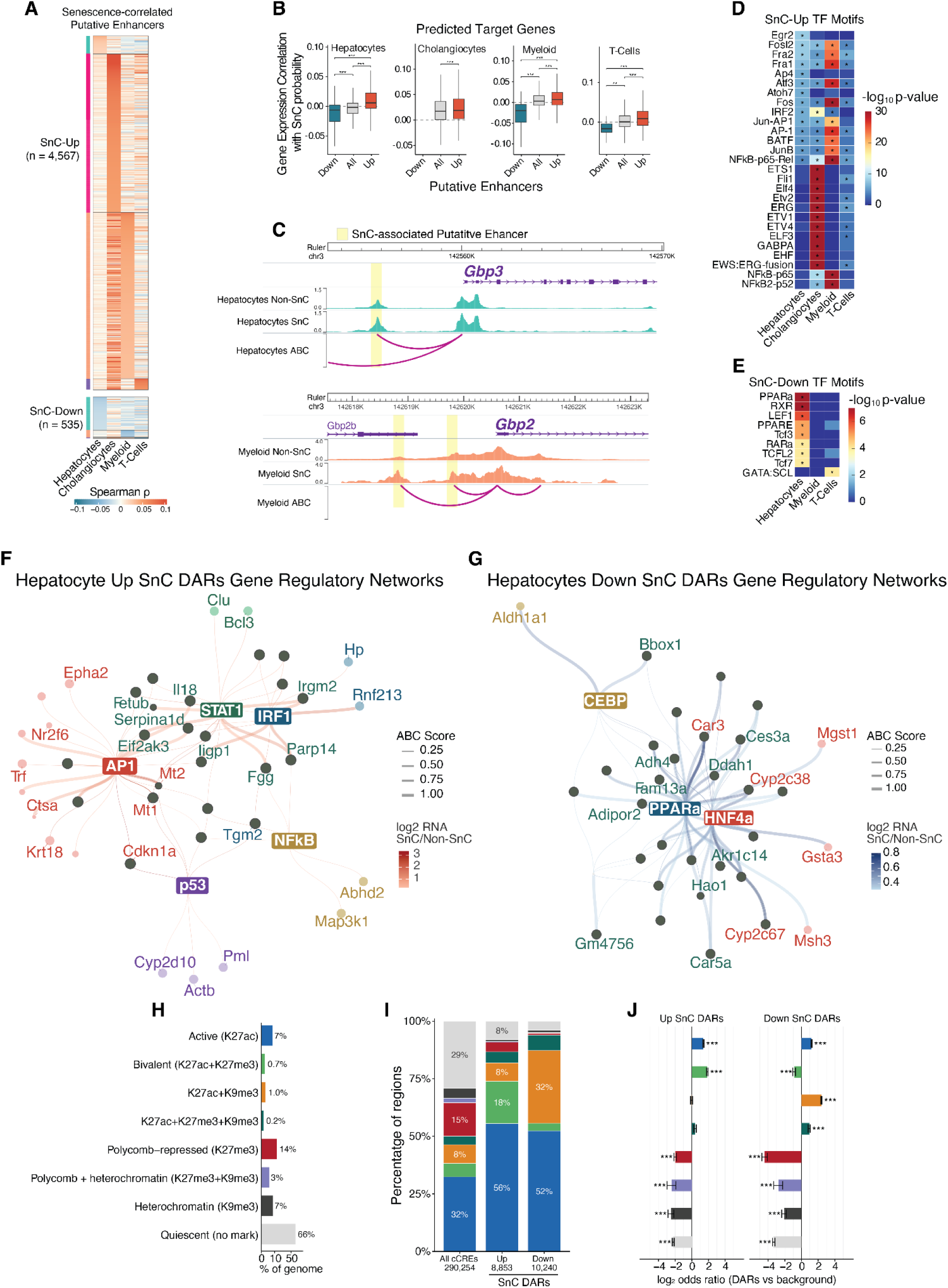
Gene regulatory programs associated with cellular senescence. **A**, Spearman correlation between the accessibility of ABC-predicted putative enhancers and SenLearner probability in each cell type, for the 4,567 enhancers that gained (SnC-Up) and 535 that lost (SnC-Down) accessibility with increasing probability (|ρ| > 0.02, adjusted P < 0.05). **B**, Correlation between target gene expression and SenLearner probability for genes predicted by ABC to be regulated by SnC-Down (Down), SnC-Up (Up) or all (All) putative enhancers, in each cell type. Box, median and IQR; whiskers, 1.5 × IQR. **P < 0.01, ***P < 0.001, Wilcoxon rank-sum test with Benjamini-Hochberg correction. Cholangiocytes had too few SnC-Down enhancers to test. **C**, Aggregate ATAC-seq signal in senescent and non-senescent cells at *Gbp3* in hepatocytes (top) and *Gbp2* in myeloid cells (bottom). Yellow shading, SnC-associated putative enhancers; arcs, ABC-predicted enhancer–gene links. **D**, **E**, Transcription factor motifs enriched in SnC-Up (**D**) and SnC-Down (**E**) putative enhancers in each cell type (HOMER known motifs). Color, −log10(P); asterisks, q < 0.05. Cell types with too few SnC-Down enhancers are omitted from **E**. **F**, **G**, Gene regulatory networks linking transcription factor motifs within differentially accessible regions (DARs) to ABC-predicted target genes upregulated (**F**) or downregulated (**G**) in senescent hepatocytes. Edge width, ABC score; node color, log2 fold change in expression between senescent and non-senescent hepatocytes. **H**, Fraction of the hepatocyte genome assigned to each of eight ChromHMM states derived from H3K27ac, H3K27me3 and H3K9me3 Paired-Tag data (200-bp bins; Methods). **I**, Chromatin state composition of all hepatocyte cCREs (n = 290,254) and of DARs that gained (n = 8,853) or lost (n = 10,240) accessibility in senescent hepatocytes. Colors as in **H**. **J**, Enrichment of each chromatin state among gained (left) and lost (right) DARs relative to all cCREs. Bars, log2 odds ratio; error bars, 95% confidence interval. ***adjusted P < 0.001, Fisher’s exact test with Benjamini-Hochberg correction. Colors as in **H**.

We next performed motif enrichment analysis on these enhancers in each cell type. SnC-Up enhancers were enriched for AP-1 family motifs (FOSL2, FRA1, FRA2, ATF3, JUNB and BATF) in hepatocytes, myeloid cells and T cells, for ETS family motifs (ETS1, FLI1, ERG and ELF3) and IRF2 in cholangiocytes, and for NF-κB motifs in myeloid cells (Fig. 4D, Supplementary Table 16). By contrast, SnC-Down enhancers in hepatocytes were enriched for nuclear receptor motifs, including PPARα, RXR and RARα, and for TCF/LEF motifs (Fig. 4E, Supplementary Table 16), factors that regulate hepatic lipid metabolism and zonated gene expression.^60^

To link these factors to their target genes in hepatocytes, we compared chromatin accessibility between senescent and non-senescent hepatocytes and identified 8,853 gained and 10,240 lost differentially accessible regions (DARs) (Supplementary Table 17). Gained DARs were enriched for motifs of the AP-1, IRF, NF-κB, STAT and p53 families, whereas lost DARs were enriched for motifs of PPARα, HNF4α and C/EBP, core regulators of hepatocyte identity and metabolic function. We then constructed gene regulatory networks connecting these motifs within DARs to differentially expressed genes predicted by ABC (Methods; Fig. 4F,G, Supplementary Table 18). The upregulated network recovered the canonical induction of *Cdkn1a* by p53 (Fig. 4F). AP-1 was the most connected factor, targeting the stress-responsive metallothioneins *Mt1* and *Mt2*, the receptor tyrosine kinase *Epha2* and the lysosomal carboxypeptidase *Ctsa*. Cathepsin A stabilizes lysosomal β-galactosidase (GLB1), the enzyme whose activity underlies the senescence-associated β-galactosidase stain,^61^ and higher *Ctsa* expression is consistent with the lysosomal expansion characteristic of senescent cells.^62^ STAT1 and IRF1 motifs were connected to interferon-response genes, including *Parp14*, *Irgm2* and *Iigp1*, and to the inflammatory cytokine *Il18*, whereas NF-κB motifs were connected to the kinase *Map3k1* and the hydrolase *Abhd2*. Several targets were shared between factors, suggesting that inflammatory, interferon and stress-response programs converge on a common set of senescence genes.

Transcription factors HNF4α and PPARα were predicted to regulate multiple downregulated genes (Fig. 4G). HNF4α is a master regulator of hepatocyte differentiation, and its loss is associated with hepatocyte dedifferentiation in chronic liver disease,^63^ whereas PPARα controls hepatic fatty acid oxidation and lipid metabolism.^60^ Consistent with these roles, both factors targeted genes of lipid, xenobiotic and drug metabolism, including the cytochrome P450 epoxygenases *Cyp2c38* and *Cyp2c67*, the glutathione S-transferases *Gsta3* and *Mgst1*, the peroxisomal oxidase *Hao1* and the adiponectin receptor *Adipor2*, while C/EBP targeted the retinoid-metabolizing enzyme *Aldh1a1*. Reduced accessibility at motifs for these factors suggests a regulatory basis for two features of senescent hepatocytes observed in Fig. 3: the loss of metabolic programs, including the arachidonate and icosanoid metabolism mediated by CYP2C enzymes (Fig. 3G), and the reduced expression of hepatocyte identity genes (Fig. 3H). Together, these networks indicate that hepatocyte senescence is accompanied by a gain of inflammatory, interferon and stress-response regulatory programs and a loss of programs that maintain identity and metabolic function.

Using H3K27ac, H3K27me3 and H3K9me3 profiles from Paired-Tag, we segmented the hepatocyte epigenome into eight chromatin states (Fig. 4H, Supplementary Table 19) and checked their overlap with DARs that gain accessibility (Up SnC) or lose accessibility (Down SnC) in senescence. Both Up SnC and Down SnC DARs were enriched in active chromatin marked by H3K27ac alone and depleted from repressed and chromatin states that lacked these marks (quiescent states) (Fisher’s exact test, Benjamini–Hochberg-adjusted P < 0.001; Fig. 4I,J). Up SnC DARs were selectively enriched in bivalent regions co-marked by H3K27ac and H3K27me3 (18.4% of gained DARs versus 5.9% of cCREs; odds ratio 3.6), which were depleted among Down SnC DARs. These elements may be held in a poised state in hepatocytes and become accessible upon senescence, consistent with the relief of Polycomb-mediated repression of inflammatory and stress-response genes during senescence.^64^

### Hepatocyte senotypes exhibit distinct gene regulatory programs, spatial organization, and zone-specificity

The heterogeneity of senescence motivated us to investigate whether senescent hepatocytes could be resolved into distinct states, or senotypes.^65^ To test this, we asked whether the senescence program itself is uniform across senescent hepatocytes or if these programs resolved into distinct co-expression modules. Genes significantly upregulated in senescent relative to non-senescent hepatocytes (adjusted P < 0.01, detected in more than 10% of cells) across the 6,342 senescent hepatocytes resolved four distinct co-expression modules, which we designate hepatocyte senotypes S1–S4 (Fig. 5A; Methods). These modules comprised 527 genes with significant module membership. Motif enrichment analysis of putative enhancers of these senotype genes revealed distinct transcription factor motifs enriched for each senotype (Fig. 5B). Specifically, senotype 1 (S1) genes (*Itih1*, *Serpina1a*, *Vtn*) were associated with hepatocyte secretory functions, with enrichment of SP2-binding motifs in their regulatory peaks. Senotype 2 (S2), characterized by a prominent interferon response signature including *Stat1*, *Ifih1*, *Pml* and *B2m*, was potentially regulated by STAT1-IRF programs. Senotype 3 (S3) was enriched for stress-response genes, including *Cdkn1a*, *Trp53inp1* and *Serpine1*, with putative enhancers showing enrichment of transcription factor motifs from the AP-1 family, including JUNB and FOS. In contrast, senotype 4 (S4) exhibited an inflammatory SASP (*Nfkb1*, *Cxcl1*, *Icam1*, *Stat3*), potentially driven by NF-κB-mediated transcriptional regulation. Interestingly, senotype 4 genes were also enriched for wound healing pathways, suggesting a potential involvement of this inflammatory senotype in tissue remodeling and repair in the aging liver (Fig. 5C).^66^

**Fig. 5:**
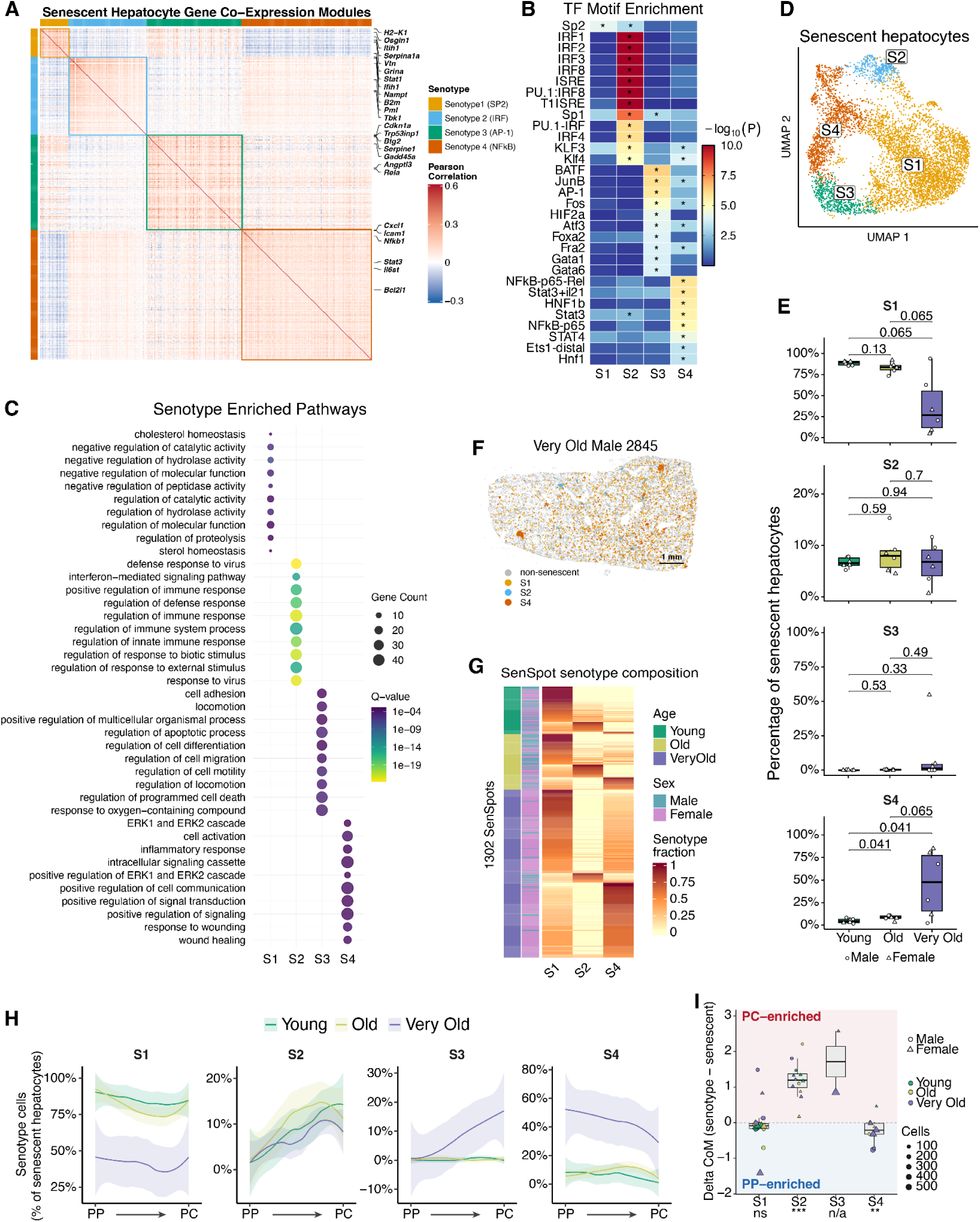
Hepatocyte senotypes have distinct regulatory programs, spatial organization and zonation. **A**, Pearson correlation between the expression of 527 senescence-associated genes across 6,342 senescent hepatocytes. Four co-expression modules, termed senotypes S1–S4, were identified by hierarchical clustering with Ward’s D2 linkage on the correlation distance (1 − r) and are outlined; selected genes are labelled. **B**, Transcription factor motifs enriched in ABC-predicted regulatory peaks linked to the genes of each senotype (HOMER known motifs). Color, −log10(P); asterisks, q < 0.05. **C**, Gene ontology biological processes enriched among the genes of each senotype. Dot size, number of genes; color, adjusted q value. **D**, UMAP of senescent hepatocytes clustered on senotype genes, colored by assigned senotype (n = 6,342 cells). **E**, Percentage of senescent hepatocytes assigned to each senotype, by age group. Points, individual mice (n = 6 per group); point shape, sex; box, median and IQR; whiskers, 1.5 × IQR. Wilcoxon rank-sum test; all P values shown. **F**, Senotype cells mapped onto a MERFISH section from a representative very old male mouse; non-senescent cells in gray. Scale bar, 1 mm. **G**, Senotype composition of 1,302 SenSpots. Each row is one SenSpot, annotated by the age group and sex of its mouse; color, fraction of senescent hepatocytes in that SenSpot belonging to each senotype. **H**, Percentage of senescent hepatocytes assigned to each senotype across 10 bins spanning the periportal (PP) to pericentral (PC) axis, by age group. Line, fitted mean across mice; shading, 95% confidence interval. **I**, Difference in the centre of mass of the zonal distribution between each senotype and all senescent hepatocytes; positive values indicate pericentral and negative values periportal enrichment. Points, individual mice; color, age group; shape, sex; size, number of cells. Only mice with at least 20 cells of that senotype are shown. **P < 0.01, ***P < 0.001, cell-count-weighted bootstrap test (Methods); n.s., not significant; n/a, not tested.

Having identified distinct transcriptional and regulatory programs associated with each senotype, we next identified cells belonging to these senotypes. By clustering senescent hepatocytes using senotype genes, we identified cell populations corresponding to the four senotypes (Fig. 5D, Supplementary Fig. 7A, Supplementary Table 20). These senotype cell populations showed differential distributions across age groups, with S4 cell proportion significantly increased with age (Fig. 5E). We next investigated how these senotype cells were spatially organized in liver tissues. S3 cells were excluded from this spatial analysis as they were predominantly from a single very old mouse (Fig. 5E). Although senotype genes were sparsely represented in the MERFISH panel, with no S1 genes included, classifiers based on the available SDTMC genes achieved high predictive performance in distinguishing senotype populations (Supplementary Fig. 7B), supporting their use for spatial mapping. The spatially mapped cells exhibited expression patterns consistent with their assigned senotypes (Supplementary Fig. 7C). Notably, the mapped senotype cells demonstrated distinct spatial organization patterns, where S1 cells were more dispersed whereas S2 and S4 cells were enriched in distinct SenSpots (Fig. 5F, Supplementary Fig. 7D).

To further characterize senotype-association with SenSpots, we quantified the proportion of each senotype within individual SenSpots, and revealed that different SenSpots were dominated by distinct senotypes (Fig. 5G). Therefore, we classified individual SenSpots according to their predominant senotype, with SenSpots lacking a single dominant senotype labeled as mixed SenSpots. The proportion of S4-dominant SenSpots increased significantly with age, whereas S1-dominant and mixed SenSpots decreased (Supplementary Fig. 7E), consistent with changes in the relative abundance of the corresponding senotype cell populations. Comparison of SenSpot sizes across senotypes revealed that S2-dominant SenSpots were significantly larger than other SenSpots (Supplementary Fig. 7F). Interestingly, the senotype cells in the SenSpots showed non-homogenous spatial distribution. S2 cells tended to be close to the SenSpot centroid while S1 cells were always close to the periphery of the SenSpots, even in the S1-dominant SenSpots (Supplementary Fig. 7G). Furthermore, S2-dominant SenSpots showed no pronounced immune cell enrichment whereas S4-dominant SenSpots exhibited enrichment of myeloid cells and T cells, suggesting senotype-specific associations with the immune microenvironment (Supplementary Fig. 7H). Given the spatial heterogeneity of hepatocyte senotypes, we next investigated whether their distributions were associated with hepatic zonation. Using the same periportal-to-pericentral zone bins defined above, we quantified the proportion of senotype cells across these bins. The proportion of S2 cells progressively increased from periportal to pericentral bins in every age group, whereas S4 cells, predominately in very old mice, exhibited the opposite trend, with highest proportions in the periportal bins (Fig. 5H, Supplementary Fig. 8A). Analysis of the center of mass of senotype cell proportion distributions revealed preferential pericentral localization of S2 cells and periportal localization of S4 cells (Fig. 5I). Consistently, S2 cells exhibited greater expression of pericentral markers, whereas S4 cells showed higher expression of periportal markers and lower expression of pericentral markers (Supplementary Fig. 8B).

Given the established role of Wnt/β-catenin signaling in maintaining pericentral hepatocyte identity,^67^ and the zone-dependent expression pattern of Wnt target genes (Supplementary Fig. 8C), we examined the expression of Wnt target genes across hepatocyte senotypes. Consistent with their distinct zonation patterns, S2 cells exhibited elevated Wnt target gene expression, whereas S4 cells showed reduced expression (Supplementary Fig. 8D). To further assess the relationship between senotype identity, hepatocyte zonation, and Wnt signaling, we correlated senotype signature scores with periportal, pericentral, and Wnt target gene scores across all hepatocytes. Despite being derived from non-overlapping gene sets, S4 signature scores were positively correlated with periportal scores and negatively correlated with both pericentral and Wnt target scores (Supplementary Fig. 8E–F), further supporting the association between S4 identity, periportal zonation and reduced Wnt-associated transcriptional activity. Collectively, these findings reveal four hepatocyte senotypes with distinct transcriptional and regulatory programs, spatial organization and lobular zonation, highlighting the heterogeneity of cellular senescence in the aging liver.

### Age-associated loss of DNA methylation at Constitutive Heterochromatin

DNA methylation is dynamic during aging^68^, therefore we examined the distribution of CG methylation (mCG) fractions across liver cells from our snm3C-seq data. Within myeloid cells, T cells, B cells, and endothelial cells, a minority of cells formed subclusters with globally lower methylation (Fig. 6A). We classified these cells as high, intermediate, or low mCG based on their mCG fractions (Fig. 6B). Low-mCG cells were more abundant in old and very old animals (Fig. 6C). To characterize these cells further, we transferred labels from snm3C-seq to the snATAC-seq data from the 10x multiome assay, which let us identify the corresponding low-mCG cells in a second assay. Low-mCG cells in the multiome data showed a significant association with age in myeloid and T cells (Fig. 6D), and trended toward higher abundance with age in all four cell types without reaching significance (Supplementary Fig. 9A).

**Fig. 6.**
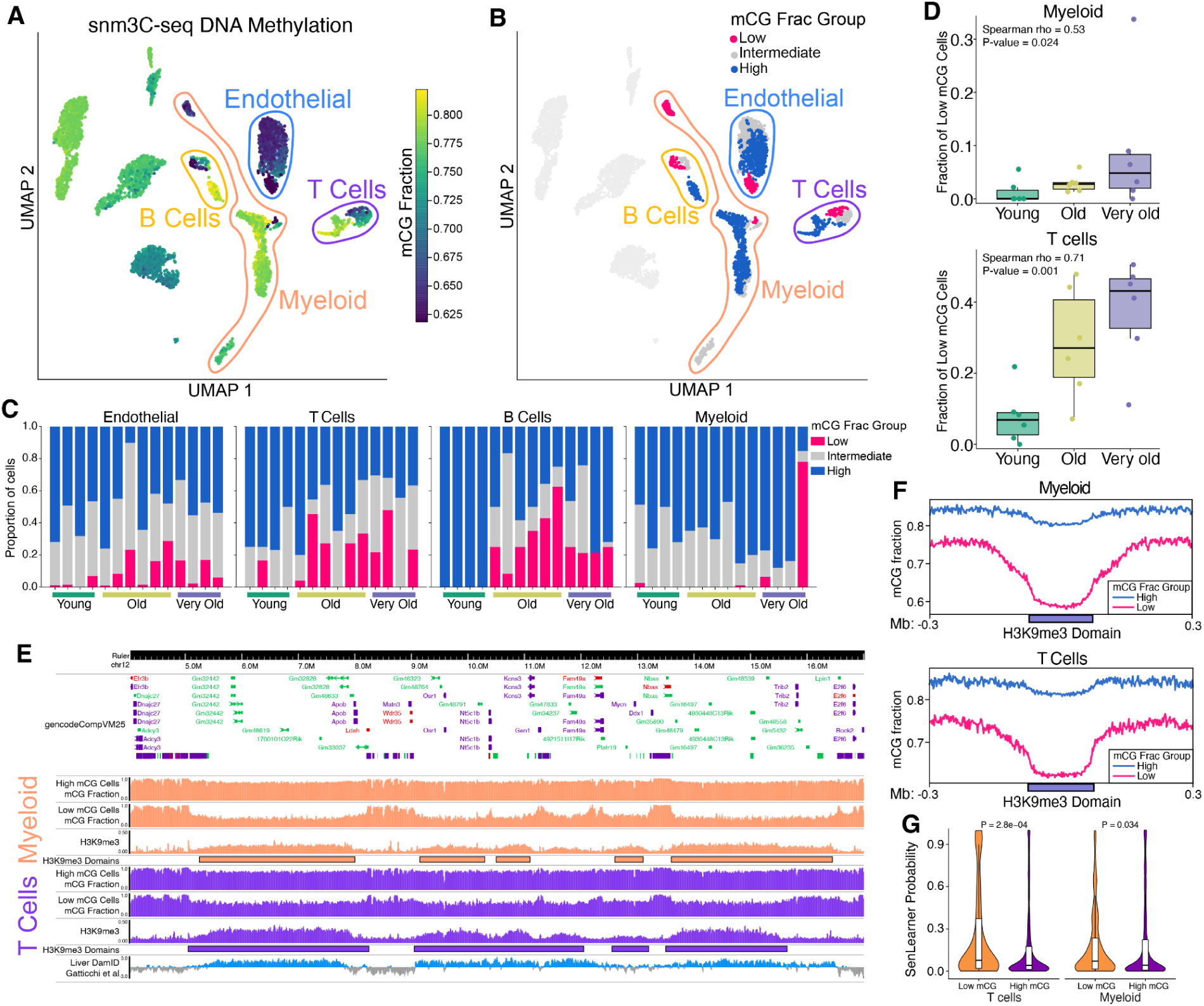
Age-associated loss of DNA methylation at domains of H3K9me3. **A**, UMAP of snm3C-seq nuclei based on mCG in 5-kb bins, coloured by global mCG fraction. Endothelial cells, B cells, T cells and myeloid cells are outlined. **B**, The embedding in A, with nuclei of the four outlined cell types coloured by mCG fraction group (low, intermediate or high; Methods); nuclei of other cell types are shown in light grey. **C**, Proportion of low, intermediate and high mCG nuclei in each mouse of the snm3C-seq cohort (n = 14 mice), by cell type. Each bar is one mouse, ordered by age group. **D**, Fraction of low-mCG nuclei per mouse in the 10x Multiome cohort (n = 18 mice, 6 per age group), calculated as low / (low + high) among nuclei assigned to a group by integration with snm3C-seq (Methods), for myeloid cells (top) and T cells (bottom). Box, median and IQR; whiskers, 1.5 × IQR; points, individual mice. Two-sided Spearman correlation with age in months. **E**, Genome browser view of chr12:4.2–16.8 Mb showing pseudobulk mCG fraction of high- and low-mCG nuclei, H3K9me3 signal and H3K9me3 domains called from Paired-Tag (Methods) for myeloid cells (orange) and T cells (purple). The bottom track shows lamina association from LaminB1 DamID in primary mouse hepatocytes^130^; this is a general reference for lamina association and is not cell type-matched. **F**, mCG fraction of high- and low-mCG nuclei across cell type-specific H3K9me3 domains, scaled to a common length, with 0.3-Mb flanking regions, in myeloid cells (n = 594 domains; top) and T cells (n = 552 domains; bottom). **G**, SenLearner senescence probability of 10x Multiome nuclei assigned to the low- or high-mCG group, in T cells and myeloid cells. Violin, density; box, median and IQR. P values, two-sided Wilcoxon rank-sum test on nuclei; a per-mouse paired analysis is shown in Supplementary Figure 9.

In the low-mCG cells, hypomethylation was concentrated in large, multi-megabase domains that coincided with high H3K9me3 signal, indicating loss of methylation at constitutive heterochromatin (Fig. 6E–F, Supplementary Fig. 9B). These domains overlapped lamina-associated domains^69^ (LADs; Fig. 6E, Supplementary Fig. 9C), which are typically enriched for heterochromatin^70^. Because reduced methylation at LADs has been reported in replicatively senescent human fibroblasts^71^, we compared SenLearner probabilities between groups. Low-mCG cells had significantly higher SenLearner probabilities than high-mCG cells (Fig. 6G), suggesting that the low-mCG cells with hypomethylation at H3K9me3 domains are consistent with a senescent-like state.

### p21-directed *Stat1* knockout alters inflammatory signaling in the aged liver

Our previous analyses identified multiple signaling pathways, including Stat1-IRF, NF-κB and AP-1, in senescence-associated regulatory programs, particularly in different hepatocyte senotypes (Fig. 4F, 5B). To further prioritize transcription factors for functional perturbation, we systematically evaluated their senescence-associated expression changes and co-expression with *Cdkn1a* in hepatocytes (Fig. 7A). Notably, *Stat1* was both upregulated during senescence and broadly expressed within the *Cdkn1a*+ cells, supporting its selection as a candidate regulator for genetic perturbation in senescent hepatocytes. We therefore used a p21 promoter-directed, Cre-dependent CRISPR-Cas9 system to target *Stat1* in p21-expressing cells of aged mice and examined the resulting transcriptional changes in the liver (Fig. 7B, Supplementary Fig. 10A). Sanger sequencing of genomic DNA from liver samples confirmed the *Stat1* knockout at the target locus (Supplementary Fig. 10B). Furthermore, the percentage of *Stat1*-expressing cells among p21+ cells was significantly decreased after the *Stat1* knockout (Stat1KO) compared to the control (Supplementary Fig. 10C–D), confirming the *Stat1* knockout in p21-directed cells. To assess the transcriptional consequences of p21-directed *Stat1* knockout, we performed bulk RNA sequencing of liver tissues from aged Stat1KO and control mice. Gene set enrichment analysis (GSEA)^72^ showed that immune-related programs, including immune response and response to interferon-β, were significantly downregulated following *Stat1* knockout (Fig. 7C), which were elevated in senescent cells (Fig. 3F). Conversely, pathways associated with catabolic processes were upregulated after *Stat1* knockout, which were suppressed in senescent cells (Fig. 3G), suggesting at least a partial reversal of senescence-associated transcriptional alterations. To further investigate the specific transcriptional effect of *Stat1* knockout in senescent cells, we focused on predicted Stat1 targets by the ABC model (Supplementary Table 5) that were upregulated in senescent hepatocytes and assessed their expression changes after p21-directed *Stat1* knockout (Fig. 7D). The majority of these genes exhibited reduced expression in Stat1KO livers compared with controls (Fig. 7D). Furthermore, the log2 fold changes of predicted STAT1 targets between SnCs and non-SnCs were negatively correlated with their log2 fold changes between Stat1KO and control livers (Fig. 7E), indicating that genes more strongly upregulated during senescence tended to exhibit greater downregulation following *Stat1* knockout. Consistently, gene set variation analysis (GSVA)^73^ showed that the enrichment score of predicted STAT1 targets was significantly reduced in Stat1KO livers compared with controls (Fig. 7F). The senotype 2 signature, characterized by elevated interferon signaling and STAT1-IRF regulatory activity, also showed a significant reduction following *Stat1* knockout (Fig. 7F). In contrast, the signatures of other hepatocyte senotypes, as well as gene sets comprising all senescence-upregulated or senescence-downregulated genes, showed no significant changes (Supplementary Fig. 10E–F). Together, these findings suggest that p21-directed *Stat1* knockout preferentially attenuates STAT1-associated and interferon-responsive senescence programs, and partially reverses senescence-associated metabolic alterations in the aged liver.

**Fig. 7:**
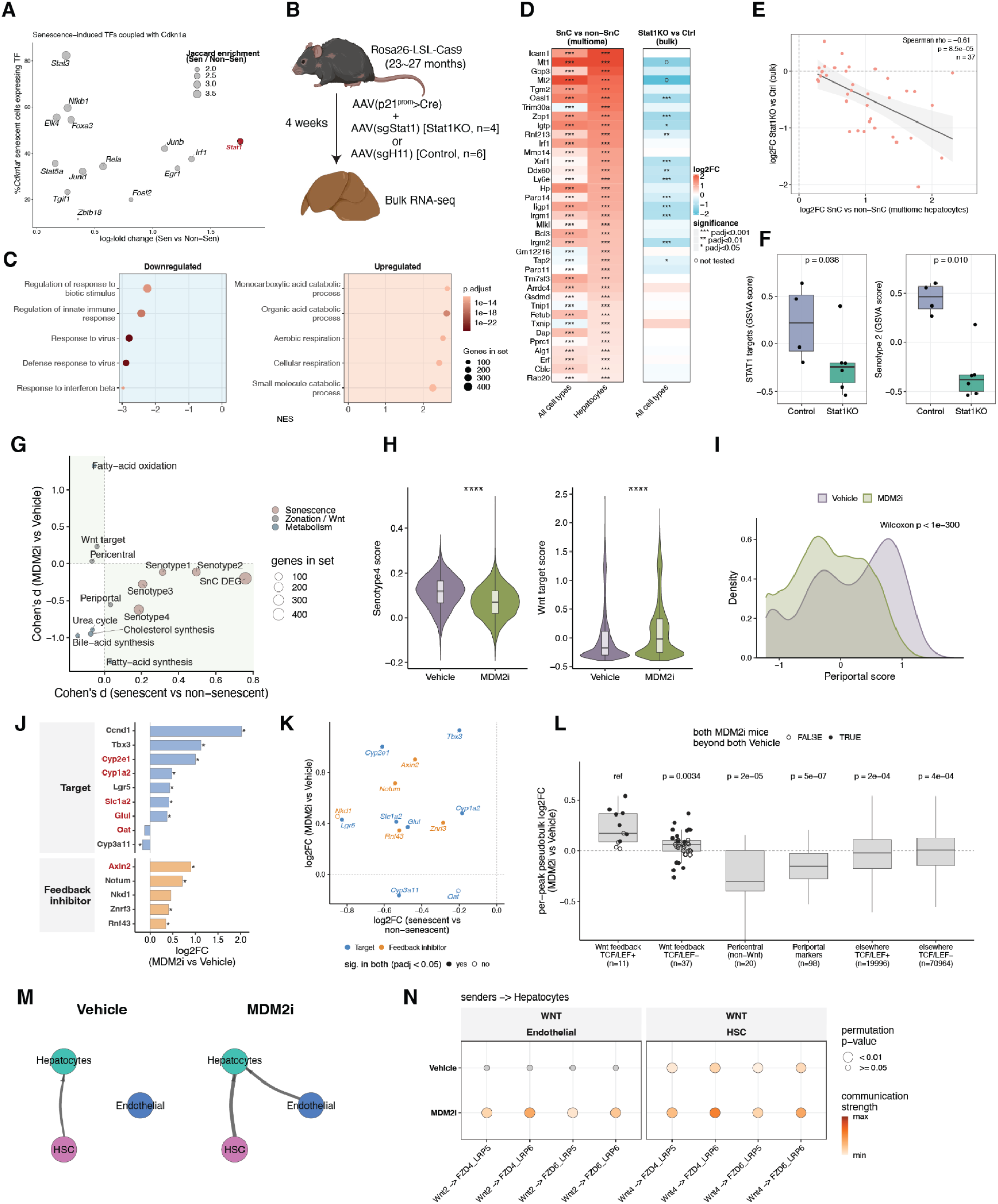
*Stat1* knockout and MDM2 inhibition act on different hepatocyte senotypes. **A**, Transcription factors induced in senescent hepatocytes, plotted by log2 fold change between senescent and non-senescent cells against the percentage of *Cdkn1a*-positive senescent cells expressing them. Point size, Jaccard enrichment of co-expression with *Cdkn1a*. *Stat1* is highlighted. **B**, Experimental design (BioRender). *Rosa26*-LSL-Cas9 mice aged 23–27 months received AAV expressing Cre from the *p21* promoter together with AAV-sgStat1 (Stat1KO, n = 4) or AAV-sgH11 (control, n = 6); liver was collected for bulk RNA sequencing after 4 weeks. **C**, Gene set enrichment analysis of Stat1KO versus control liver, showing the top 5 downregulated (left) and upregulated (right) gene sets. NES, normalized enrichment score; dot size, number of genes in the set; color, adjusted P value. **D**, Log2 fold change of ABC-predicted STAT1 target genes that were upregulated in senescent hepatocytes, in senescent versus non-senescent cells (10x Multiome; all cell types and hepatocytes alone) and in Stat1KO versus control liver (bulk RNA-seq). *adjusted P < 0.05, **adjusted P < 0.01, ***adjusted P < 0.001; open circles, not tested. **E**, Log2 fold change of predicted STAT1 targets in Stat1KO versus control liver against the same genes in senescent versus non-senescent hepatocytes. Line, linear fit; shading, 95% confidence interval; rho, Spearman correlation (n = 37 genes). **F**, Gene set variation analysis (GSVA) scores for predicted STAT1 targets (left) and the senotype 2 signature (right) in control and Stat1KO liver. Points, individual mice; box, median and IQR; whiskers, 1.5 × IQR. Wilcoxon rank-sum test. **G**, Effect size (Cohen’s *d*) of gene sets in senescent versus non-senescent hepatocytes against MDM2i- versus vehicle-treated hepatocytes. Color, gene set category; size, number of genes. Shaded quadrants contain sets that changed in opposite directions in the two comparisons. **H**, Senotype 4 (left) and Wnt target (right) module scores in hepatocytes from vehicle- and MDM2i-treated mice. Violin, distribution across nuclei; box, median and IQR. ****P < 0.0001, Wilcoxon rank-sum test. n = 2 mice per group. **I**, Distribution of periportal signature scores in hepatocytes from vehicle- and MDM2i-treated mice. Wilcoxon rank-sum test. **J**, Log2 fold change (MDM2i versus vehicle) of Wnt target genes and Wnt negative-feedback regulators in hepatocytes. *adjusted P < 0.05. Genes in red, pericentral marker genes. **K**, Log2 fold change of the genes in **J** in MDM2i- versus vehicle-treated hepatocytes against senescent versus non-senescent hepatocytes. Filled points, significant in both comparisons (adjusted P < 0.05). **L**, Per-peak pseudobulk log2 fold change (MDM2i versus vehicle) for ABC-predicted regulatory peaks of Wnt feedback inhibitor genes containing (TCF/LEF+) or lacking (TCF/LEF−) a TCF/LEF motif, alongside pericentral non-Wnt peaks, periportal marker peaks and all remaining peaks. Box, median and IQR; whiskers, 1.5 × IQR; n, peaks. P values are relative to the Wnt feedback TCF/LEF+ set (ref), Wilcoxon rank-sum test. Filled points, peaks at which both MDM2i mice exceeded both vehicle mice. **M**, CellChat-inferred Wnt signalling networks among hepatocytes, hepatic stellate cells (HSC) and endothelial cells in vehicle- and MDM2i-treated mice. Edge width, communication strength. **N**, Inferred ligand–receptor communication to hepatocytes from endothelial cells, and hepatic stellate cells, through canonical Wnt signalling. Dot size, permutation P value; color, communication strength.

### MDM2 inhibition suppresses senescence-associated signatures in aged hepatocytes

Previous work showed that p53 restrains the SASP.^8^ Pharmacological activation of p53 with the MDM2 inhibitor HDM201 enhances DNA repair, suppresses senescence-associated inflammation, and partially reverses aging-associated transcriptional changes in the liver^74,75^. To further characterize the effects of MDM2 inhibition (MDM2i) on hepatocyte senescence and senotypes, we performed single-nucleus multiome profiling of aged female mice treated with HDM201 (n = 2) or vehicle (n = 2). We first examined whether MDM2i rescued the transcriptional alterations associated with cellular senescence. Comparing pathway-level effect sizes between senescent and non-senescent hepatocytes with those between MDM2i-treated and vehicle-treated hepatocytes revealed broadly opposing patterns (Fig. 7G). These included senescence-associated differentially expressed genes, senotype 3 and senotype 4 signatures, periportal and pericentral zonation markers, Wnt target genes, and fatty acid oxidation pathways, suggesting that MDM2i reversed multiple senescence-associated transcriptional changes. Among senescence-associated signatures, the inflammatory senotype S4 exhibited a pronounced decrease following MDM2i treatment (Fig. 7H, Supplementary Fig. 10G–H). Given the association of S4 with Wnt target gene expression and hepatocyte zone scores (Supplementary Fig. 8E–F), we investigated the effects of MDM2i on hepatocyte Wnt signaling and zonation. MDM2i increased Wnt target gene score (Fig. 7H), accompanied by decreased periportal (Fig. 7I) and increased pericentral signature scores (Supplementary Fig. 10I). To determine whether these changes reflected shifts in hepatocyte zonal distribution or transcriptional alterations within zones, we examined individual pericentral and periportal marker gene expression across zonation bins. Periportal markers consistently exhibited reduced expression following MDM2i treatment, even when comparing hepatocytes within the same zone bins (Supplementary Fig. 10J), suggesting that MDM2i induced changes in hepatocyte zonation transcriptional programs beyond shifts in the hepatocytes distribution along the portal-central axis.

To characterize the Wnt response after MDM2i treatment in greater detail, we examined genes encoding different components of the Wnt signaling pathway. Whereas Wnt receptors and signal transducers exhibited heterogeneous transcriptional responses (Supplementary Fig. 10K), Wnt targets, and negative-feedback regulators showed more consistent increases after MDM2i treatment (Fig. 7J). Notably, among these 13 Wnt target and feedback inhibitor genes, 11 of them exhibited opposite expression changes between senescence and MDM2i treatment (Fig. 7K). Wnt negative-feedback regulators, such as *Axin2* and *Notum*, are transcriptionally induced by canonical Wnt/β-catenin signaling and subsequently act as negative regulators to finetune Wnt pathway activity.^76–78^ Thus, the engagement of negative-feedback mechanisms demonstrated increased Wnt pathway activation. We next investigated whether these transcriptional changes were accompanied by alterations in chromatin accessibility. Among regulatory peaks predicted by the ABC model to control Wnt feedback inhibitor genes (Supplementary Table 5), peaks containing TCF/LEF motifs exhibited greater accessibility increase after MDM2i treatment than those lacking these motifs (Fig. 7L). Finally, we used CellChat^79^ to infer potential changes in Wnt signaling among hepatic stellate cells, endothelial cells, and hepatocytes. Canonical Wnt signaling demonstrated increased inferred communication strength after MDM2i treatment (Fig. 7M), with consistently stronger inferred interactions across individual Wnt ligand-receptor pairs (Fig. 7N). Together, these findings indicate that MDM2 inhibition counteracts multiple senescence-associated transcriptional changes in aged hepatocytes, particularly the inflammatory senotype 4 signatures, while enhancing Wnt target gene expression.

## Discussion

Here, we present a single-cell multimodal atlas of the aging mouse liver integrating gene expression, chromatin accessibility, histone modifications, DNA methylation, three-dimensional genome architecture and spatial organization. This integrated view reveals two interconnected but distinct dimensions of liver aging. First, aging remodels the tissue broadly, shifting it towards an immune-enriched and inflammatory state while eroding the specialized functions that define individual cell types. Second, senescent cells accumulate predominantly in late life, forming spatially organized niches and adopting distinct regulatory and transcriptional states.

A prominent feature of liver aging was the loss of hepatocytes accompanied by an expansion of immune and stellate cell populations. This shift in cellular composition was accompanied by a striking convergence in inflammatory gene expression across otherwise distinct cell types: inflammatory programs were induced not only in immune cells but also in hepatocytes, stellate cells and endothelial cells. By contrast, genes downregulated with age were largely specific to individual cell types, reflecting the reduced capacity of their specialized functions rather than a uniform loss of transcriptional activity. Aging therefore drives two complementary transcriptional changes: a shared inflammatory response across the liver and a cell type-specific decline in differentiated identity. The loss of lipid transport programs in endothelial cells and vascular support programs in stellate cells is consistent with the sinusoidal dysfunction described in aged liver,^43,44^ indicating that age-related functional decline extends beyond hepatocytes to the cellular networks that sustain the hepatic microenvironment.

The relationship between epigenetic erosion and transcriptional dysfunction has been proposed as a central feature of aging,^54^ but much of the evidence in mammals has relied on bulk tissue measurements,^80^ leaving unresolved how epigenetic changes relate to transcriptional alterations within individual cells in vivo. By combining chromatin profiling and gene expression measurements in the same nuclei using Paired-Tag, we found that nuclei with lower histone modification signal exhibited greater transcriptional drift. Because these measurements were obtained from the same nucleus and compared within cell types, this relationship cannot be explained by age-related changes in cellular composition. In hepatocytes, this erosion was accompanied by extensive remodeling of genome organization, including increased trans-chromosomal contacts, reduced intra-TAD interactions and a strongly asymmetric loss of H3K9me3, with more than 8,000 domains lost and only a few hundred gained. These losses occurred at promoters of DNA damage response, autophagy and senescence genes, which were correspondingly induced with age. Depletion of H3K9me3 can induce cellular senescence and premature aging in mice,^57^ and we therefore speculate that the erosion observed here does not simply accompany cellular aging but creates a permissive state for the acquisition of senescence-associated programs. Whether this erosion precedes senescence or accumulates as a consequence of cellular stress remains to be determined.

A further challenge in studying senescence in vivo is its substantial cellular and molecular heterogeneity. No single marker reliably identifies senescent cells across tissues and cell types, and the limited detection of *Cdkn2a* transcripts by single-cell RNA sequencing further complicates their identification, motivating recommendations from the SenNet consortium to use cell type-informed signatures rather than individual genes.^26^ Consistent with this principle, we developed SenLearner, a machine-learning framework that learns cell type-specific senescence classifiers from the SDTMC genes. Most genes recovered by SenLearner were not in the original list, highlighting the extent to which senescence-associated programs differ between cell types, while the classifiers also identified shared inflammatory and interferon-associated components across cell types, consistent with the SenMayo and IFN SASP signatures. These cell type-specific programs may have distinct consequences for liver function. Myeloid cells acquired inflammatory and damage-associated features, whereas senescent T cells showed altered TGF-β superfamily signalling, which is associated with impaired T cell function in aging.^59^ Because senescent cells are removed in part through immune surveillance,^11^ senescence within the immune compartment itself may compromise the clearance of senescent cells elsewhere in the tissue. In hepatocytes, the emergence of wound-healing programs alongside the loss of metabolic functions suggests a shift from differentiated hepatic activity towards stress-responsive tissue remodeling.

This transition was evident in the hepatocyte regulatory landscape. Senescent hepatocytes exhibited reduced accessibility at elements containing HNF4α and PPARα motifs, accompanied by decreased expression of cytochrome P450 enzymes and genes involved in detoxification and fatty acid metabolism. HNF4α is a central regulator of hepatocyte differentiation, and its loss is associated with progressive chronic liver disease and inflammatory cytokine signaling.^63^ Conversely, senescent hepatocytes gained accessibility at elements containing AP-1 and NF-κB motifs, central regulators of SASP gene expression, providing a potential link between the loss of hepatocyte identity and the acquisition of inflammatory programs; a recent study of human and mouse liver similarly reported increased AP-1 and NF-κB motif accessibility in senescent cells.^22^ Notably, these changes were not distributed uniformly across the epigenome but concentrated within specific chromatin states: gained elements were enriched in bivalent regions, whereas 31.6% of lost elements fell within an H3K27ac and H3K9me3 co-marked state covering only 1% of the genome. Rather than undergoing indiscriminate remodeling, senescent hepatocytes activate and repress regulatory elements according to their pre-existing epigenetic context.

Senescent cells were also not randomly distributed throughout the aging liver. They formed spatially clustered regions (SenSpots) that increased in abundance in very old animals and were enriched for myeloid cells and T cells. Within these regions, senescent hepatocytes exhibited higher SASP scores than isolated senescent hepatocytes, as did neighboring myeloid and T cells that were not themselves classified as senescent. This organization is consistent with paracrine propagation of senescence^81^ and with immune surveillance of senescent hepatocytes;^11^ these mechanisms are not mutually exclusive, and our cross-sectional data cannot distinguish them. Within these niches, senescent hepatocytes occupied distinct molecular states associated with their position in the lobule. The inflammatory S4 senotype expanded with age to approximately half of all senescent hepatocytes and was enriched periportally, whereas the interferon-associated S2 senotype was enriched pericentrally with elevated Wnt target expression. Given the metabolic and signaling gradients across the lobule, including oxygen availability and Wnt signaling,^17^ the zonal environment may shape the regulatory program a hepatocyte adopts as it becomes senescent. Senescent hepatocytes have also been reported to cluster and to show periportal enrichment in the human liver,^22^ suggesting that this spatial organization is a general feature of hepatic senescence.

The functional significance of this heterogeneity was evident in the response to intervention. p21-directed *Stat1* knockout preferentially reduced the S2 signature and the expression of predicted STAT1 targets, with the reduction scaling with how strongly each target was induced during senescence (ρ = −0.61), without substantially affecting other senotypes. MDM2 inhibition instead reduced the S4 signature and increased Wnt target expression, accessibility at TCF/LEF motif-containing elements and predicted Wnt signaling from stellate and endothelial cells. Senescent states are therefore not uniformly susceptible to intervention, and targeting one regulatory program may leave others unaffected. Strategies aimed at eliminating or modifying senescent cells may benefit from considering not only the abundance of senescent cells but also their cell type, regulatory state and spatial distribution. Further work will be needed to establish causality, including validation with protein-level senescence markers, perturbations restricted to senescent cells and larger intervention cohorts.

In summary, our multimodal atlas reveals that liver aging involves widespread loss of cellular specialization and epigenetic organization alongside the emergence of spatially organized and molecularly heterogeneous senescence. These processes are interconnected but distinct: inflammatory and functional remodeling occurs broadly across cell types, whereas senescence emerges in specific populations and local environments. By combining multimodal profiling, machine-learning-based senescence identification and spatial analysis, this study shows that understanding tissue aging requires distinguishing the pervasive changes that affect most cells from the specialized cellular states that accumulate within the aging microenvironment.

## Methods

### Mice and housing

All animal experiments were approved by the Institutional Animal Care and Use Committee of Sanford Burnham Prebys Medical Discovery Institute (AUF 22-010, 22-040, 23-026, 25-053 and 26-008) and performed in accordance with NIH guidelines. C57BL/6JN mice were obtained from the National Institute on Aging Aged Rodent Colony and housed under specific pathogen-free conditions at 21–24 °C and 30–70% humidity on a 12 h light/dark cycle with ad libitum access to food (Teklad 2018) and water. Young (3–5 months), old (21–24 months) and very old (29–31 months) mice were used as indicated in the figure legends. Mice were allowed to acclimate to the facility for 2 weeks on arrival before any procedure.

### Nuclei isolation and sorting for 10x Multiome and Paired-Tag

Nuclei were isolated from pre-sectioned, flash-frozen tissue stored at −80 °C. All buffers were prepared fresh, pre-chilled on ice, and supplemented with 1 U/µL RNase inhibitor (Promega,

PAN2515) for 10x Multiome experiments; 0.5 U/µL SUPERaseIN (ThermoFisher Scientific, AM2696) and 0.5 U/µL RNAseOUT (ThermoFisher Scientific, 10777019), and all centrifugation steps were performed at 4 °C in a swinging-bucket rotor with low acceleration and deceleration settings. Frozen tissue was transferred on dry ice to gentleMACS M tubes containing ice-cold MACS buffer (5 mM CaCl₂, 3 mM magnesium acetate, 2 mM EDTA, 10 mM Tris-HCl pH 8, 0.6 mM DTT, and 1× protease inhibitor (Pierce A32965), thawed on ice for approximately 60 s, and homogenized on a gentleMACS Octo Dissociator using the Protein_01_01 program (Miltenyi Biotec). The homogenate was filtered through a 30 µm CellTrics filter, and the M tube was rinsed with 1 mL MACS buffer, which was passed through the same filter. Nuclei were pelleted at 500 × g for 5 min, resuspended in 500 µL sort buffer (1% BSA and 1× protease inhibitor in PBS) containing 2 µm 7-AAD, and incubated on ice for 15 min. After a second centrifugation (500 × g, 5 min), nuclei were resuspended in 500 µL sort buffer and 150,000 7-AAD-positive nuclei per sample were sorted by fluorescence-activated sorting with Sony, SH800S into 100 µL collection buffer (5% BSA in PBS).

### 10x Multiome library generation

Following FANS, final collection volumes were determined and 5X Permeabilization Buffer (50mM Tris-HCl pH 7.4, 50mM NaCl, 15mM MgCl2, 0.05% Tween-20, 0.05% IGEPAL, 0.005% Digitonin, 5% fatty acid-free BSA in PBS, 5mM DTT, 1U/μL Recombinant RNase inhibitor, 5X Protease Inhibitor) was added to achieve a 1X solution. Nuclei were incubated on ice for 1 minute, then centrifuged (500 rcf, 5min at 4C) in a swinging bucket centrifuge. Supernatant was discarded and 650uL of Wash Buffer (10mM Tris-HCl pH 7.4, 10mM NaCl, 3mM MgCl2, 0.1%.Tween-20, 1% fatty acid-free BSA in PBS, 1mM DTT, 1U/μL Recombinant RNase inhibitor, 1X Protease Inhibitor) was added without disturbing the pellet followed by centrifuging (500 rcf, 5 min at 4°C) in a swinging bucket centrifuge. Supernatant was removed, and the pellet was resuspended in 7uL of 1X Nuclei Buffer (Nuclei Buffer (10x Genomics), 1mM DTT, 1 U/μL Recombinant RNase inhibitor). 1μL was used for counting on a hemocytometer after staining with Trypan Blue (Invitrogen, T10282). 16-20k nuclei were used for tagmentation reaction and 10x Genomics controller loading. Libraries were then generated following the manufacturer’s recommended protocol (https://www.10xgenomics.com/support/single-cell-multiome-atac-plus-gene-expression). 10x multiome ATAC-seq and RNA-seq libraries were paired-end sequenced on a NovaSeq X Plus to a target depth of ∼50,000 reads per cell for each modality.

### 10x Multiome data processing, clustering and annotation

Sequencing reads were aligned to the mouse reference genome (mm10) and processed with Cell Ranger ARC (v2.0.2) to generate gene expression count matrices and ATAC fragment files for each sample. Ambient RNA was removed with CellBender (v0.3.0)^82^ using default parameters, and putative doublets were identified in each sample with DoubletFinder (2.0.4)^83^ using default parameters. Samples were merged in Seurat (v5.3.0)^84^ and Signac (v1.14.0),^85^ and TSS enrichment scores were computed with Signac^85^ using default parameters. Nuclei were retained if they had more than 500 ATAC fragments, more than 200 detected genes, a TSS enrichment score greater than 1 and less than 10% mitochondrial reads.

Gene expression counts were normalized with SCTransform,^86,87^ and principal component analysis was performed on variable genes.To account for sample-associated batch effects, principal component analysis (PCA) embeddings were integrated across samples using Harmony^88^ through the IntegrateLayers function in Seurat (v5.3.0). The first 30 Harmony-corrected components were subsequently used to construct a shared nearest-neighbor (SNN) graph for downstream clustering and analysis. To remove residual multiplets, nuclei were first clustered at high resolution (Leiden, resolution 5), and the 12 clusters in which more than 30% of nuclei were classified as doublets by DoubletFinder were excluded, together with all remaining nuclei classified as doublets. The remaining nuclei were then reclustered (Leiden, resolution 0.1) and visualized by UMAP. Clusters were annotated as hepatocytes, cholangiocytes, hepatic stellate cells, endothelial cells, myeloid cells, T cells and B cells on the basis of canonical marker gene expression (Supplementary Fig. 2A). After quality control, 154,645 nuclei were retained for downstream analyses.

### Cell type composition analysis

For each mouse, the proportion of each cell type was calculated as the number of nuclei assigned to that cell type divided by the total number of nuclei passing quality control in that sample (10x Multiome; n = 6 mice per age group). Proportions were compared between young and very old mice for each cell type using a two-sided Wilcoxon rank-sum test with normal approximation.

### Age-associated differential expression and accessibility

For each cell type, raw counts were summed across nuclei within each mouse to generate pseudobulk profiles, of genes for RNA and of cCREs for ATAC. Profiles were analysed with edgeR (v4.4.2)³⁹: features were retained using filterByExpr, libraries were normalized by the trimmed mean of M-values (TMM) method, and dispersions were estimated with estimateDisp. A negative binomial generalized linear model with age group as the sole covariate was fitted with glmFit, and pairwise contrasts between age groups (old versus young, very old versus young, very old versus old) were tested with glmLRT. Genes with a Benjamini–Hochberg FDR < 0.05 were considered differentially expressed, and cCREs with an FDR < 0.1 differentially accessible.

### Cell type-specific enhancers and their target genes

For each cell type, non-promoter enhancers predicted by the ABC model⁴⁶ (ABC score > 0.02) were identified as described above. Pseudobulk accessibility of each enhancer and expression of each target gene were computed per cell type with Seurat (v5.3.0)⁸⁴ AggregateExpression as log2(CPM + 1) and scaled by row. Each target gene was represented by its highest-scoring enhancer, and rows were ordered by the cell type with the highest accessibility.

To assess concordance between age-associated changes in enhancers and their target genes, ABC-predicted enhancers of genes differentially expressed with age (FDR < 0.1) were classified by the direction of their accessibility change between very old and young mice, without applying a significance threshold to the enhancer, since a regulatory element need not change accessibility significantly to contribute to a change in target gene expression. Enhancers were retained when their accessibility changed in the same direction as their target gene, giving enhancers of age-upregulated and age-downregulated genes; enhancers of genes not differentially expressed with age formed the remaining group. The log2 fold change in expression of each target gene between very old and young hepatocytes was then compared between groups using two-sided Wilcoxon rank-sum tests.

### Chromatin accessibility peak calling

To define candidate cis-regulatory elements (cCREs), ATAC fragments were aggregated into pseudobulk profiles for each cell type, each cell type within each age group, the senescent cells of each cell type, all senescent cells, and each individual sample, so that peaks specific to particular ages, senescent states or animals would be captured. Peaks were called on each profile with MACS3 (v3.1.0)^89,90^ using the parameters ‘--nomodel --shift -75 --extsize 150 --call-summits -q 0.05 -g mm’, and summits on non-canonical chromosomes were removed. Summits were extended to fixed-width 500-bp peaks, and peaks overlapping the mm10 ENCODE blacklist or extending beyond chromosome boundaries were removed.^91^ Overlapping peaks were resolved with an iterative overlap procedure^92^: within each profile, the peak with the highest MACS3 score was retained and overlapping peaks were discarded iteratively, after which scores were normalized to score-per-million (SPM) to make them comparable across profiles of different depth.^92^ Profiles were then combined, and overlaps were resolved by SPM in the same way, yielding a union set of 475,985 cCREs. The same procedure applied within the profiles of each cell type produced cell type-specific peak sets, and cCREs of each cell type were defined as union peaks overlapping that cell type’s peak set (for example, 290,254 cCREs in hepatocytes). A cell-by-peak count matrix was generated from the union peak set with Signac’s ‘FeatureMatrix’ function.^85^

### Paired-Tag library generation

#### Nuclei permeabilization

Sorted nuclei were centrifuged at 300 × g for 3 min followed by 500 × g for 7 min. Pellets were resuspended in 250 µL of lysis/OMNI buffer (10 mM Tris-HCl pH 7.5, 10 mM NaCl, 3 mM MgCl₂, 0.1% IGEPAL CA-630, 0.1% Tween-20, 0.01% digitonin, 1 mM DTT, 1× protease inhibitor cocktail, 0.5 U/µL SUPERaseIn, 0.5 U/µL RNaseOUT) diluted to 0.1× detergent concentration, mixed by pipetting 10 times, transferred to 0.5 mL LoBind tubes and incubated for 3 min on ice. Nuclei were pelleted (500 × g, 6 min, 4 °C), resuspended in 100 µL 1× wash buffer (20 mM HEPES, 150 mM NaCl, 0.5 mM spermidine, 1× protease inhibitor cocktail, 2% BSA, 0.5 U/µL SUPERaseIn, 0.5 U/µL RNaseOUT), and pelleted again under the same conditions. Nuclei were then resuspended in Incubation Buffer from the Epigenome Technologies Multiplex Droplet Paired-Tag kit (MDP16101), counted, and kept on ice.

#### Antibody–PA-Tn5 complex assembly and nuclei incubation

Paired-Tag was performed with the Epigenome Technologies Multiplex Droplet Paired-Tag kit according to the manufacturer’s protocol, with the modifications described here. Incubation Buffer 1 was prepared by combining Incubation Buffer, Permeabilizer, kit RNase inhibitor and Stabilizer (90.5, 1.5, 7.5 and 0.5 µL per reaction, respectively) and kept on ice for no more than 5 h. Primary antibodies against H3K27ac (Abcam Ab177178), H3K27me3 (Abcam Ab6002) and H3K9me3 (Abcam Ab176916) were each pre-incubated with the tagmentation enzyme (PA-Tn5) in Incubation Buffer 1. Amounts were calculated for 200,000 nuclei per reaction using the kit’s Paired-Tag input optimization worksheet: 0.33, 0.97 and 0.39 µL antibody and 0.48, 1.06 and 0.54 µL enzyme for H3K27ac, H3K27me3 and H3K9me3, respectively, with Incubation Buffer 1 added to a final volume of about 50 µL. Complexes were assembled in PCR strip tubes (mixed by pipetting 10 times, then briefly centrifuged) and rotated at 18 rpm for 1 h at room temperature.

Each animal was assigned its own barcode (tag set A–D), giving 12 reactions (4 animals × 3 histone marks). Approximately 200,000 permeabilized nuclei per reaction were pelleted (500 × g, 7 min, 4 °C), leaving about 5 µL of supernatant when no pellet was visible. Pellets were resuspended in 25 µL Incubation Buffer 1 by gentle pipetting and transferred into the pre-assembled antibody–enzyme mixture. Reactions were mixed gently, avoiding bubbles, and incubated overnight at 4 °C on an 18 rpm rotator.

#### Tagmentation quality control

Before droplet generation, an aliquot of nuclei sufficient for the droplet step was set aside on ice. A portion of the remaining nuclei was used to check tagmentation. Tagmented DNA was purified with the MinElute PCR Purification Kit (Qiagen): 5 volumes of Buffer PB and 10 µL of 3 M sodium acetate (pH 5) were added per volume of reaction, the mixture was bound to a MinElute column, washed with 750 µL Buffer PE, dried by an additional spin, and eluted in 11 µL water. Purified DNA (10.5 µL) was amplified in a 25 µL reaction containing 1 µL each of 10 µM Nextera i5 and i7 primers and 12.5 µL NEBNext Q5 2× Master Mix. Cycling was 72 °C for 5 min; 98 °C for 30 s; 13 cycles of 98 °C for 10 s, 63 °C for 30 s and 72 °C for 1 min; then 72 °C for 2 min, with a 4 °C hold. A 1:10 dilution of the product was analyzed on an Agilent TapeStation to confirm the expected nucleosomal fragment size pattern. Nuclei that passed QC proceeded to droplet generation and library construction following Step 2 of the 10x Genomics Multiome protocol with 20k – 30k nuclei loaded into each droplet reaction. Libraries were paired-end sequenced on a NovaSeq X Plus to a target depth of ∼30,000 reads for DNA and ∼50,000 reads for RNA.

### snm3C-seq nuclei isolation and 3C procedure

10-15 mg of fresh frozen liver was transferred to gentleMACS M tubes containing ice-cold MACS buffer (5 mM CaCl₂, 3 mM magnesium acetate, 2 mM EDTA, 10 mM Tris-HCl pH 8, 0.6 mM DTT, and 1× protease inhibitor (Pierce A32965), thawed on ice for approximately 60 s, and homogenized on a gentleMACS Octo Dissociator using the Protein_01_01 program (Miltenyi Biotec). Homogenate was filtered through a 30μm CellTrics filter (Sysmex, 04-0042-2316) into a LoBind tube (Eppendorf, 22431021) and pelleted (1000 rcf, 10 min at 4°C) (Eppendorf, 5920 R). Pellets were resuspended in 2% formaldehyde in DPBS for 10 minutes at room temperature on a rotator. 95 μL of 2.5M Glycine was added, and incubated for 5 min at room temperature on a rotator. Homogenate was centrifuged at 1,000 x g for 10 min at 4°C (in bucket rotor). Supernatant was removed and the pellet was resuspended in 1mL of cold DPBS then centrifuged at 2,500 x g for 5 min at 4°C (in bucket rotor). Supernatant was removed and the pellets were snap-frozen in liquid nitrogen and stored in −80°C. Nuclei were subjected to in situ 3C following using the Arima Single Cell 3C Kit (Arima Genomics), with some modifications. Pellets were resuspended in 20uL of water and 24uL of Conditioning Solution, and transferred to a 0.2 mL PCR tube, and mixed gently by pipetting. Tubes were incubated at 62°C for 30 min in a thermal cycler, with the lid temperature set to 85°C. 20µL of Stop Solution 2 was added, and mixed gently by pipetting. Tubes were incubated at 37°C for 15 min in a thermal cycler with a lid temperature set to 85°C. 28µL of the restriction enzyme digestion mix (9.8uL water, 9.2uL Buffer H, 4.5uL Enzyme H1, 4.5uL Enzyme H2) was added and gently pipette mixed. Tubes were incubated in a thermal cycler for 37°C for 60 min, 65°C for 20 min, then 25°C for 10 min. 82uL of ligation mix was added (70uL Buffer C, 12uL Enzyme C) and gently mixed by pipetting, then incubated at 25°C for 15 min. Nuclei were transferred to a tube containing 850μL 1% BSA in DPBS on ice and filtered through a 30um Celltrics strainer to a new tube on ice. Nuclei were centrifuged at 1,000 x g for 10 min at 4°C (in bucket rotor). Supernatant was removed and pellets were resuspended each sample in 800uL of 1% BSA in DPBS. Nuclei were stained with DRAQ7 at 1:200 and mixed gently with a pipette and incubated on ice for 5 min. One tube of Proteinase K (Zymo cat. no. D3001-2-D) was dissolved in 1mL of Proteinase K Resuspension Buffer (Zymo cat. no. D3001-2-B) then added to a mix of 15mL M-Digestion Buffer 2x (Zymo cat. no. D5021-9) and 14mL water to make pK Digestion Buffer. 2uL of pK Digestion Buffer were added to each well of an Eppendorf twin.tec® PCR Plate 384 LoBind®, skirted, 45 µL, PCR clean (Eppendorf cat. No. 0030129547). Single DRAQ7 positive nuclei were sorted into individual wells of the 384-well plate using a Sony SH800 cell sorter. Plates were sealed then incubated in a thermal cycler at 50°C for 20 min with a heated lid at 85°C for nuclei digestion.

### Library generation and illumina sequencing for snm3C-seq

The snm3C-seq bisulfite conversion procedure and library preparation protocol was carried out as previously described.^93,94^ This procedure was automated using the Beckman Coulter Biomek i7 liquid handlers for all reactions in 384 and 96-well plates. The snm3C-seq libraries were sequenced on the NovaSeq X Plus 25B flow cell at 150 paired-end reads to ∼2 million raw reads per cell.

### Mapping of snm3C-seq

Sequencing data was processed using YAP (cemba-data v1.6.9).^93^ Following demultiplexing of reads by cell barcode, each library was aligned to the mm10 reference genome in two rounds using Bismark (v0.20.0)^95^ with the Bowtie aligner (v1.3.1).^96^ The resulting BAM files were sorted, deduplicated, and quality-checked using SAMtools (v1.9)^97^ and Picard (v2.20.4).^98^ ALLCools (v1.0.23)^93^ was subsequently used to generate per-cell methylation counts and identify chromatin contacts from the same sequencing alignments.

### Quality control of snm3C-seq

Nuclei were retained if they had an R1 mapping rate above 0.5, more than 3 × 10⁷ deduplicated mapped bases, a global mCG level above 0.5, a global mCH level below 0.2 and an mCCC level below 0.03, the last serving as an estimate of the bisulfite non-conversion rate. After quality control, 4,914 nuclei from 14 mice were retained for downstream analyses. Two of the old mice were vehicle-treated controls from a separate study.

### Methylome clustering

DNA methylation was quantified in non-overlapping 5-kb genomic bins and in gene bodies extended by 2 kb on each side (GENCODE vM22), and combined into a methylome cell dataset using ALLCools (v1.1.1).^93^ For clustering, bins were removed if they overlapped the mm10 ENCODE blacklist (v2) by at least 20%^91^ or lay on chrX, chrY or chrM. CG hypomethylation scores were binarized at 0.95, and bins with low coverage were removed (291,485 bins retained). Dimensionality was reduced by latent semantic indexing (LSI), an approach adapted from single-cell chromatin accessibility analysis,^99^ and the 18 components that passed a significance test (P < 0.08) were retained. Clusters were identified by consensus Leiden clustering (500 runs, resolution 0.2, 25 nearest neighbours, consensus rate 0.5, minimum cluster size 10), based on the Leiden community detection algorithm,^100^ which gave 13 clusters. A supervised classifier reassigned these clusters with a tenfold cross-validation accuracy of 0.987. Cells were visualized by UMAP.^101^

### Integration with 10x Multiome and cell type annotation

To annotate the methylome clusters, snm3C-seq nuclei^102^ were integrated with 10x Multiome snATAC-seq data (10x Genomics), randomly downsampled to at most 10,000 nuclei per cell type (50,905 nuclei in total) so that abundant cell types did not dominate the integration. ATAC fragments were aggregated into the same 5-kb bins. Both matrices were binarized and filtered in the same way, restricted to shared bins and concatenated. They were then decomposed jointly by latent semantic indexing (LSI),^99^ and the first 10 components were corrected for assay with Harmony (θ = 10, λ = 0.1).^88^ For each snm3C-seq nucleus, the 15 nearest snATAC-seq nuclei were identified in the corrected space with PyNNDescent; approximate nearest-neighbour search, Euclidean distance,^103^ and cell type labels were transferred by a distance-weighted vote. Methylome clusters were assigned to seven cell types on the basis of the transferred labels and of gene body mCG at canonical marker genes (Supplementary Fig. 2A), following the general strategy of cross-modality cell type annotation used in single-cell multiomic studies.

### Contact processing and imputation

Single-cell snm3C contacts aligned to the mouse reference genome (mm10) were filtered to remove blacklisted genomic regions using hicluster filter-contact and stored as contact.rmbkl tables^91^. Raw pseudobulk contact matrices were generated using hicluster merge-cell-raw at 100-kb and 25-kb resolutions, with min_pos_dist set to 0. Imputation was prepared using hicluster prepare-impute at 100-kb resolution, with the window size, step size and output distance each set to 500 Mb. Single-cell imputed coolers were also generated at 25-kb resolution using scHiCluster^104^. All downstream analyses described here used blacklist-filtered raw contacts or raw pseudobulk contact matrices, rather than imputed contacts. Although imputation can improve the representation of sparse single-cell contact maps, it estimates unobserved interactions and can consequently alter genome-wide contact distributions and apparent differences between age groups^104^.

### Genome-wide contact matrices

For each age group, raw contacts from all hepatocytes were merged into a single 100-kb-resolution cooler^105^. Mitochondrial and Y chromosomes were excluded. Contact matrices were ICE-balanced using the corresponding cooler bin weights, following the iterative correction approach^106^. Balanced pixel values were normalized to counts per million (CPM) by dividing by the total number of raw contacts in the corresponding matrix and multiplying by 10⁶. The resulting 100-kb pixels were aggregated into 6-Mb bins, with each 6-Mb pixel representing the mean contact value across the constituent 100-kb bin pairs. The Young matrix was displayed on a logarithmic scale. Age-associated differences were calculated by subtracting the Young matrix from the Very Old matrix and visualized using a diverging colour scale centred at zero. These genome-wide matrices were not depth-downsampled.

### Cis and trans contact fractions

Cis contacts were defined as pairs of loci on the same chromosome, whereas trans contacts were defined as pairs of loci on different chromosomes. The trans-to-total contact ratio was calculated as trans/(cis + trans). Contacts were counted per cell from the raw contact tables, and cells with fewer than 1,000 contacts were excluded. Counts were summed across cells within each mouse before calculating the ratio. As each mouse belonged to a single age group, comparisons between age groups were unpaired. Statistical significance was assessed using two-sided Student’s t-tests.

### Locus-specific contact maps

Contact maps for chr5:100–110 Mb were generated from equal-depth-downsampled, 25-kb-resolution hepatocyte pseudobulk coolers, with one matrix generated for each age group. Contact counts were normalized to CPM by dividing raw counts by the total number of contacts in the corresponding cooler and multiplying by 10⁶. ICE balancing was not applied. Matrices were displayed as 45°-rotated, near-diagonal contact maps using a shared colour scale across age groups. Difference matrices were calculated by subtracting the Young CPM matrix from the corresponding age-group matrix and visualized using a shared diverging colour scale. Blue indicates fewer contacts relative to Young. These differences represent absolute CPM differences rather than log₂ fold changes.

### Topologically associating domain identification

Consensus topologically associating domains (TADs) were identified using a pooled contact matrix independent of individual age groups. Equal-depth-downsampled 50-kb-resolution coolers from Young, Old and Very Old hepatocytes were summed before TAD calling. The 50-kb matrices were generated by merging adjacent bins from the depth-matched 25-kb matrices using hicMergeMatrixBins^107^ (two bins). TADs were identified using HiCExplorer hicFindTADs^107^,with a minimum depth of 150 kb, maximum depth of 500 kb, step size of 50 kb, delta of 0.01 and Benjamini–Hochberg false discovery rate correction at q = 0.05. At 25-kb resolution, the corresponding depth window was 75–250 kb, with the other parameters unchanged. X, Y and mitochondrial chromosomes were excluded when assigning contacts to TADs.

### Intra-TAD contact fraction

Using the fixed 50-kb consensus TAD set, raw single-cell contacts were classified according to the corresponding per-cell contact tables. A contact was defined as intra-TAD if both anchors fell within the same TAD. The denominator comprised all contacts in each cell, including both cis and trans contacts. Intra-TAD and total contact counts were summed across cells within each mouse, and the intra-TAD contact fraction was calculated as intra-TAD contacts divided by total contacts. Statistical significance was assessed using a one-sided Student’s t-test for the directional hypothesis that the intra-TAD contact fraction was lower in Very Old than in Young hepatocytes.

### Paired-Tag data processing and quality control

H3K27ac, H3K27me3 or H3K9me3 was profiled jointly with the transcriptome in single nuclei from young (4 months), old (22 to 24) months and very old (30 months) mice (14), yielding 40 sample-by-mark libraries, using the Paired-Tag approach.^108,109^ Multiplexed libraries were demultiplexed and filtered for multiplets. First FASTQ files from DNA libraries were demultiplexed to separate sequences corresponding to each sample. From each 10x droplet reaction reads from each of 4 combined samples were separated based on Tn5 adapters, as each sample was tagmented with a different pA-Tn5 containing a unique 2-nucleotide sequence in the 5’ end of the AdapterA oligo (/5Phos/NNTCGTCGGCAGCGTCAGATGTGTATAAGAGACAG) and a corresponding complimentary sequence in the pMents-Bridge oligo (/5Phos/CTGTCTCTTATACACATCTGACGCTGCCGACGANNCAGACGCG), allowing sample multiplexing into the same 10x droplet reaction. FASTQ files were aligned to the mosue reference genome (mm10, gencode vM23) separately for each modality using Cell Ranger ATAC (v2.2) and Cell Ranger RNA (10x Genomics, v8.0.1) with --chemistry ARC-v1 and otherwise default parameters. Cells from chromatin profiles were matched to their corresponding RNA profiles within the same nucleus via the shared cellular barcode provided by Cell Ranger ARC barcode whitelist. Cell barcodes with >90% of the DNA reads containing 1 pA-Tn5 barcode were retained. Gene expression and chromatin data were imported into Seurat (v5.3.0),^84^ retaining genes detected in at least three nuclei. Chromatin fragments were counted with Signac (v1.14.0)^85^ and annotated with EnsDb.Mmusculus.v79. In each library, putative doublets were identified with DoubletFinder (v2.0)^83^ using RNA profiles (2,000 variable genes, 20 principal components; pK chosen per library by maximizing the bimodality coefficient, pN = 0.25, expected doublet rate 5% adjusted for homotypic doublets),^83^ and were flagged for later removal.

### Integration, clustering and doublet removal

Libraries were merged (178,484 nuclei), and nuclei with more than 200 detected genes, less than 10% mitochondrial transcripts and more than 200 fragments in peaks were retained (168,543 nuclei). Counts were normalized per library with SCTransform,^86^ principal component analysis was performed on protein-coding variable genes, and library effects were corrected with Harmony using Seurat’s IntegrateLayers function.^84,88^ Clusters were identified with the Louvain algorithm^110^ on the first 30 Harmony dimensions and visualized by UMAP.^101^ To remove residual multiplets, nuclei were over-clustered (resolution 15), and clusters in which more than 30% of nuclei were DoubletFinder doublets, or which showed low quality or co-expressed markers of multiple lineages, were removed, together with all remaining DoubletFinder doublets.^83^ After a second round of normalization, integration and over-clustering (resolution 20), further low-quality or mixed-identity clusters were removed, leaving 138,392 nuclei.

### Cell type annotation

Cell type labels were transferred from the 10x Multiome reference using Seurat’s FindTransferAnchors and TransferData functions (30 reference principal components),^84,85^ and clusters were annotated as hepatocytes, cholangiocytes, hepatic stellate cells, endothelial cells, myeloid cells, T cells and B cells on the basis of the transferred labels and canonical marker expression (Supplementary Fig. 2A).

### Manifold drift

Transcriptional drift was quantified for each nucleus as its mean distance to the 20 nearest young nuclei of the same cell type in Harmony-corrected principal component space (30 components). Within each cell type, all nuclei from young mice were used as the reference set; for every nucleus the 20 nearest young neighbours were identified by Euclidean distance and their distances averaged. Because young nuclei belong to the reference set themselves, their drift values are expected to be lower than those of old and very old nuclei. Drift was computed separately for 10x Multiome RNA (Fig. 2A) and Paired-Tag RNA (Fig. 2G,H), in each case within the embedding used for clustering of that dataset, and compared between age groups with two-sided Wilcoxon rank-sum tests.

### Histone mark peak calling

Paired-Tag fragments were pooled by cell type across all mice. For H3K9me3 and H3K27me3, broad enrichment was called for each cell type with MACS2 (v2.2.9.1) in paired-end fragment mode (-f BEDPE --broad --nomodel -g 1.87e9 -q 0.05).^90^ For analyses of H3K9me3 heterochromatin, peaks within 1 kb of each other were merged with bedtools merge (-d 1000) to define H3K9me3 domains.^111^ Because H3K27ac marks active regulatory elements, we used the 10x Multiome cCREs as the reference regions for this mark rather than calling peaks de novo. Each cCRE was extended by 750 bp on each side to give 2-kb windows, and overlapping windows were merged.

### Coupling of manifold drift to histone modification signal

For each Paired-Tag nucleus, the fraction of reads in peaks (FRiP) was computed as the proportion of that nucleus’s fragments falling within the peak set of its histone mark and cell type (see Histone mark peak calling). FRiP was compared with the manifold drift of the same nucleus, derived from its matched RNA profile, using Spearman correlation, computed separately for each cell type and histone modification in very old mice only.

### Differential H3K9me3 domains

H3K9me3 domains were defined as described above (Histone mark peak calling). For each cell type, fragments per domain were counted per nucleus from each sample’s fragment file using Signac FeatureMatrix,^85^ and nuclei with fewer than 200 fragments in domains were excluded (54,879 nuclei retained). Counts were summed per mouse, combining both libraries of each animal, and differential domains were identified with edgeR (v4.4.2).^39^ Domains were filtered with filterByExpr, libraries were normalized using the trimmed mean of M-values (TMM) method, and a quasi-likelihood negative binomial model with age group as the sole covariate was tested with glmQLFTest for old versus young, very old versus young and very old versus old. Domains with Benjamini–Hochberg FDR < 0.05 were considered gained (log₂ fold change > 0, older relative to younger) or lost (log₂ fold change < 0) with age.

### Annotation and enrichment

Domains overlapping ±1 kb of an annotated transcription start site (GENCODE(vM22)) were classed as promoter-associated. Domains were assigned to the nearest gene with HOMER annotatePeaks.pl^112^ when the transcription start site lay within 50 kb; results were similar when all genes within ±50 kb were assigned. Gene ontology (biological process) enrichment was computed with clusterProfiler enrichGO,^113^ using genes linked to all tested domains in the same cell type and comparison as background, a minimum gene set size of 10 and Benjamini–Hochberg-adjusted q < 0.05; redundant terms were collapsed with simplify (similarity cutoff 0.7).

### Expression of genes linked to age-altered H3K9me3 domains

For each cell type and direction (lost or gained), genes linked to domains differential between very old and young mice (nearest TSS within 50 kb) were intersected with genes detected in the 10x Multiome snRNA-seq data and scored per nucleus with Seurat AddModuleScore⁸⁴ (seed 42), separately for each cell type. Gene sets with fewer than 3 genes and cell types with fewer than 20 nuclei were skipped. Scores were compared between young and very old nuclei using two-sided Wilcoxon rank-sum tests.

### Senescence-associated enhancers

Within each cell type, Spearman correlation was computed between the TF-IDF-normalized accessibility of each peak and the SenLearner senescence probability across all cells, with P values from a t-test (n − 2 degrees of freedom) adjusted by the Benjamini–Hochberg method within each cell type. All peaks were tested except those with zero variance across cells. Peaks with |ρ| > 0.02 and adjusted P < 0.05 that coincided with non-promoter ABC^46^-predicted enhancers (ABC^46^ score > 0.02; see Enhancer target gene predictions) in the same cell type were considered senescence-associated, and were classified as SnC-Up (positive correlation) or SnC-Down (negative correlation), giving 5,102 unique enhancers (4,567 SnC-Up and 535 SnC-Down). For display (Fig. 4A), enhancers that were senescence-associated in more than one cell type were assigned to the cell type with the largest |ρ|. To relate these enhancers to target gene expression, all genes linked to each enhancer by the ABC^46^ model (ABC^46^ score > 0.02) were taken as its targets, and the Pearson correlation between each gene’s expression and senescence probability across cells of the same cell type was used (see Senescence-correlated genes). Within each cell type, these correlations were compared between targets of SnC-Up enhancers, targets of SnC-Down enhancers and all ABC^46^-predicted target genes using two-sided Wilcoxon rank-sum tests (SnC-Down versus all, SnC-Up versus all, and SnC-Down versus SnC-Up), with Benjamini–Hochberg correction across all tests (Fig. 4B). Genes linked to both SnC-Up and SnC-Down enhancers were included in both groups. Cholangiocytes had no SnC-Down enhancers and were tested only for SnC-Up targets.

### Gene regulatory network construction

Gene regulatory networks were constructed for hepatocytes, separately for genes upregulated and downregulated in senescent relative to non-senescent cells. Transcription factor nodes were selected from known motifs enriched in DARs that gained (AP-1, IRF1, NF-κB, STAT1 and p53) or lost (HNF4α, PPARα and C/EBP) accessibility in senescent hepatocytes; enrichment was tested with HOMER^112^ (v5.1) findMotifsGenome.pl (-size given) against all peaks included in the differential accessibility analysis, and motifs with Benjamini–Hochberg q < 0.05 were considered enriched. p53 was represented by the HOMER^112^ motif p53(p53)/mES-cMyc-ChIP-Seq and served as a positive control, given its established induction of *Cdkn1a* during senescence. A transcription factor was connected to a gene when three conditions were met. First, a DAR between senescent and non-senescent hepatocytes (MAST^114^, Benjamini–Hochberg-adjusted P < 0.01, either direction) contained an instance of that factor’s motif (see Motif occurrences for gene regulatory networks). Second, the ABC^46^ model linked that DAR to the gene (ABC^46^ score > 0.02; promoter elements included). Third, the gene was differentially expressed between senescent and non-senescent hepatocytes (MAST^114^, Benjamini–Hochberg-adjusted P < 0.01) with a log2 fold change greater than 0.3 (upregulated network) or less than −0.3 (downregulated network), and was detected in more than 20% of senescent hepatocytes.Where several elements linked the same factor to the same gene, they were collapsed into a single edge carrying the maximum ABC^46^ score. Target genes were ranked by the product of the maximum ABC^46^ score, |log2 fold change| and −log10(adjusted P), and the eight highest-ranking targets of each factor together with the top-ranking targets overall were displayed, up to 40 genes. Genes linked to more than one transcription factor were considered shared. Networks were drawn with ggraph (v2.2.1), igraph (v2.1.4) and tidygraph (v1.3.1) using a Fruchterman–Reingold layout, with edge width proportional to the ABC^46^ score and node colour indicating the log2 fold change in expression between senescent and non-senescent hepatocytes.

### Chromatin state segmentation

Chromatin states were learned for hepatocytes using ChromHMM (v1.28) from Paired-Tag H3K27ac, H3K27me3 and H3K9me3 profiles.^115^ To capture the full range of hepatocyte chromatin, fragments were pooled across ages and across senescent and non-senescent cells. Chromatin accessibility was excluded so that state definitions were independent of the accessibility data used to identify differentially accessible regions. Fragments were binarized in 200-bp bins using BinarizeBed, with fragment midpoints (-center) and the default Poisson threshold, and models were trained using LearnModel. An eight-state model was selected, and states were annotated based on their emission probabilities.

### Enrichment of differentially accessible regions across chromatin states

Differentially accessible regions (DARs) between senescent and non-senescent hepatocytes were identified with MAST^114^ on peaks accessible in at least 5% of cells in either group, and peaks with Benjamini–Hochberg-adjusted P < 0.01 were retained (8,853 gained and 10,240 lost). Each DAR and each hepatocyte cCRE (n = 290,254), used as background, was extended to at least 200 bp around its midpoint and assigned to the chromatin state with which it shared the greatest base-pair overlap. For each state, enrichment of gained or lost DARs relative to all cCREs was tested with two-sided Fisher’s exact tests. Odds ratios are conditional maximum-likelihood estimates with 95% confidence intervals, and P values were adjusted across all 16 tests (8 states × 2 directions) by the Benjamini–Hochberg method. Enrichments were considered significant at q < 0.05 with |log₂ odds ratio| ≥ 0.5. To exclude effects driven by promoters, the analysis was repeated for regions more than 1, 2 or 4 kb from the nearest transcription start site, with Benjamini–Hochberg correction applied within each subset.

### State-specific motif enrichment

Known motif enrichment was assessed for gained DARs in E8 and lost DARs in E3, the two states enriched in only one direction, using HOMER findMotifsGenome.pl (mm10, -size given, -nomotif) on 200-bp windows centered on region midpoints.^112^ Background regions comprised non-differential hepatocyte cCREs assigned to the same chromatin state, randomly sampled to a maximum of 50,000 regions using a fixed random seed (7). This state-matched background was used to identify motif enrichment specific to DARs relative to other elements within the same chromatin state.

### SenLearner: senescence labelling and classification

Senescent cells were identified separately within each cell type using SenLearner, a two-step framework that derives training labels from a senescence signature and then trains a classifier to score all cells. Counts were log-normalized per sample, and each cell was scored for the SDTMC senescence gene set using Seurat’s AddModuleScore, after excluding proliferation-associated genes (*Ccnd1, Ccnd2, Ccna2, Ccnb1* and *Lmnb1*).^84^ Within each cell type, cells in the top 10% of SDTMC scores that also expressed at least one cyclin-dependent kinase inhibitor gene (*Cdkn1a, Cdkn1b, Cdkn1c, Cdkn2a, Cdkn2b, Cdkn2c* or *Cdkn2d*; log-normalized expression > 0.5) were labelled senescent. Within each sample, cells in the bottom 30% of scores were labelled non-senescent; remaining cells were left unlabelled.For each cell type, an L1-regularized logistic regression classifier (scikit-learn v(1.6.0); C = 0.5, saga solver, balanced class weights, maximum 2,500 iterations)^116^ was trained on labelled cells using log-normalized expression of feature set; low-quality and cell cycle-associated genes excluded and applied to all cells to estimate senescence probabilities. Cells were classified as senescent if their predicted probability exceeded the lowest probability among senescent training cells of that cell type (e.g., 0.974 for hepatocytes).

### Senescence-correlated genes

Within each cell type, Pearson correlation between log-normalized gene expression and senescence probability was computed across all cells, with P values from a t-test (n − 2 degrees of freedom) adjusted by the Benjamini–Hochberg method. Genes with |r| > 0.05 and adjusted P < 0.01 were considered senescence-correlated. Genes meeting these criteria in hepatocytes, cholangiocytes, myeloid cells and T cells in the same direction were defined as the shared senescence signature (26 genes). Senescence-correlated genes were compared with the 147-gene SDTMC list to classify them as recovered, missed or additional.

### Differential expression and accessibility between senescent and non-senescent hepatocytes

Hepatocytes were classified as senescent or non-senescent using SenLearner. Differentially expressed genes were identified using Seurat’s FindMarkers function with the MAST method,^84,114^ and genes with Benjamini–Hochberg-adjusted P < 0.01 were retained. Differentially accessible regions (DARs) were identified using MAST^114^ on peaks accessible in at least 5% of cells in either group. Testing was performed in chunks of 5,000 peaks, with P values adjusted genome-wide using the Benjamini–Hochberg method. Regions with adjusted P < 0.01 were considered differentially accessible (8,853 gained and 10,240 lost). Additional expression filters used in individual analyses are described in the relevant sections.

### Hepatocyte senotype identification

Senescence-associated genes were defined as those upregulated in senescent relative to non-senescent hepatocytes (MAST^114^; Benjamini–Hochberg-adjusted P < 0.01, average log2 fold change > 0, detected in more than 10% of cells; n = 1,436). Log-normalized expression of these genes across the 6,342 hepatocytes classified as senescent by SenLearner was used to compute a gene–gene Pearson correlation matrix, and genes were grouped by hierarchical clustering with Ward’s D2 linkage on the correlation distance (1 − r), cutting the tree into four modules designated senotypes S1–S4. Within each module, module membership was defined as the mean correlation of a gene with the remaining genes of that module and tested by Fisher Z transformation as a heuristic membership filter; the 527 genes with a Benjamini–Hochberg-adjusted P < 0.05 were retained, and a module eigengene was computed as the first principal component of their expression (prcomp, centred, not scaled). The number of modules was evaluated across k = 3–12 using silhouette scores, eigengene variance explained and the proportion of genes retained after membership filtering, and the clustering was compared with an alternative solution using 1 − |r| as the distance metric (adjusted Rand index; mclust (v6.1.1), cluster (v2.1.8.1)).

### Global mCG and definition of mCG fraction groups

The global mCG fraction of each snm3C-seq nucleus was calculated as methylated over total CG basecalls with ALLCools (v1.0.23)^77^. Analyses were restricted to endothelial cells, B cells, T cells and myeloid cells, the four cell types with a distinct low-mCG population in the snm3C-seq embedding. Within each cell type, global mCG was smoothed by averaging each nucleus with its 10 nearest neighbours in UMAP space, to reduce single-nucleus sampling noise. Low-mCG nuclei were then defined in two steps. First, nuclei in the lowest smoothed-mCG percentile of their cell type were selected (endothelial cells, 10%; T cells, 20%; B cells, 25%; myeloid cells, 12%). These cutoffs corresponded to the low-mCG population in the mCG distribution of each cell type. Smoothed mCG was bimodal in myeloid cells, T cells and B cells (two-component Gaussian mixture preferred over one; ΔBIC = 1,347, 103 and 364, respectively). In myeloid cells the low-mCG mixture component matched the cutoff (11.7% versus 12% of nuclei). In T cells and B cells, all selected nuclei fell within the low-mCG component (56% and 47% of nuclei), and the cutoffs took its lowest part, the most strongly hypomethylated nuclei. Second, the selected nuclei were clustered with DBSCAN^117^, so that low-mCG nuclei had to form spatially contiguous groups in the embedding and isolated low-mCG nuclei were excluded. Eps was scaled to the median distance to the 10th nearest neighbour (1.5 × for endothelial, T and myeloid cells and 2.0 × for the smaller B-cell population), which made it comparable across cell types of different size and density; minimum samples were 8, 8, 5 and 8, respectively. Because smoothed mCG was unimodal in endothelial cells, low-mCG endothelial nuclei were defined by this spatial criterion. Nuclei with smoothed mCG at or above the 50th percentile (35th for myeloid cells) were labelled high mCG, and the remainder intermediate. Low-mCG assignments were stable across the low-mCG percentile ±5, eps 1.0–2.5 × and minimum samples 5–10 (median Jaccard similarity to the chosen set, 0.66–0.81), except at the most restrictive combination (eps 1.0 × with minimum samples ≥ 8), where the smallest low-mCG populations were not detected.

### Pseudobulk methylation tracks

For each cell type and mCG group, single-nucleus ALLC files were merged with ALLCools (v1.1.1).^93^ merge-allc in batches of 140 nuclei, which were then merged together (bin length 50 Mb). CG sites were extracted with strands merged (extract-allc, CGN context), and mCG fraction tracks were generated with allc-to-bigwig at 100-bp resolution on mm10.

### H3K9me3 domain calling

H3K9me3 Paired-Tag fragments were pooled by cell type across all mice. Fragment midpoints were counted in 10-kb bins on autosomes and chrX and summed into 50-kb bins. Bins with fewer than 10% of the median non-zero count were removed as low mappability. Counts were converted to counts per million (CPM), and enrichment was calculated as log2((CPM + 0.1) / (median CPM + 0.1)), capped at the 99.5th percentile and smoothed with a 5-bin moving average (10 bins for the lower-depth T-cell and cholangiocyte profiles). For each cell type, a three-state Gaussian hidden Markov model (hmmlearn v0.3.3) was fit genome-wide, with chromosomes as separate sequences. State means were initialized at the 15th, 50th and 85th percentiles of enrichment, and self-transition probabilities at 0.99. States with mean log2 enrichment above 0.3 were considered enriched. Bins with a posterior probability of at least 0.3 for an enriched state were used as seeds and extended across contiguous bins with smoothed enrichment above 0.2. Segments within 100 kb of each other were merged, and domains of at least 200 kb were kept. These domains are distinct from the H3K9me3 domains used for differential analysis in Fig. 2, which were defined from MACS2^90^ broad peaks merged at 1 kb (see Histone mark peak calling).

### Methylation metaplots

The average mCG fraction of high- and low-mCG pseudobulks was calculated across each cell type’s own H3K9me3 domains (see H3K9me3 domain calling), with each domain scaled to a common length, and across 0.3-Mb upstream and downstream flanking regions, using deepTools^133^ computeMatrix scale-regions and plotProfile.

### Integration of snm3C-seq with 10x Multiome

mCG groups were transferred to 10x Multiome nuclei separately for each cell type, using the chromatin accessibility modality. Both modalities were summarized over a shared set of genome-wide 5-kb bins on autosomes, with mm10 ENCODE blacklist regions (v2) removed^91^. For snm3C-seq, CG hypomethylation scores were calculated per bin (ALLCools hypo-score) and binarized at 0.8. ATAC counts were binarized as accessible or not (at least one fragment). In both modalities, bins detected in few nuclei were removed with ALLCools filter_regions. Three feature sets were considered: all shared bins; bins among the 150,000 most variable in both modalities (ATAC, Scanpy seurat_v3 highly variable features; snm3C-seq, variance); and the 50,000 bins with the highest mean accessibility. The two modalities were concatenated and reduced by latent semantic indexing (ALLCools)^93,99^, keeping significant components (ALLCools significant_pc_test, P < 0.1; at most 30), and integrated with Harmony (λ = 0.1, 30 iterations)^88^. Five configurations were compared: (A) variable bins, θ = 2, 50 clusters; (B) all shared bins, first 10 components, θ = 10, 100 clusters; (C) variable bins, θ = 10, 100 clusters; (D) as A but with continuous hypomethylation scores; and (E) the 50,000 most accessible bins, θ = 2, 50 clusters. For each snm3C-seq nucleus, the 15 nearest ATAC nuclei in the integrated space were found with PyNNDescent^103^ and weighted by distance (w = 1 − exp(−0.25 × (1 − d/d₁₅)), normalized to sum to one). Each ATAC nucleus summed the weights it received from snm3C-seq nuclei of each mCG group, and was assigned to a group when that group contributed at least 60% of its total weight. ATAC nuclei below this threshold (ambiguous) or receiving no weight (unassigned) were excluded. Modality mixing was calculated as the fraction of ATAC nuclei among the 30 nearest neighbours of each snm3C-seq nucleus in the integrated space, averaged across snm3C-seq nuclei and divided by the overall fraction of ATAC nuclei, so that 1 indicates mixing in proportion to modality size. For each cell type we used the configuration with mixing of at least 0.5 and the highest age concordance: the Spearman correlation between the age of each snm3C-seq nucleus and the weighted mean age of its matched ATAC nuclei, a measure of matching accuracy given that both cohorts have known ages. Ties within 0.02 were broken by the difference in the weighted fraction of aged neighbours between low- and high-mCG nuclei, then by the overlap of their neighbours. This selected configuration E for T cells, C for B cells and A for myeloid and endothelial cells.

### Senescence probability of mCG groups

Per-nucleus senescence probabilities were calculated with SenLearner on 10x Multiome RNA (see SenLearner: senescence labelling and classification) and compared between low- and high-mCG nuclei with two-sided Wilcoxon rank-sum tests. Because nuclei from the same mouse are not independent, these nucleus-level P values are descriptive; a per-mouse paired comparison of mean senescence probability between low- and high-mCG nuclei is shown in Supplementary Figure 9.

### Statistics for mCG group analyses

Age associations (Fig. 6D) were tested at the level of individual mice, including mice with at least 20 nuclei of the relevant cell type, by two-sided Spearman correlation between the fraction of low-mCG nuclei and age in months. Analyses used R (v4.4.3) with Seurat (v5.3.0)^84^, and Python (v3.8.18) with ALLCools (v1.0.23)^93^, scikit-learn (v1.3.2)^116^, harmonypy (v0.0.10)^88^, PyNNDescent (v0.5.11)^103^ and hmmlearn (v0.3.3).

### Motif enrichment

Known motif enrichment was assessed with HOMER findMotifsGenome.pl (v5.1)¹¹² against the HOMER known vertebrate motif collection (mm10; -size given, -nomotif), with default GC normalization. Motifs with q < 0.05 were considered enriched. Foreground and background regions were defined separately for each analysis: (i) for enhancers of age-associated genes (Fig. 1M,N), ABC-linked enhancers of genes up- or downregulated with age in each cell type were tested against the cell type-specific accessible regions for the corresponding cell type; (ii) for senescence-correlated enhancers (Fig. 4D,E), enhancers whose accessibility correlated positively or negatively with senescence probability (|r| > 0.02, adjusted P < 0.05) were tested against all ABC-predicted enhancers in the same cell type; (iii) for senescence-associated DARs (Fig. 4F,G), regions gained or lost in senescent hepatocytes were tested against all peaks included in the differential accessibility analysis; (iv) for state-specific DARs (Fig. 4K), 200-bp windows centred on gained DARs in E8 or lost DARs in E3 were tested against up to 50,000 non-differential hepatocyte cCREs from the same chromatin state, randomly sampled with seed 7; and (v) for senotype regulatory elements (Fig. 5B), ABC-predicted hepatocyte enhancers linked to genes in each senotype module were tested against all ABC-predicted hepatocyte enhancers.

### Motif occurrences for gene regulatory networks

To link transcription factors to individual regulatory elements, each HOMER^112^ known vertebrate motif was scanned across the mm10 genome with scanMotifGenomeWide.pl (HOMER v(5.1)), and motif instances lying entirely within a cCRE of the union peak set were identified with bedtools intersect (-f 1.0). A cCRE containing at least one instance was considered motif-containing.

### Gene ontology enrichment

Over-representation of Gene Ontology biological process terms was tested using clusterProfiler (v(4.14.6))^113^ enrichGO with org.Mm.eg.db and Benjamini–Hochberg adjustment; terms with q < 0.05 were considered enriched. In every analysis, the background comprised all genes included in the test from which the gene set was derived. Gene sets were defined as follows: (i) age-associated genes (Fig. 1I,J), comprising genes up- or downregulated in very old versus young mice in each cell type (edgeR, FDR < 0.05), against all genes tested in that cell type; (ii) senescence-correlated genes (Fig. 3F,G), comprising genes positively or negatively correlated with senescence probability in each cell type (|r| > 0.05, adjusted P < 0.01), against all genes tested for correlation; terms enriched in only one cell type were defined as cell type-specific (Supplementary Fig. 4G); (iii) senotype module genes (Fig. 5C), against genes expressed in more than 10% of hepatocytes; and (iv) genes linked to H3K9me3 domains gained or lost with age (Fig. 2), against genes linked to all tested domains. For the H3K9me3 domain analysis, the minimum gene set size was 10, and redundant terms were collapsed using simplify (similarity cutoff 0.7).

### Enhancer target gene predictions

Enhancer–gene links were predicted separately for each cell type using the activity-by-contact (ABC) model,^46,118^ with cell type-specific snATAC-seq accessibility as the activity measure and cell type-specific pseudobulk snm3C-seq contact maps (5KB) as the contact measure. Unless otherwise specified, non-promoter elements with an ABC score > 0.02 were considered predicted enhancers.

### MERFISH sample preparation and imaging

Fresh-frozen mouse liver sections (12 μm) were mounted on MERSCOPE Large slides (Vizgen 20400118) and transported on dry ice, then equilibrated at −20°C for 30 min. Sections were fixed in 10 mL pre-warmed 4% PFA at 47°C for 30 min, washed three times for 5 min in 1x PBS with RNase inhibitor (1:5000; New England BioLabs M0314L) at room temperature, and permeabilized in 10 mL 70% ethanol at 4°C for 3 days. Autofluorescence was quenched by photobleaching in the MERSCOPE Photobleacher for 5 hours at room temperature. For cell boundary staining, sections were blocked for 1 hour at room temperature in 200 μL Blocking Solution (Blocking Buffer C Premix with 10 μL RNase inhibitor) under a parafilm cover, then incubated for 1 hour with Primary Staining Solution (Blocking Buffer C Premix, RNase inhibitor, and 2 μL Cell Boundary Primary Stain Mix). Sections were washed three times for 5 min in 1x PBS with gentle manual rocking, incubated for 1 hour with Secondary Staining Solution (Blocking Buffer C Premix, RNase inhibitor, and 6 μL Cell Boundary Secondary Stain Mix), washed three times in 1x PBS, post-fixed in 10 mL 4% PFA for 15 min at room temperature, and washed twice in 1x PBS. Sections were then washed twice for 1 min in 10 mL Conditioning Buffer (Vizgen 20300116) and incubated in 10 mL Conditioning Buffer with 10 μL RNase inhibitor at 37°C for 15 min. Anchoring pretreatment was performed by adding 200 μL Pre-Anchoring Reaction Buffer (200 μL Conditioning Buffer, 10 μL Pre-Anchoring Activator, 10 μL RNase inhibitor) directly to the sample, covering with parafilm, and incubating in a humidified chamber protected from light at room temperature for 16 hours. After pretreatment, sections were washed with 10 mL Sample Prep Wash Buffer (Vizgen 20300001) and incubated in 10 mL Formamide Wash Buffer (Vizgen 20300002) at 37°C for 15 min. RNA anchoring was carried out in 200 μL Anchoring Buffer (Vizgen 20300117) at 37°C for 2 hours in a humidified chamber, followed by a 10 min incubation in Formamide Wash Buffer at 47°C and a 2 min wash in Sample Prep Wash Buffer. Sections were then gel-embedded using 10 mL Gel Embedding Premix (Vizgen 20300004) with 50 μL 10% Ammonium Persulfate (Sigma 09913) and 5 μL TEMED (Sigma T7024). Sections were covered in gel embedding solution for 1 min, the solution was aspirated, and 200 μL of fresh solution was applied. Gel Slick (Lonza 50640)-treated coverslips (Corning 2855-25) were placed onto the sections with excess gel aspirated. Gel was allowed to polymerize for 1.5 hours at room temperature. Following embedding, sections were digested in 200 μL Digestion Mix at 37°C for 2 hours in a humidified chamber, and cell clearing was performed by incubating sections in 10 mL Clearing Solution at 47°C overnight in a humidified chamber. Sections were then washed 3x5 minutes in Sample Prep Wash Buffer and incubated in Formamide Wash Buffer at 37°C for 30 min. For probe hybridization, 200 μL Encoding Probe Mix (MERSCOPE 300 Gene Panel V2.0, panel ID B2P2153, 306 genes) was applied directly on the sample, covered with parafilm, and hybridized for approximately 21 hours at 47°C in a humidified chamber. Post-hybridization washes were performed twice in Formamide Wash Buffer at 47°C for 30 min each. 200 μL of Enhancer (Vizgen 20300194) was then applied to sections and covered with parafilm for a 24-hour hybridization at 37°C in a humidified chamber. Sections were washed twice with Enhancer Wash Buffer (Vizgen 20300192) at 47°C for 20 min, quickly washed twice in Sample Prep Wash Buffer, stained with DAPI and PolyT Mix for 15 min at room temperature with rocking, washed for 10 min with 10 mL Formamide Wash Buffer, and imaged on the MERSCOPE Ultra system (Vizgen 10000008) using the MERSCOPE Large 300 Gene Imaging Kit V 2.0 (Vizgen 10400172). A fully detailed, step-by-step instruction on the MERFISH sample prep can be found in Vizgen’s MERFISH 2.0 Sample Preparation User Guide for Sectioned Tissue Samples (Vizgen 91600132). Full instrumentation protocol can be found in MERSCOPE Ultra Instrument User Guide (Vizgen 91600131).

### MERFISH data processing

25 liver sections across 19 mice were imaged with a 300-gene MERFISH panel and segmented into single cells using DAPI-based Cellpose segmentation.^119^ Cells were removed if they failed any of the following criteria: fewer than 20 detected transcripts; fewer than 10 detected genes; segmented volume below 100 µm³ or above 9,000 µm³; anisotropy (elongation ratio) above 4.0; solidity (convexity of the segmentation boundary) below 0.50; or a blank-probe fraction above 0.10, where the blank fraction is the number of blank-probe counts divided by the total of real plus blank counts. These rules target empty or fragmentary segmentation masks, over-merged cells, implausibly shaped boundaries, and cells dominated by ambient signal.

Cell-type labels were transferred from the liver single-nucleus multiome reference using canonical-correlation co-embedding with mutual-nearest-neighbour anchors.^120^ Annotation was performed in two rounds, following the same strategy in Zhang et al.^121^ The first round integrated the reference with all 3,485,824 MERFISH cells and transferred a binary label only (Hepatocyte vs non-hepatocyte). The second round then took every non-hepatocyte from both modalities and transferred the six non-hepatocyte types. Cells were retained only if both rounds reached a confidence of 0.80. At this threshold 697,508 cells (20.0%) were excluded, leaving 2,788,316 confidently annotated cells.

### Mapping multiome senescent cells to onto MERFISH

Senescence labels were transferred from the multiome reference to MERFISH within each cell type separately. For each MERFISH section, the mapping was against multiome reference nuclei of the same age group and sex. Within a cell type, cosine similarity was computed between every multiome nucleus and every MERFISH cell over the 147 SDTMC genes. Each reference nucleus was linked to its top 300 most similar MERFISH cells, and links with cosine similarity < 0.2 were discarded. For each MERFISH cell, the cosine similarities of its links were summed over senescent (w₁) and non-senescent (w₂) multiome nuclei. A mapped senescence score was then computed as the proportion of the total similarity contributed by linked senescent nuclei: S_senescence_ = w₁ / (w₁ + w₂). Cells with fewer than three links were removed. For the remaining cells within each cell type, senescence labels were assigned based on prevalence matching. Cells were ranked by their senescence scores, and the top *k = round(π × N)* cells were classified as senescent, where *N* represents the number of remaining MERFISH cells and *π* denotes the senescent fraction in the age- and sex-matched multiome reference for the corresponding cell type. Other remaining cells were labeled as non-senescent.

### Cell-type neighborhood enrichment analysis

Spatial colocalization was assessed for mapped senescent and non-senescent cells across 25 MERFISH sections from 19 mice. Cells were classified into nine categories: seven non-senescent cell types and two senescent groups (senescent hepatocytes and senescent non-hepatocytes). Within each section, neighboring cells within a 50-µm radius were identified using Euclidean distances and a BallTree algorithm scikit-learn^116^, excluding the center cell. Neighbor counts and cell-type abundances were aggregated across all sections before calculating enrichment. For each center–neighbor category pair, fold enrichment was calculated as the observed proportion of neighbors belonging to a given category divided by its overall abundance. Statistical significance was assessed using two-sided exact binomial tests, with Benjamini–Hochberg correction.

### Identification of spatial clusters of senescent cells and SenSpots

Senescent cells were spatially clustered within each tissue section using DBSCAN,^117^ implemented in scikit-learn^116^, based on their physical coordinates, with a neighborhood radius of 50 µm (eps = 50) and a minimum of five cells per cluster (min_samples = 5). Senescent cells not assigned to any cluster were classified as isolated. Each identified cluster was subsequently expanded to define a senescence-enriched spatial neighborhood, termed as SenSpot. Specifically, cells within a tissue section, regardless of cell type or senescence status, were assigned to a SenSpot if they were within 50 µm to any clustered senescent cells. Each cell was assigned to the SenSpot corresponding to its nearest clustered senescent cell.

### Cell-type enrichment in SenSpots

All cells were included in the cell-type enrichment analysis regardless of senescence status and classified into seven major cell types. For each mouse, cell counts were pooled across sections. For each cell type, fold enrichment was calculated as its proportion among SenSpot cells divided by its proportion among all cells. Enrichment was assessed using one-sided Fisher’s exact tests, with Benjamini–Hochberg correction across the seven cell types.

### Mapping multiome senotype cells to onto MERFISH

Hepatocyte senotypes were defined in the multiome by clustering senescent hepatocytes based on senotype gene signature scores. Because S3 cells were predominantly derived from a single very old mouse, this senotype was excluded from the multiome reference prior to spatial mapping to minimize potential sample-specific effects. Senotype labels were then transferred to senescent MERFISH hepatocytes, using age- and sex-matched multiome senescent hepatocytes as the reference. Mapping was performed independently for each tissue section using the same similarity-based approach as the senescence label transfer described above. Senotype assignment was performed using a similarity-weighted voting approach. For each MERFISH cell, the cosine similarities of links were summed separately for each senotype and normalized by the total similarity weight across all links. Cells were assigned to the senotype with the highest weighted vote fraction. Cells without any retained links remained unassigned.

### Hepatocyte zonation analysis

Hepatocyte positions along the portal-central axis of the liver lobule were inferred from the expression of established zonation markers. Nine periportal markers (*Ass1*, *Cyp2f2*, *Asl*, *Arg1*, *Hsd17b13*, *Hal*, *Sds*, *Pck1*, and *G6pc*) and eight pericentral markers (*Glul*, *Cyp2e1*, *Axin2*, *Cyp1a2*, *Cyp2a5*, *Oat*, *Cyp7a1*, and *Slc1a2*) were selected based on previous studies^17,20,122,123^.

For all 91,185 hepatocytes, zonation scores were calculated as follows. First, log-normalized expression values were standardized (z-scored) for each zonation gene across all hepatocytes and then averaged within each marker set to calculate periportal and pericentral scores, respectively. Second, a continuous zonation score was defined as the difference between the pericentral and periportal scores, with higher values indicating stronger pericentral marker expression. Hepatocytes were then assigned percentile ranks based on their zonation scores across the entire hepatocyte population. To define discrete zonation bins, hepatocytes were divided into ten zone bins (zone 1 to zone 10) along the inferred portal-central axis. Zone boundaries were determined from the deciles of the zonation score distribution in young hepatocytes and applied consistently across all age groups.

### Zonation bias test of senotypes

Zonation bias was assessed for 6,342 senescent hepatocytes from 18 mice across ten lobular zones (Zone 1, most periportal; Zone 10, most pericentral). For each mouse and senotype, the zonal center of mass (CoM) was calculated as the mean zone bin index, and zonation bias (ΔCoM) was defined as the difference between the senotype-specific CoM and that of all senescent hepatocytes from the same mouse. Positive and negative ΔCoM values indicate pericentral and periportal enrichment, respectively. Only mice with ≥20 cells of a given senotype were included (S1, n = 17 mice; S2, n = 12 mice; S3, n = 2 mice; S4, n = 9 mice). The overall zonation bias was estimated as the cell-count-weighted mean ΔCoM across mice. Statistical significance was assessed using 10,000 bootstrap resamples of mice, with two-sided *P* values calculated from the bootstrap distribution. S3 was excluded from statistical testing due to insufficient mice.

### Cell–cell communication analysis of Wnt signaling

Cell–cell communication was inferred using CellChat^79^ (v2.1.2) in R 4.3.3, separately for vehicle- and MDM2i-treated mice (2 mice per group). Cells were grouped into seven annotated cell types, and log-normalized RNA expression was analyzed using the mouse CellChatDB. Overexpressed genes and ligand–receptor interactions were identified using default settings. Communication probabilities were calculated using computeCommunProb (10% truncated mean, 100 permutations), with population.size = FALSE to account for differences in cell-type abundance. Interactions involving cell groups with fewer than 10 cells were excluded.

Canonical Wnt ligand–receptor interactions targeting hepatocytes were extracted using subsetCommunication (*P* < 0.05).

### In vivo *Stat1* knockout in p21-expressing cells

Aged male B6J.129(B6N)-Gt(ROSA)26Sor^tm1(CAG-cas9*,-EGFP)Fezh^/J mice (JAX 026175), carrying a Cre-dependent CAG-LSL-3xFLAG-Cas9-P2A-EGFP cassette at the Rosa26 locus, were used at 24.2 to 28.3 months of age at tissue collection. All mice received an AAV8 vector expressing Cre recombinase under a p21 (*Cdkn1a*) promoter fragment (pAAV-p21>Cre; VectorBuilder VB230928-1027pun), restricting Cas9 and EGFP expression to p21-expressing cells. Mice were co-injected with a second AAV8 vector encoding CAG-driven mCherry together with two U6 promoter-driven single-guide RNAs (sgRNAs) targeting either *Stat1* (Stat1KO group, VectorBuilder VB260406-1226jnu; n = 6) or, as a control, a scramble sequence plus the Hipp11 (H11) safe-harbor locus (VectorBuilder VB260406-1236nds; n = 4). Both vectors were delivered by retro-orbital injection at 2.5 × 10^11^ genome copies per vector per mouse, and livers were collected one month after injection. Snap-frozen whole liver was homogenized on a Precellys tissue disruptor (Bertin Technologies), and total RNA was extracted using the Direct-zol RNA Miniprep kit (Zymo Research) according to the manufacturer’s instructions. RNA integrity was assessed on an Agilent TapeStation; all samples had RIN > 7. Purified RNA was shipped on dry ice to Plasmidsaurus (Louisville, KY), where library preparation and sequencing were performed using their standard 3′ end counting RNA-seq workflow. In brief, mRNA was reverse-transcribed with a poly(dT)VN primer incorporating a unique molecular identifier (UMI) per molecule, and the resulting stranded libraries were sequenced on an Illumina NovaSeq X Plus, yielding mRNA reads of approximately 90 bases enriched near the 3′ end of each transcript.

### *Stat1* knockout RNA-seq analysis

Initial RNA-seq processing was performed by Plasmidsaurus using their standard RNA-seq pipeline. In summary, raw sequencing reads were processed using fastp^124^ v0.24.0 to remove poly-X tails and trim low-quality bases from the 3′ ends, with a minimum Phred quality score of 15 and a minimum retained read length of 25 bp. Read quality was evaluated using FastQC v0.12.1. Filtered reads were aligned to the mouse reference genome GRCm39 (Ensembl release 114) using STAR^125^ v2.7.11b, with noncanonical splice junctions excluded. Alignments were coordinate-sorted using samtools^97^ v1.21, and reads sharing a unique molecular identifier at the same alignment position were collapsed using UMIcollapse^126^ v1.1.0 with the directional algorithm. Alignment quality, strand specificity, and read distribution were assessed using RustQC v0.2.1, and quality-control metrics were aggregated using MultiQC^127^ v1.33. Gene-level counts were generated with featureCounts (Subread v2.1.1)^128^ using strand-specific counting over exon and 3′-UTR features grouped by gene, with a minimum read-overlap fraction of 10%, a minimum mapping quality of 1, and fractional assignment of multi-mapping and multi-overlapping reads. Features corresponding to transgenes were excluded from downstream expression analyses.

Differential gene expression analysis was performed using DESeq2^129^ v1.38.3 in R v4.2.2. Genes with at least 10 counts in a minimum of four samples, corresponding to the smaller experimental group, were retained, yielding 14,592 genes for downstream analysis. Differential expression between *Stat1* knockout (Stat1KO) and control mice was assessed using a generalized linear model with a negative binomial distribution and a design formula of ∼ condition, with Control specified as the reference level. Differential expression was evaluated using the Wald test, and *P* values were adjusted for multiple testing using the Benjamini-Hochberg (BH) procedure. Gene set enrichment analysis (GSEA) was performed using fgsea^130^ v1.24.0, with genes ranked by their DESeq2 Wald statistics and 10,000 simple permutations (nPermSimple = 10000). Gene sets were obtained from the mouse Molecular Signatures Database (MSigDB)^131^ using msigdbr v25.1.1.

Gene set activity was assessed using GSVA^73^ (v1.50.5) in R 4.3.3 on variance-stabilized bulk RNA-seq expression data from 10 mice (Stat1KO, n = 6; Control, n = 4). The analysis included 14,592 genes with ≥10 counts in ≥4 samples, using a Gaussian kernel (kcdf = “Gaussian”) and default settings otherwise. Three categories of gene sets were analyzed: (1) 37 hepatocyte STAT1 target genes identified from the 10x Multiome gene regulatory network; (2) four hepatocyte senotype signatures (S1–S4); and (3) senescence-associated genes upregulated (453 genes) or downregulated (99 genes) in senescent versus non-senescent hepatocytes, identified from 10x Multiome data (adjusted *P* < 0.05, |log₂FC| > 0.5). GSVA scores were compared between Stat1KO and control mice using two-sided exact Wilcoxon rank-sum tests.

### Lamina-associated domain (LAD) calling

LADs were called from a previously published DamID dataset in wild-type mouse cells^69^ (GEO: GSE151044), by converting a bigwig file to bedgraph using bigWigToBedGraph from UCSC tools. Next the bedgraph was converted from mm9 to mm10 genome using UCSC liftOver. Using the processed, smoothed, bedGraph track, genomic bins with a DamID score greater than 1.0 were retained, sorted by coordinate, and merged with BEDTools merge (v2.31.1), joining intervals separated by 50 kb or less. Merged intervals shorter than 200 kb were discarded, and the remaining intervals were taken as LADs (n = 1,074 domains).

### Immunofluorescence

Paraffin liver sections were stained with antibodies against STAT1 (Cell Signaling Technology, cat# 9172S) and GFP (Abcam, cat# ab13970), followed by Alexa Fluor 647- and Alexa Fluor 488-conjugated secondary antibodies, and counterstained with DAPI. Images were acquired on a Nikon Ti2 microscope. GFP+, STAT1+ and double-positive cells were counted manually in ImageJ (Fiji)^132^ by an experimenter blinded to treatment, in three to four fields per mouse, and knockout efficiency was expressed as the percentage of GFP+ cells that were also STAT1+. Fields were averaged to give one value per mouse, and groups were compared with a two-sided Welch’s unpaired t-test (sgH11, n = 4; sgStat1, n = 6 mice).

### Genomic DNA isolation and Sanger sequencing

The sgRNA sequences were: sgStat1_g1, 5′-GGTACTGTCTGATTTCCATG-3′; sgStat1_g2, 5′-GAGGAGGTCATGGAAGCGGA-3′; sgH11 (control), 5′-GTCATGAGACTCATACTACG-3′. Genomic DNA was extracted from whole liver using the Monarch Genomic DNA Purification kit (New England Biolabs, ref T3010S), followed by PCR amplification of the 389 bp region surrounding the sgStat1_g2 target site with 2X Taq PCR Master Mix (APExBIO, ref K1034) and primers forward 5′-TGACATAGAAAGCTCAGACCCC-3′ and reverse 5′-AGGTTTTAAGTCAGCGGAGACA-3′ at an annealing temperature of 60 °C for 35 cycles. Products were purified with the QIAquick PCR Purification Kit (QIAGEN, ref 28104) and Sanger sequenced with the forward amplification primer (Genewiz).

## Data availability

Mouse 10x Multiome, Paired-Tag, snm3C and MERFISH datasets are available through the SenNet Data Portal (https://sennetconsortium.org/), with corresponding dataset IDs provided in Supplementary Table 1.

## Code availability

Custom code for data processing and analysis in this study is available on Github at https://github.com/sainathmamde/Spatial-and-epigenetic-programs-underlie-heterogeneous-cell ular-senescence-in-the-aging-liver.

---

**Supplementary Fig. 1.**
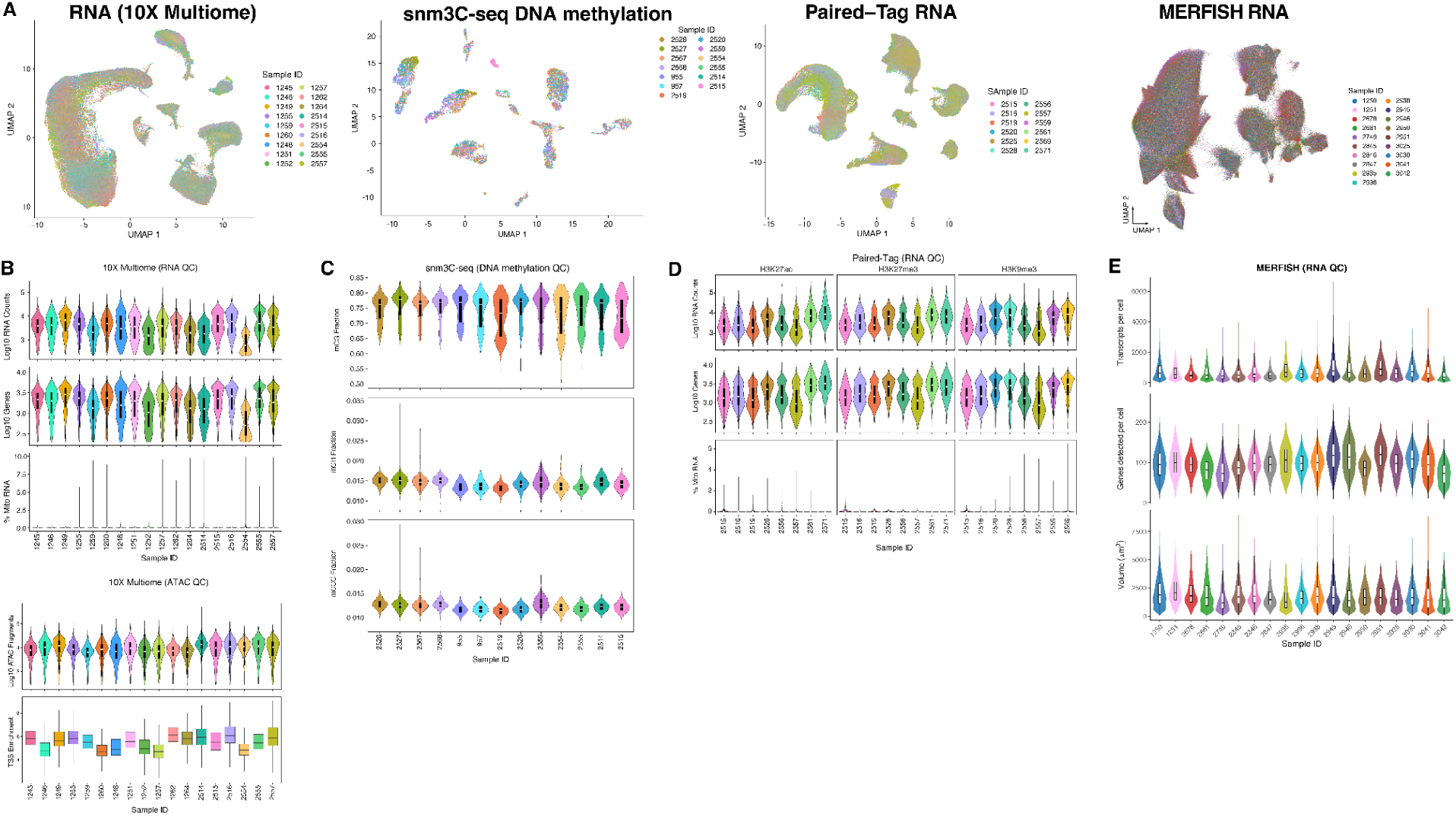
10x Multiome, snm3C, Paired-Tag and MERFISH quality metrics across samples. **A**, UMAP embeddings of 10x multiome RNA, snm3C-seq DNA methylation, Paired-Tag RNA and MERFISH RNA clustering, colored by mouse. **B**–**E**, Violin plots and boxplots showing the distribution of quality metrics in 10x Multiome RNA and ATAC (**B**), snm3C DNA methylation (**C**), Paired-Tag RNA (**D**) and MERFISH RNA (**E**) across mouse samples.

**Supplementary Fig. 2.**
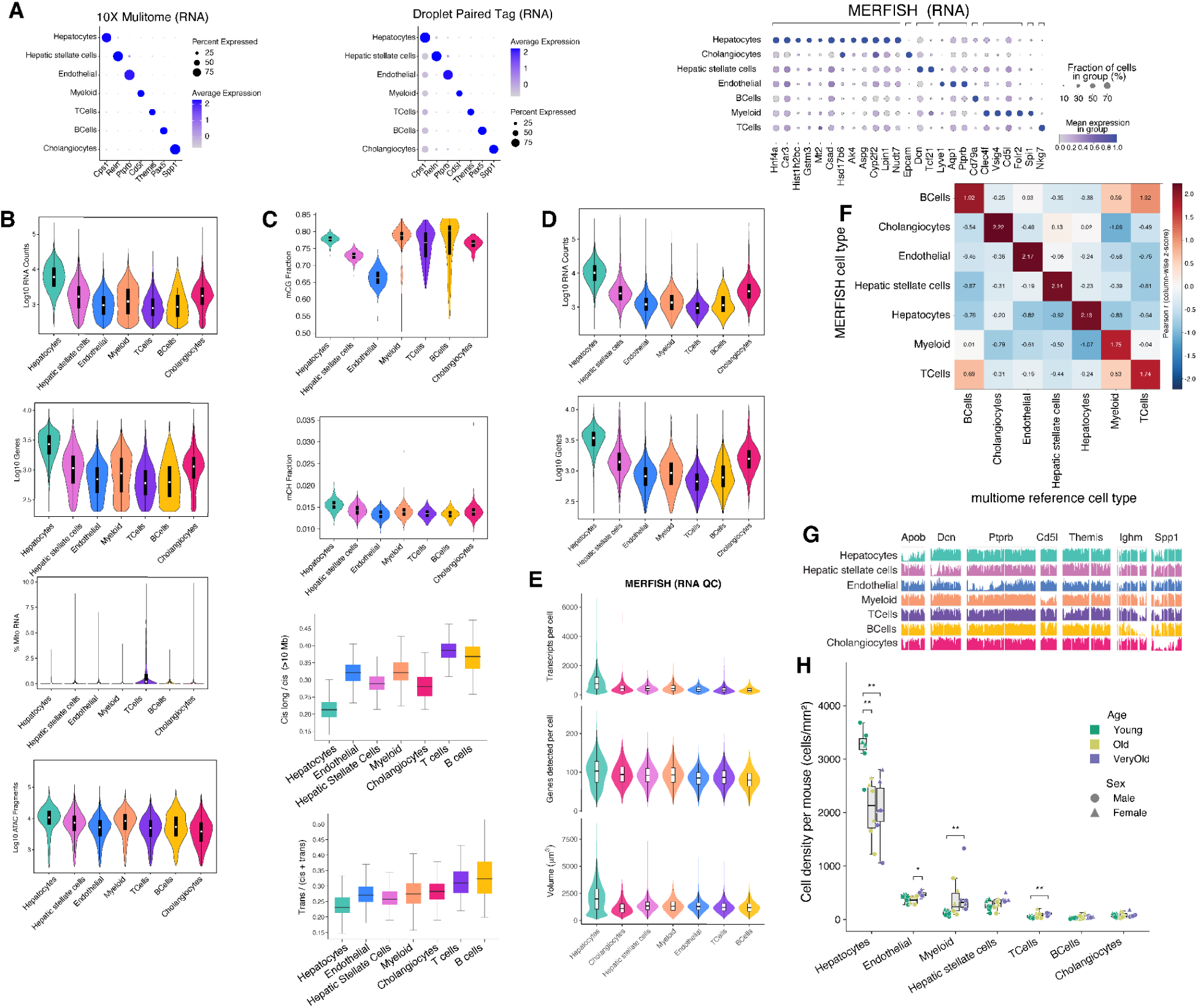
Single cell annotations and feature metrics. **A**, Dot plots showing the expression of cell type marker genes across cell types, for 10x Multiome RNA, Paired-Tag RNA and MERFISH RNA. Dot size, percentage of cells expressing the genes; dot color, average expression level. **B**–**E**, Violin plots and boxplots showing the distribution of quality metrics in 10x Multiome RNA and ATAC (**B**), snm3C DNA methylation and chromatin contacts (**C**), Paired-Tag RNA (**D**) and MERFISH RNA (**E**) across cell types. **F**, Heatmap showing Pearson correlation coefficients between cell-type-averaged gene expression profiles from MERFISH and the 10x Multiome reference. **G**, mCG fraction in cell type marker gene regions at the pseudo bulk level for each cell type. **H**, Density of each cell type in young, old and very old mice. Points, individual mice; box, median and IQR; whiskers, 1.5 × IQR. *P < 0.05, **P < 0.01, Wilcoxon rank-sum test.

**Supplementary Fig. 3.**
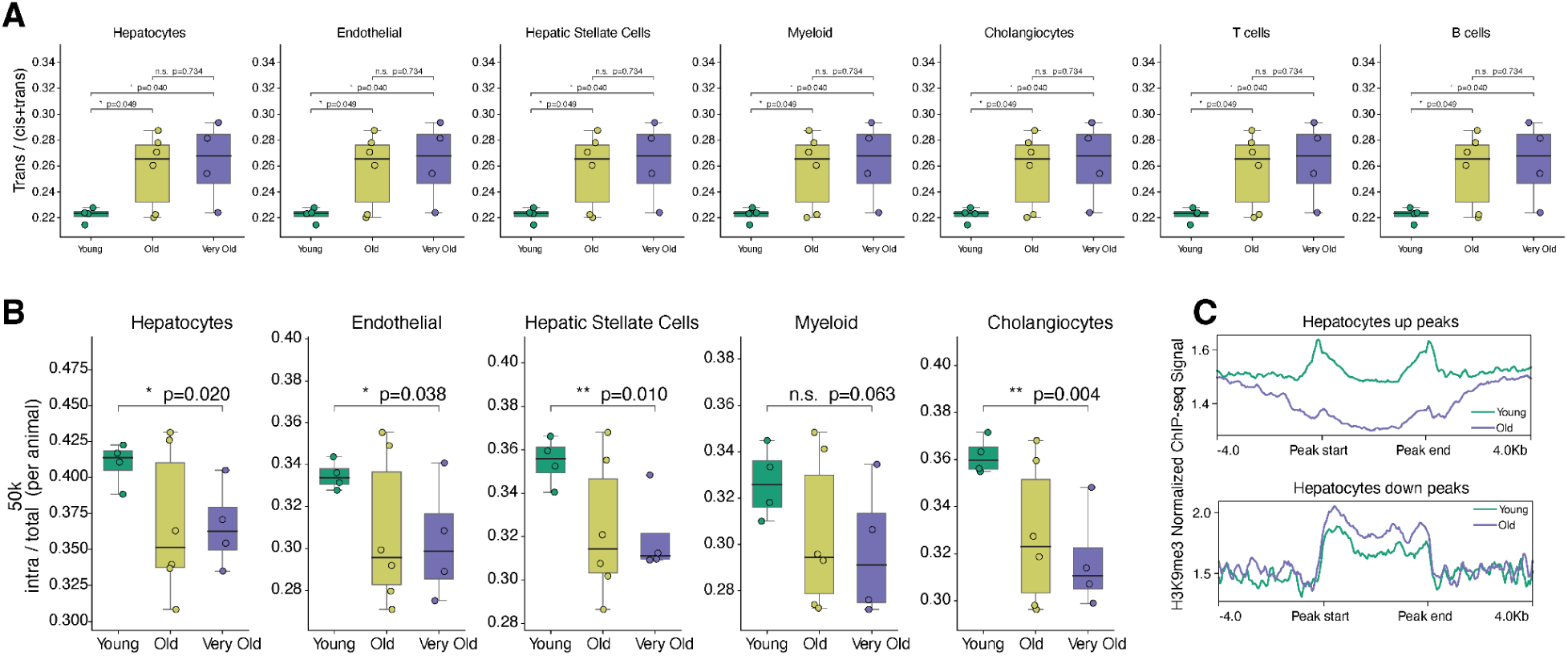
Age-associated loss of 3D chromatin organization and epigenetic information across cell types during aging. **A**, Fraction of contacts that were trans-chromosomal in each cell type, by age group. Points, individual mice; box, median and IQR; whiskers, 1.5 × IQR. P values, t-test; n.s., not significant. **B**, Fraction of contacts falling within TADs in each cell type, by age group; plotted and tested as in **A**. **C**, Mean H3K9me3 signal from ChIP-seq^132^ over domains gained (left) and lost (right) with age in hepatocytes identified in our Paired-Tag data, extended 2 kb either side, for each age group.

**Supplementary Fig. 4.**
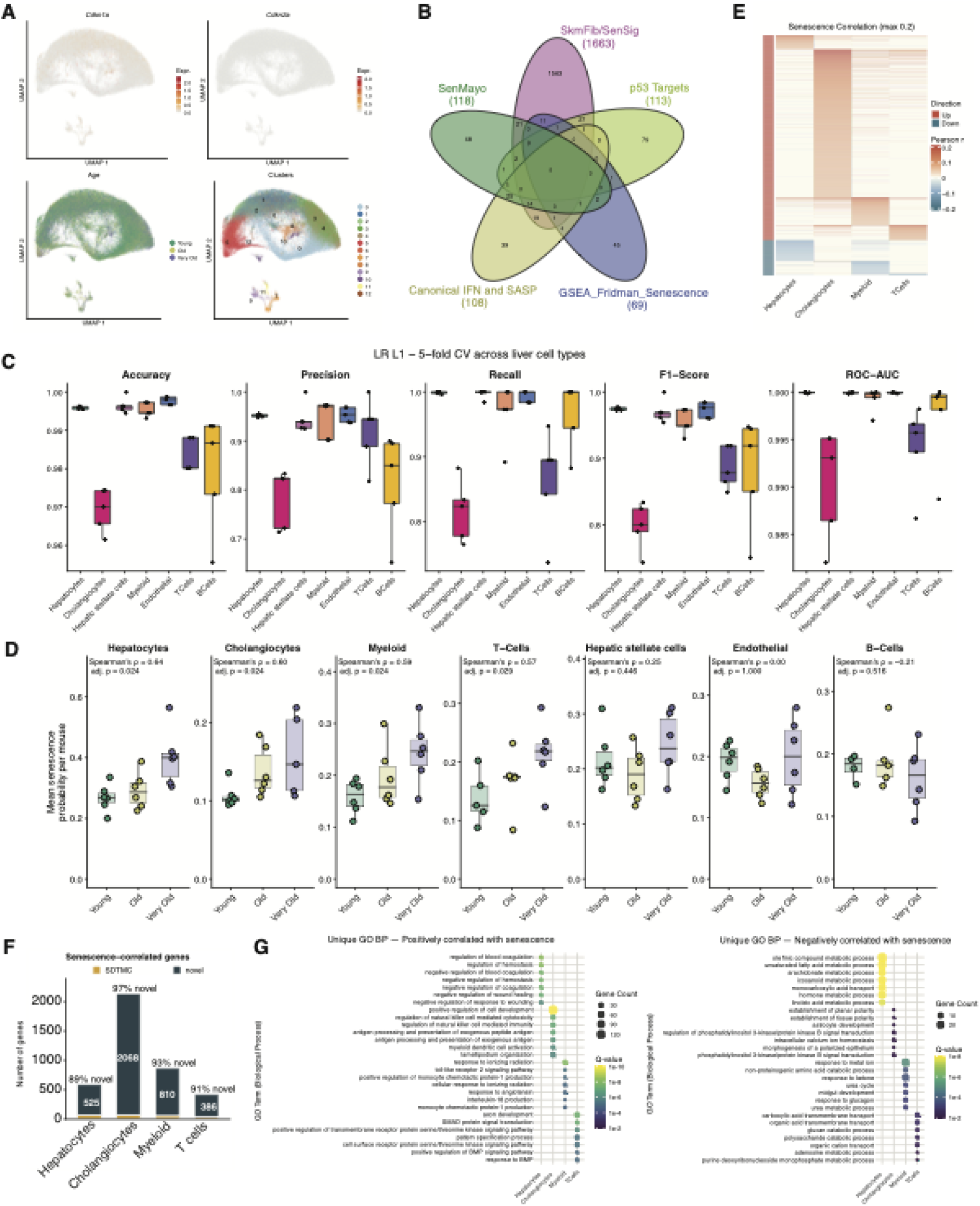
Identification of SnCs and senescence-associated genes using SenLearner. **A**, UMAP plots of hepatocytes colored by expression of *Cdkn1a* (top left) and *Cdkn2a* (top right), age (bottom left) and clusters (bottom right). **B**, Venn diagram showing the overlaps among five published gene lists. **C**, Performance metrics of L1-regularized logistic regression models across cell types, evaluated using fivefold cross-validation. Points, individual folds; box, median and IQR; whiskers, 1.5 × IQR. **D**, Mean SenLearner probability per donor in each cell type, by age group. Points, individual mice; box, median and IQR; whiskers, 1.5 × IQR. Spearman correlation between per-mouse mean SenLearner probability and age (two-sided test); P values were adjusted across the seven cell types using the Benjamini–Hochberg method. **E**, Heatmap showing Pearson correlation between gene expression and SenLearner probability for the genes positively (Up) or negatively (Down) correlated with senescence in any of the four cell types (|r| > 0.05, adjusted P < 0.01). **F**, Number and proportion of novel senescence-correlated genes in each cell type. **G**, Cell type-specific gene ontology biological processes uniquely enriched among genes positively (left) and negatively (right) correlated with senescence probability. Dot size, number of genes; color, adjusted q value.

**Supplementary Fig. 5.**
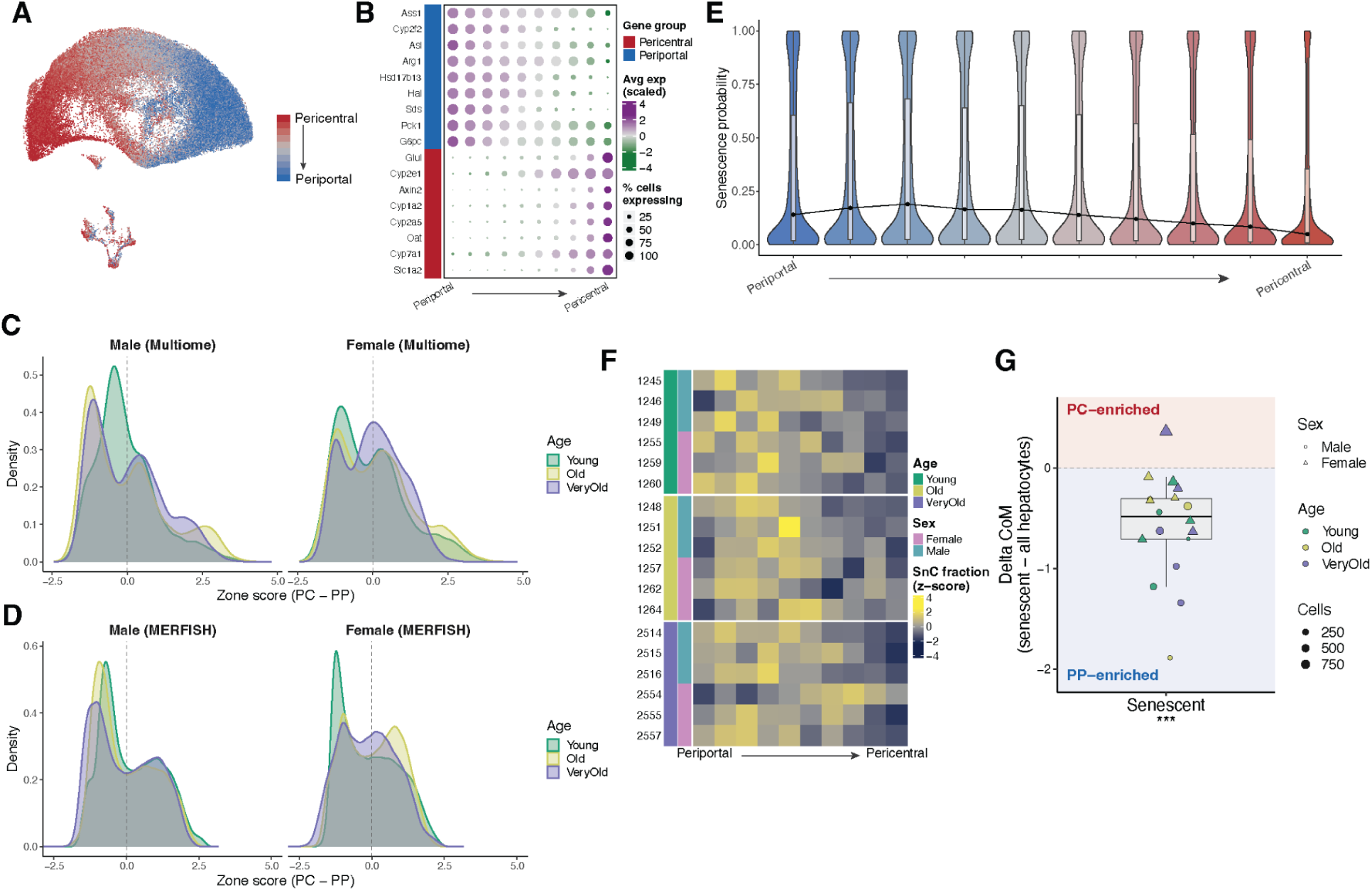
Periportal enrichment of senescent hepatocytes. **A**, UMAP embedding showing hepatocyte across 10 zone bins from periportal to pericentral bins. **B**, Zonation marker gene expression across 10 zone bins from periportal to pericentral bins. **C**–**D**, Distribution of zone score, measured by difference between pericentral and periportal signature scores, by age group, separated by sex, in 10x Multiome (**C**) and MERFISH (**D**). **E**, Violin plots showing senescence probability of all hepatocytes across 10 zone bins. **F**, Heatmap showing the proportion of senescent hepatocytes in each zone bin and in each mouse. **G**, Difference in the centre of mass of the zonal distribution between senescent hepatocytes and all hepatocytes; positive values indicate pericentral and negative values periportal enrichment. Points, individual mice; color, age group; shape, sex; size, number of cells. ***P < 0.001, cell-count-weighted permutation test.

**Supplementary Fig. 6.**
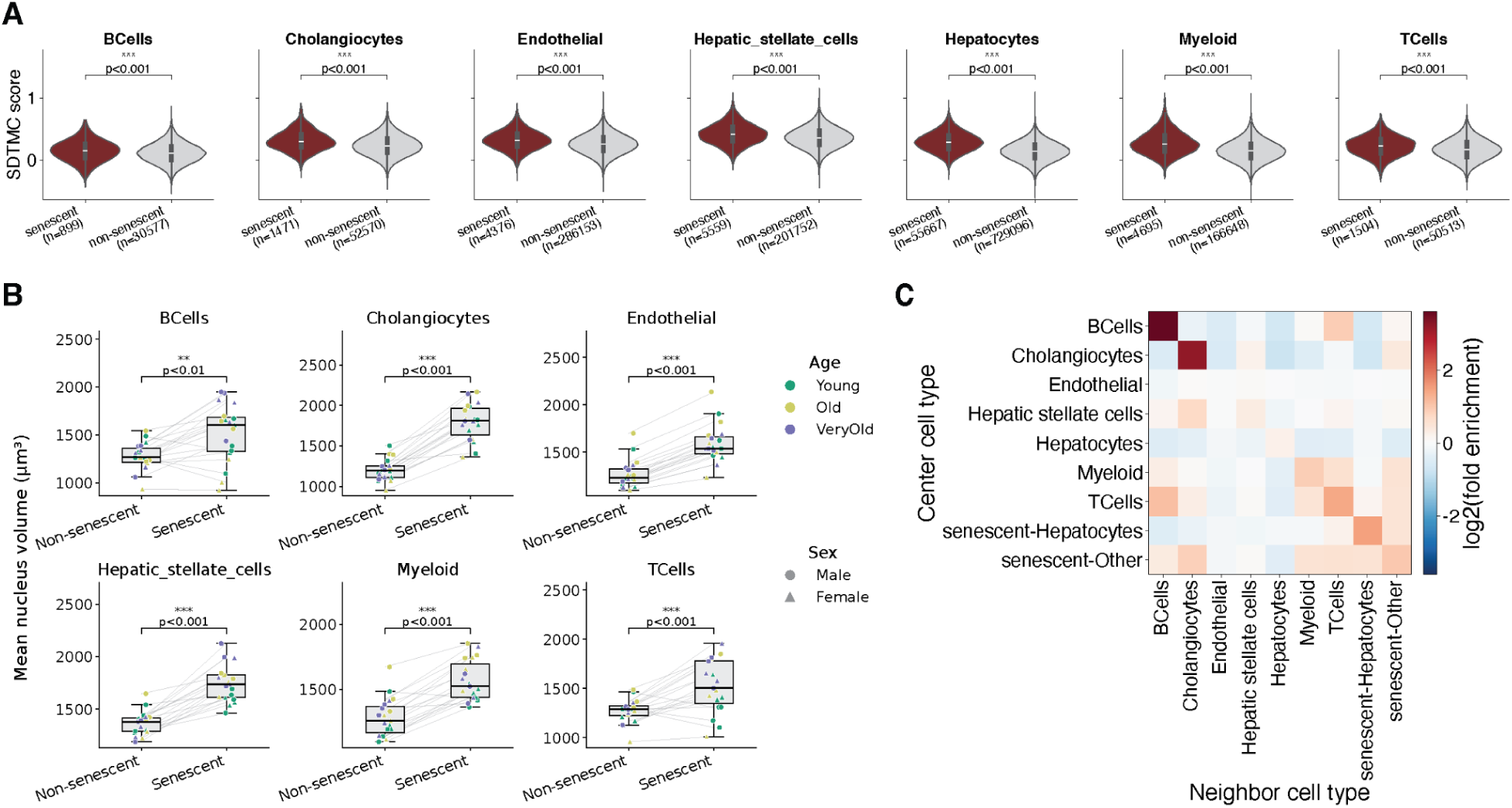
Senescent cells mapped from 10x Multiome onto MERFISH. **A**, Violin plots comparing SDTMC score between senescent and non-senescent cells in each cell type. P values, Wilcoxon rank-sum tests. **B**, Mean nuclear volume of senescent and non-senescent cells in each cell type. Points, individual mice (n = 19 mice); lines connect paired measurements in each mouse. P-values, paired Wilcoxon rank-sum tests. **C**, Heatmap of cell-type neighborhood enrichment across 25 MERFISH sections from 19 mice. Senescent cells were classified as senescent hepatocytes (senescent-Hepatocytes) or other senescent cell types (senescent-Other). For each center cell type (rows), neighboring cells (columns) within a 50 µm radius were counted, excluding the center cell. Enrichment was calculated as the fraction of neighbors of each type relative to its overall abundance across all sections.

**Supplementary Fig. 7.**
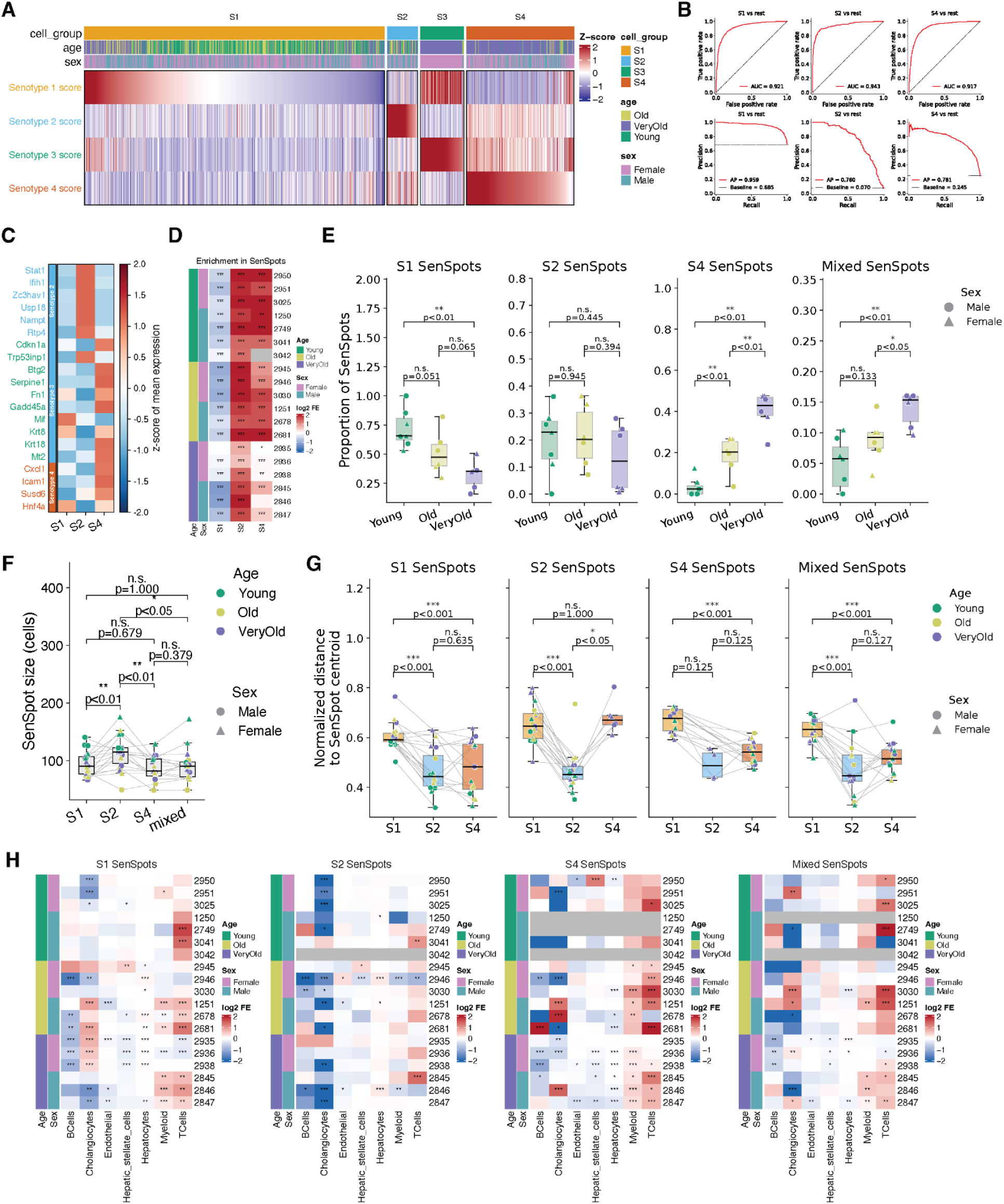
Hepatocyte senotypes mapped from 10x Multiome onto MERFISH. **A**, Heatmap of four senotype signature scores (rows) across 6,342 senescent hepatocytes (columns), grouped by senotype cell groups and ordered by decreasing corresponding senotype score within each group. Top annotations indicate senotype cell group, age, and sex. **B**, Performance of a multinomial logistic regression classifier trained on SDTMC genes to distinguish senotypes in senescent hepatocytes, excluding S3 due to its predominance in a single mouse. Receiver operating characteristic (ROC) (top) and precision-recall (bottom) curves for each senotype versus the others, evaluated using stratified fivefold cross-validation. Values indicate area under the ROC curve (top) or average precision (bottom); dashed lines indicate chance-level performance. **C**, Expression of senotype genes in the senotype cells mapped from 10x Multiome onto MERFISH. **D**, Enrichment of each senotype within SenSpots relative to all senescent hepatocytes, for 19 mice ordered by age group and annotated by sex. Color, log2 fold enrichment. *P < 0.05, **P < 0.01, ***P < 0.001, test. **E**, Proportion of SenSpot types, defined by dominant senotype hepatocytes, by age group. Points, individual mice; point shape, sex; box, median and IQR; whiskers, 1.5 × IQR. P values, Wilcoxon rank-sum tests. **F**, SenSpot size (number of cells) by dominant hepatocyte senotype. Points, individual mice; color, age; shape, sex; box, median and IQR; whiskers, 1.5 × IQR. Lines connect paired measurements. P values, paired Wilcoxon rank-sum tests. **G**, Relative distance of senescent hepatocytes from SenSpot centroids, stratified by SenSpot type (S1, S2, S4, or mixed) and individual cell senotype. Distances were normalized to the maximum centroid-to-cell distance within each SenSpot (0 = center, 1 = edge) and averaged per mouse. Points, individual mice; color, age; shape, sex; box, median and IQR; whiskers, 1.5 × IQR. Lines connect paired measurements. P values, paired Wilcoxon rank-sum tests. **H**, Enrichment of each cell type within senotype SenSpots relative to all senescent hepatocytes, for 19 mice ordered by age and sex. Color, log2 fold enrichment. *q < 0.05, **q < 0.01, ***q < 0.001, one-sided Fisher’s exact test with Benjamini-Hochberg FDR correction.

**Supplementary Fig. 8.**
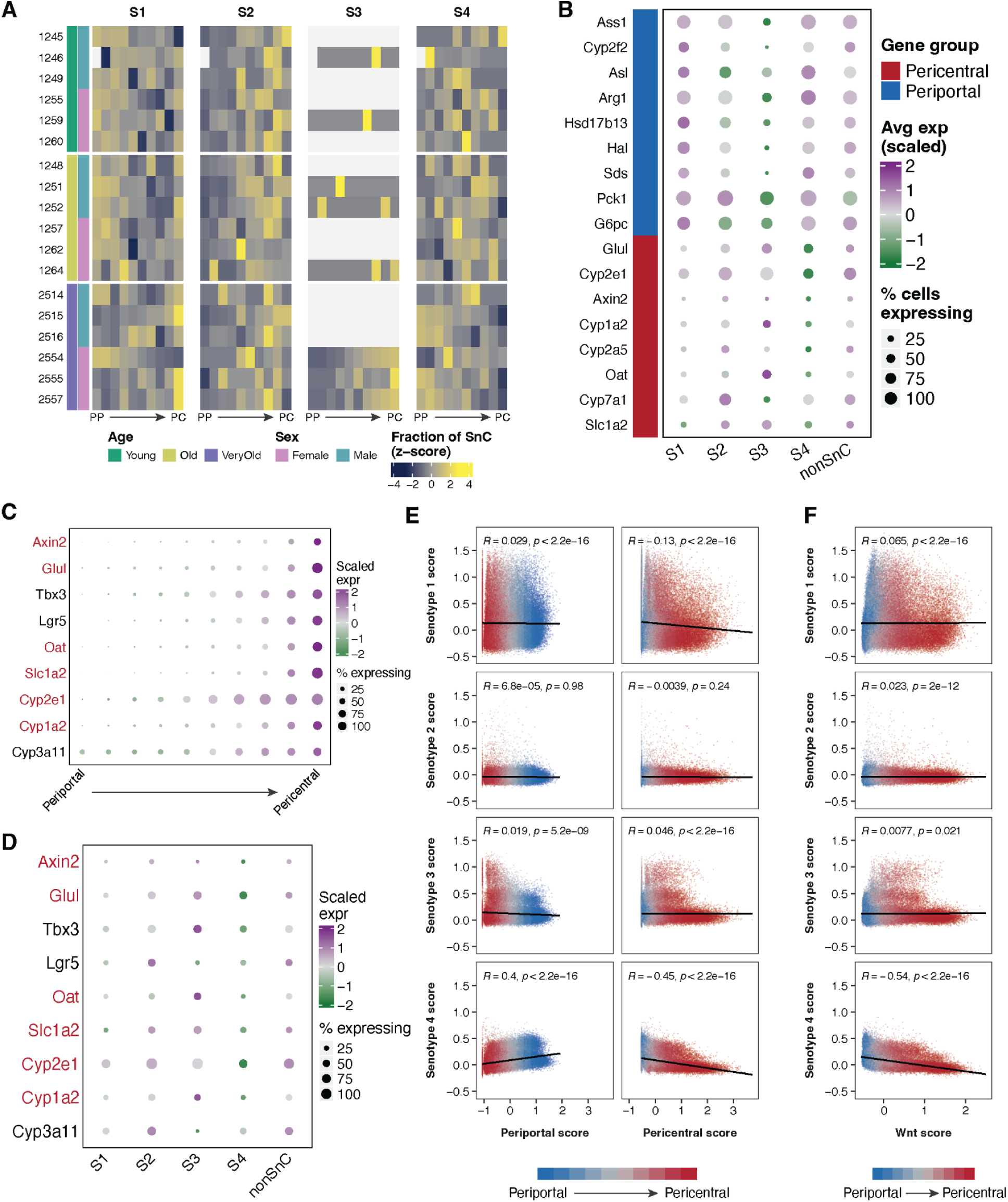
Zone specificity of hepatocyte senotypes. **A**, Heatmap showing the proportion of senotype hepatocytes among senescent hepatocytes in each zone bin and in each mouse. **B**, Dot plot showing the expression of zonation marker genes across senotype hepatocytes and non-senescent hepatocytes. Dot size, percentage of cells expressing the genes; dot color, average expression level. **C**, Dot plot showing the expression of Wnt target genes across 10 hepatocyte zone bins. Dot size, percentage of cells expressing the genes; dot color, average expression level. Genes in red, pericentral marker genes. **D**, Dot plot showing the expression of Wnt target genes across senotype hepatocytes and non-senescent hepatocytes. Plotted as in **C**. **E**–**F**, Scatter plots showing associations of senotype sores with periportal and pericentral scores (**E**) or Wnt target gene scores (**F**) across all hepatocytes. Points, individual cells colored by zone bin; black lines, linear regression fits with 95% confidence intervals. Spearman’s ρ and *P* values are indicated in each panel.

**Supplementary Fig. 9.**
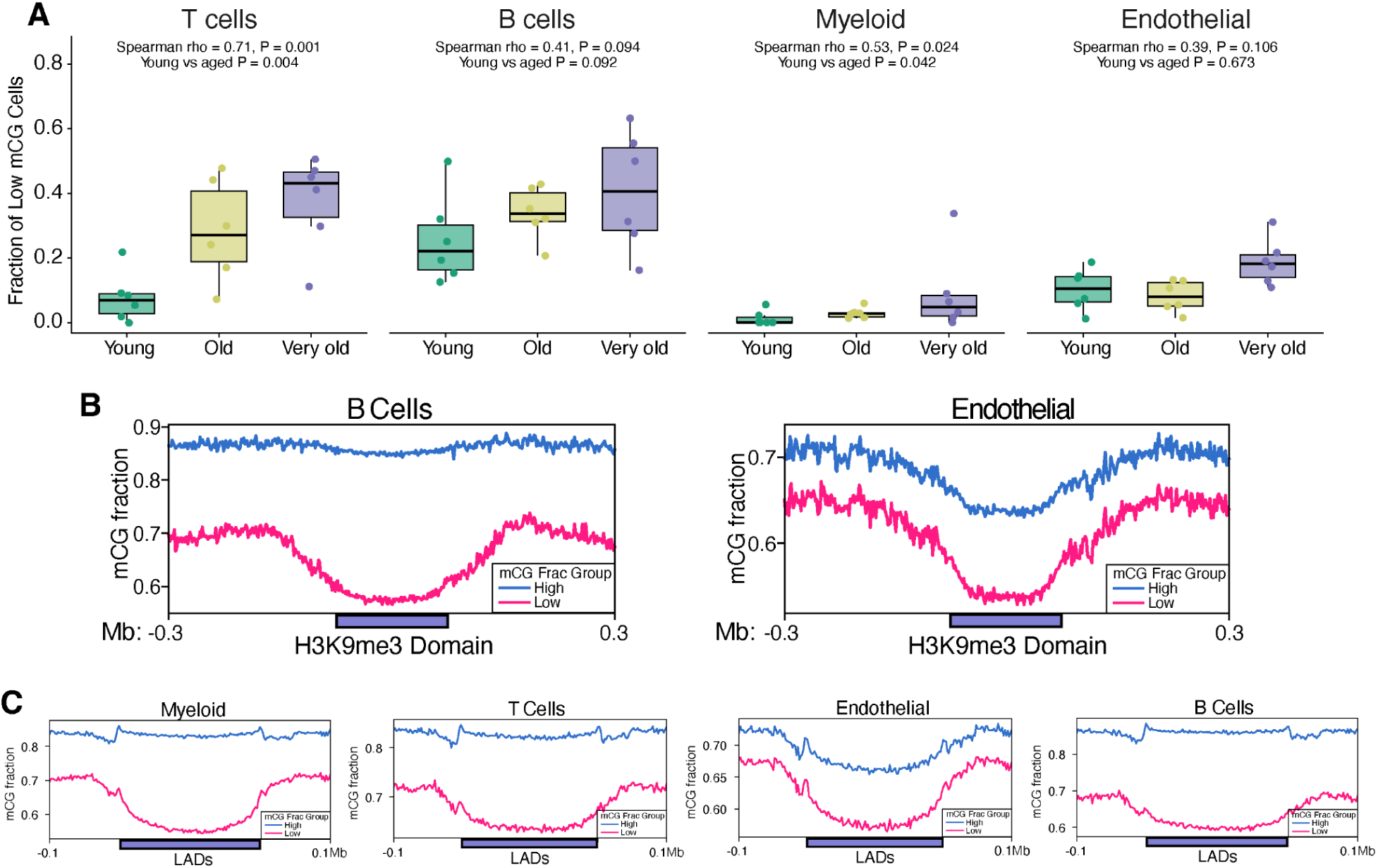
Age-associated DNA methylation loss in constitutive heterochromatin. **A**, Box plots of proportions of cells with globally low-mCG across age groups. Each dot represents an animal. Box, median and IQR; whiskers, 1.5 × IQR; points, individual mice. Two-sided Spearman correlation with age in months. **B**, mCG fraction of high- and low-mCG nuclei across cell type-specific H3K9me3 domains, scaled to a common length, with 0.3-Mb flanking regions, in B cells (n = 477 domains; left) and endothelial cells (n = 418 domains; right). **C**, mCG fraction of high- and low-mCG nuclei across mouse liver LADs (PMID: 33330474; n = 1,074) scaled to a common length, with 0.1-Mb flanking regions.

**Supplementary Fig. 10.**
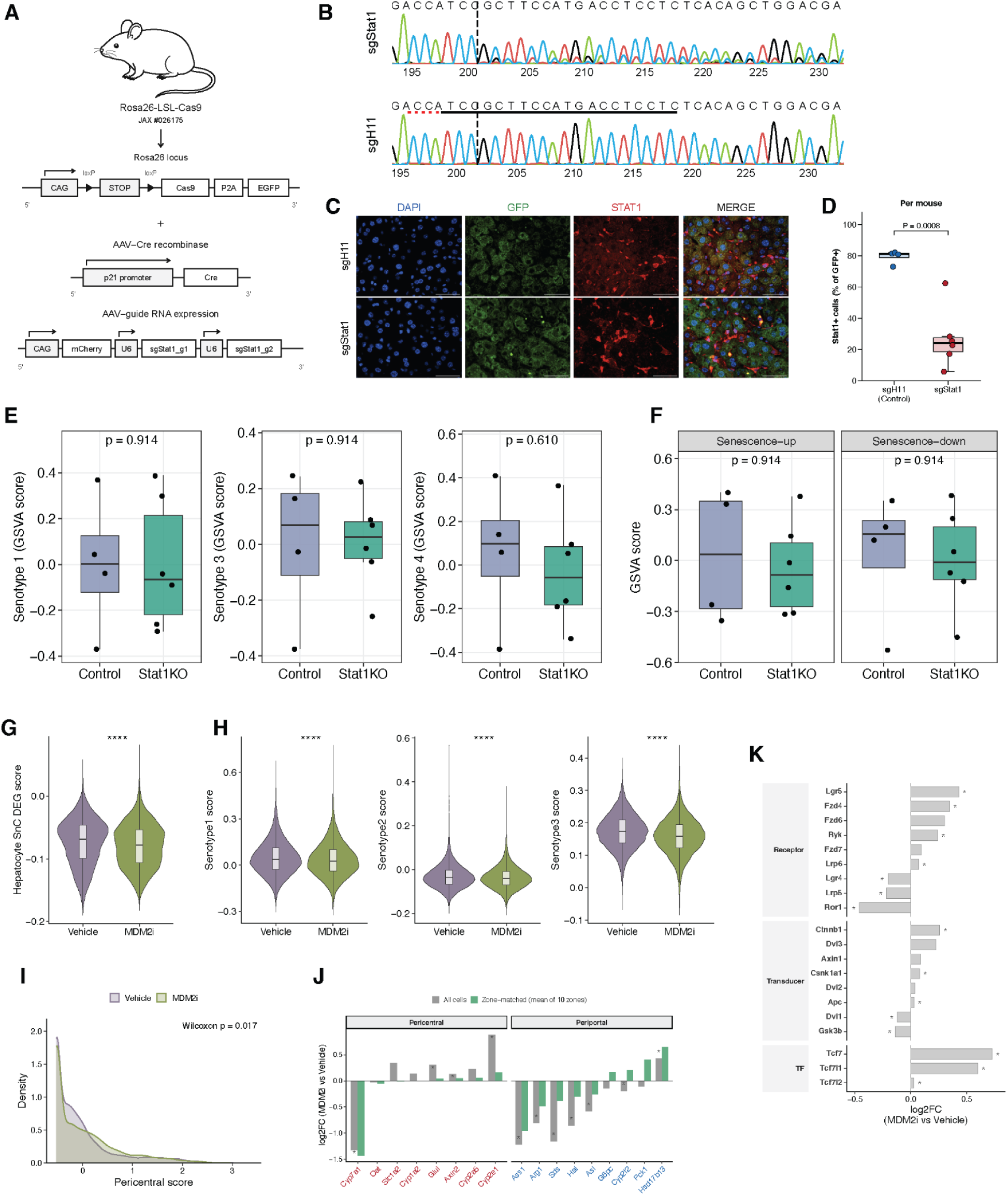
Senotype-specific effect of *Stat1* knockout and MDM2 inhibition. **A**, Schematic representation of the approach used to delete Stat1 in p21-expressing cells. Aged mice carrying a Cre-dependent Rosa26-CAG-loxP-STOP-loxP-Cas9-P2A-EGFP allele (JAX #026175) were treated with an AAV8 expressing Cre under the control of the p21 promoter, together with an AAV expressing mCherry and two U6 promoter-driven sgRNAs targeting either Stat1 (sgStat1) or the H11 safe-harbor locus (sgH11, control). **B**, Representative Sanger sequencing traces of gDNA isolated from the whole liver of sgStat1- and sgH11-treated mice at the sgStat1_g2 locus. Dashed line indicates the predicted Cas9 cut site. **C**, Representative immunofluorescence images of liver sections from sgH11 (control) and sgStat1 mice stained for DAPI (blue), GFP (green) and STAT1 (red), and the merged channels. GFP marks Cre-recombined, p21-expressing cells. Scale bar, 50 µm. **D**, Quantification of STAT1+ cells as a percentage of GFP+ cells from **C** (sgH11, n = 4; sgStat1, n = 6 mice). Each point represents one mouse and is the mean of 3–4 fields. P = 0.0008. (**D**) Statistical significance was assessed with a two-sided Welch’s unpaired t-test on per-mouse values. Box shows the 25th–75th percentile, center line shows the median, whiskers extend to 1.5 × IQR. ***P < 0.001. **E**, Gene set variation analysis (GSVA) scores for senotype 1, senotype 3 and senotype 4 signatures in control and Stat1KO liver. Points, individual mice; box, median and IQR; whiskers, 1.5 × IQR. Wilcoxon rank-sum test. **F**, GSVA scores for genes up- or down-regulated in senescent hepatocytes in control and Stat1KO liver. Plotted and tested as in **E**. **G**, Module scores of senescence-associated differentially expressed genes identified in wild-type hepatocytes, evaluated in hepatocytes from vehicle- and MDM2i-treated mice. Violin, distribution across nuclei; box, median and IQR. ****P < 0.0001, Wilcoxon rank-sum test. n = 2 mice per group. **H**, Module scores of senotype 1, senotype 2 and senotype 3 genes, evaluated in hepatocytes from vehicle- and MDM2i-treated mice. Plotted and tested as in G. **I**, Distribution of pericentral signature scores in hepatocytes from vehicle- and MDM2i-treated mice. Wilcoxon rank-sum test. **J**, Log2 fold change (MDM2i versus vehicle) of pericentral (left) and periportal (right) marker genes across all hepatocytes (grey) or averaged across 10 zonation bins (green). Asterisks indicate consistent directional changes across both mice per group. **K**, Log2 fold change (MDM2i versus vehicle) in hepatocyte expression of upstream Wnt pathway components, grouped into Wnt receptors/co-receptors, cytoplasmic signal transducers, and TCF/LEF transcription factors. Genes are ordered by log2 fold change within each group. *adjusted P < 0.05.

## Notes

### Competing Interest Statement

Bing Ren has equity interests in the following companies: Arima Genomics, Inc. Epigenome Technologies, Inc.

### Funding

- NIH Common Fund: U54 AG079758

## References

1. Bochkis, I. M., Przybylski, D., Chen, J. & Regev, A. Changes in nucleosome occupancy associated with metabolic alterations in aged mammalian liver. Cell Rep. 9, 996–1006 (2014).

2. Bozukova, M. et al. Aging is associated with increased chromatin accessibility and reduced polymerase pausing in liver. Mol. Syst. Biol. 18, e11002 (2022).

3. Tyshkovskiy, A. et al. Universal transcriptomic hallmarks of mammalian ageing and mortality. Nature 654, 173–188 (2026).

4. López-Otín, C., Blasco, M. A., Partridge, L., Serrano, M. & Kroemer, G. Hallmarks of aging: An expanding universe. Cell 186, 243–278 (2023).

5. Gorgoulis, V. et al. Cellular senescence: defining a path forward. Cell 179, 813–827 (2019).

6. Serrano, M., Lin, A. W., McCurrach, M. E., Beach, D. & Lowe, S. W. Oncogenic ras provokes premature cell senescence associated with accumulation of p53 and p16INK4a. Cell 88, 593–602 (1997).

7. Noda, A., Ning, Y., Venable, S. F., Pereira-Smith, O. M. & Smith, J. R. Cloning of senescent cell-derived inhibitors of DNA synthesis using an expression screen. Exp. Cell Res. 211, 90–98 (1994).

8. Coppé, J.-P. et al. Senescence-associated secretory phenotypes reveal cell-nonautonomous functions of oncogenic RAS and the p53 tumor suppressor. PLoS Biol. 6, e301 (2008).

9. Acosta, J. C. et al. Chemokine signaling via the CXCR2 receptor reinforces senescence. Cell 133, 1006–1018 (2008).

10. Kuilman, T. et al. Oncogene-induced senescence relayed by an interleukin-dependent inflammatory network. Cell 133, 1019–1031 (2008).

11. Kang, T.-W. et al. Senescence surveillance of pre-malignant hepatocytes limits liver cancer development. Nature 479, 547–551 (2011).

12. Baker, D. J. et al. Naturally occurring p16Ink4a-positive cells shorten healthy lifespan. Nature 530, 184–189 (2016).

13. De Cecco, M. et al. L1 drives IFN in senescent cells and promotes age-associated inflammation. Nature 566, 73–78 (2019).

14. McHugh, D., Durán, I. & Gil, J. Senescence as a therapeutic target in cancer and age-related diseases. Nat. Rev. Drug Discov. 24, 57–71 (2025).

15. Chang, J. et al. Clearance of senescent cells by ABT263 rejuvenates aged hematopoietic stem cells in mice. Nat. Med. 22, 78–83 (2016).

16. Xu, M. et al. Senolytics improve physical function and increase lifespan in old age. Nat. Med. 24, 1246–1256 (2018).

17. Halpern, K. B. et al. Single-cell spatial reconstruction reveals global division of labour in the mammalian liver. Nature 542, 352–356 (2017).

18. Ogrodnik, M. et al. Cellular senescence drives age-dependent hepatic steatosis. Nat. Commun. 8, 15691 (2017).

19. Mohammed, S. et al. Necroptosis contributes to chronic inflammation and fibrosis in aging liver. Aging Cell 20, e13512 (2021).

20. Nikopoulou, C. et al. Spatial and single-cell profiling of the metabolome, transcriptome and epigenome of the aging mouse liver. Nat. Aging 3, 1430–1445 (2023).

21. Laux, L. et al. Cellular senescence is associated with age-related loss of liver zonation and hepatocyte function. Preprint at bioRxiv 10.64898/2026.08.08.743614 (2026).

22. Karpova, A. et al. Cellular senescence in human liver under normal aging and cancer. Cell Genom. 6, 101133 (2026).

23. Tabula Muris Consortium. A single-cell transcriptomic atlas characterizes ageing tissues in the mouse. Nature 583, 590–595 (2020).

24. Krizhanovsky, V. et al. Senescence of activated stellate cells limits liver fibrosis. Cell 134, 657–667 (2008).

25. Grosse, L. et al. Defined p16High senescent cell types are indispensable for mouse healthspan. Cell Metab. 32, 87–99.e6 (2020).

26. Suryadevara, V. et al. SenNet recommendations for detecting senescent cells in different tissues. Nat. Rev. Mol. Cell Biol. 25, 1001–1023 (2024).

27. Ogrodnik, M. et al. Guidelines for minimal information on cellular senescence experimentation in vivo. Cell 187, 4150–4175 (2024).

28. Hall, B. M. et al. p16(Ink4a) and senescence-associated β-galactosidase can be induced in macrophages as part of a reversible response to physiological stimuli. Aging (Albany NY) 9, 1867–1884 (2017).

29. Freund, A., Laberge, R.-M., Demaria, M. & Campisi, J. Lamin B1 loss is a senescence-associated biomarker. Mol. Biol. Cell 23, 2066–2075 (2012).

30. Gurkar, A. U. et al. Spatial mapping of cellular senescence: emerging challenges and opportunities. Nat. Aging 3, 776–790 (2023).

31. Chandra, T. et al. Global reorganization of the nuclear landscape in senescent cells. Cell Rep. 10, 471–483 (2015).

32. Chandra, T. et al. Independence of repressive histone marks and chromatin compaction during senescent heterochromatic layer formation. Mol. Cell 47, 203–214 (2012).

33. Narita, M. et al. Rb-mediated heterochromatin formation and silencing of E2F target genes during cellular senescence. Cell 113, 703–716 (2003).

34. Sati, S. et al. 4D genome rewiring during oncogene-induced and replicative senescence. Mol. Cell 78, 522–538.e9 (2020).

35. Saul, D. et al. A new gene set identifies senescent cells and predicts senescence-associated pathways across tissues. Nat. Commun. 13, 4827 (2022).

36. Fridman, A. L. & Tainsky, M. A. Critical pathways in cellular senescence and immortalization revealed by gene expression profiling. Oncogene 27, 5975–5987 (2008).

37. Cherry, C. et al. Transfer learning in a biomaterial fibrosis model identifies in vivo senescence heterogeneity and contributions to vascularization and matrix production across species and diverse pathologies. GeroScience 45, 2559–2587 (2023).

38. Fischer, M. Census and evaluation of p53 target genes. Oncogene 36, 3943–3956 (2017).

39. Robinson, M. D., McCarthy, D. J. & Smyth, G. K. edgeR: a Bioconductor package for differential expression analysis of digital gene expression data. Bioinformatics 26, 139–140 (2010).

40. Poisson, J. et al. Liver sinusoidal endothelial cells: Physiology and role in liver diseases. J. Hepatol. 66, 212–227 (2017).

41. Sørensen, K. K., Simon-Santamaria, J., McCuskey, R. S. & Smedsrød, B. Liver sinusoidal endothelial cells. Compr. Physiol. 5, 1751–1774 (2015).

42. Kostallari, E. & Shah, V. H. Pericytes in the liver. in Pericyte Biology in Different Organs (ed. Birbrair, A.) vol. 1122 153–167 (Springer International Publishing, Cham, 2019).

43. Schwabe, R. F. & Brenner, D. A. Hepatic stellate cells: balancing homeostasis, hepatoprotection and fibrogenesis in health and disease. Nat. Rev. Gastroenterol. Hepatol. 22, 481–499 (2025).

44. Maeso-Díaz, R., et al. Effects of aging on liver microcirculatory function and sinusoidal phenotype. Aging Cell 17, e12829 (2018).

45. Franceschi, C., Garagnani, P., Parini, P., Giuliani, C. & Santoro, A. Inflammaging: a new immune–metabolic viewpoint for age-related diseases. Nat. Rev. Endocrinol. 14, 576–590 (2018).

46. Fulco, C. P. et al. Activity-by-contact model of enhancer–promoter regulation from thousands of CRISPR perturbations. Nat. Genet. 51, 1664–1669 (2019).

47. Mannaerts, I. et al. The Hippo pathway effector YAP controls mouse hepatic stellate cell activation. J. Hepatol. 63, 679–688 (2015).

48. Yu, X., Wang, Y., Song, Y., Gao, X. & Deng, H. AP-1 is a regulatory transcription factor of inflammaging in the murine kidney and liver. Aging Cell 22, e13858 (2023).

49. Martinez-Jimenez, C. P. et al. Aging increases cell-to-cell transcriptional variability upon immune stimulation. Science 355, 1433–1436 (2017).

50. Yang, Y. et al. A measure of transcriptional dyscoordination for quantifying aging in single cells. Preprint at bioRxiv 10.64898/2026.01.24.701460 (2026).

51. Amaral, M. L. et al. Single-cell epigenomics uncovers heterochromatin instability and transcription factor dysfunction during mouse brain aging. Cell Rep. 45, 117073 (2026).

52. Zemke, N. R. et al. Epigenetic and 3D genome reprogramming during the aging of the human hippocampus. Science 393, eadt8307 (2026).

53. Ma, F. & Sen, R. Physiological aging in three dimensions. Trends Cell Biol. 36, 230–245 (2026).

54. Yang, J.-H. et al. Loss of epigenetic information as a cause of mammalian aging. Cell 186, 305–326.e27 (2023).

55. Padeken, J., Methot, S. P. & Gasser, S. M. Establishment of H3K9-methylated heterochromatin and its functions in tissue differentiation and maintenance. Nat. Rev. Mol. Cell Biol. 23, 623–640 (2022).

56. Bulut-Karslioglu, A. et al. Suv39h-dependent H3K9me3 marks intact retrotransposons and silences LINE elements in mouse embryonic stem cells. Mol. Cell 55, 277–290 (2014).

57. Mrabti, C. et al. Loss of H3K9 trimethylation leads to premature aging. Preprint at bioRxiv 10.1101/2024.07.24.604929 (2024).

58. Tominaga, K. & Suzuki, H. I. TGF-β signaling in cellular senescence and aging-related pathology. Int. J. Mol. Sci. 20, 5002 (2019).

59. Mittelbrunn, M. & Kroemer, G. Hallmarks of T cell aging. Nat. Immunol. 22, 687–698 (2021).

60. Montagner, A. et al. Liver PPARα is crucial for whole-body fatty acid homeostasis and is protective against NAFLD. Gut 65, 1202–1214 (2016).

61. Lee, B. Y. et al. Senescence-associated β-galactosidase is lysosomal β-galactosidase. Aging Cell 5, 187–195 (2006).

62. Bonten, E. J., Annunziata, I. & d’Azzo, A. Lysosomal multienzyme complex: pros and cons of working together. Cell. Mol. Life Sci. 71, 2017–2032 (2014).

63. Gunewardena, S. et al. Progressive loss of hepatocyte nuclear factor 4 alpha activity in chronic liver diseases in humans. Hepatology 76, 372–386 (2022).

64. Ito, T., Teo, Y. V., Evans, S. A., Neretti, N. & Sedivy, J. M. Regulation of cellular senescence by Polycomb chromatin modifiers through distinct DNA damage- and histone methylation-dependent pathways. Cell Rep. 22, 3480–3492 (2018).

65. Schafer, M. J. et al. Senotypes define the diverse landscape of senescent cells. Nat. Aging 6, 1771–1780 (2026).

66. Umbaugh, D. S., Nguyen, N. T., Smith, S. H., Ramachandran, A. & Jaeschke, H. The p21+ perinecrotic hepatocytes produce the chemokine CXCL14 after a severe acetaminophen overdose promoting hepatocyte injury and delaying regeneration. Toxicology 504, 153804 (2024).

67. Russell, J. O. & Monga, S. P. Wnt/β-catenin signaling in liver development, homeostasis, and pathobiology. Annu. Rev. Pathol. 13, 351–378 (2018).

68. Cole, J. J. et al. Diverse interventions that extend mouse lifespan suppress shared age-associated epigenetic changes at critical gene regulatory regions. Genome Biol. 18, 58 (2017).

69. Gatticchi, L. et al. Tm7sf2 Disruption Alters Radial Gene Positioning in Mouse Liver Leading to Metabolic Defects and Diabetes Characteristics. Front. Cell Dev. Biol. 8, 592573 (2020).

70. Guelen, L. et al. Domain organization of human chromosomes revealed by mapping of nuclear lamina interactions. Nature 453, 948–951 (2008).

71. Cruickshanks, H. A. et al. Senescent cells harbour features of the cancer epigenome. Nat. Cell Biol. 15, 1495–1506 (2013).

72. Subramanian, A., et al. Gene set enrichment analysis: a knowledge-based approach for interpreting genome-wide expression profiles. Proc. Natl. Acad. Sci. U.S.A. 102, 15545–15550 (2005).

73. Hänzelmann, S., Castelo, R. & Guinney, J. GSVA: gene set variation analysis for microarray and RNA-seq data. BMC Bioinformatics 14, 7 (2013).

74. Miller, K. N. et al. p53 enhances DNA repair and suppresses cytoplasmic chromatin fragments and inflammation in senescent cells. Nat. Commun. 16, 2229 (2025).

75. Teneche, M. G. et al. p53 supports healthy aging. Preprint at bioRxiv 10.64898/2026.11.16.766897 (2026).

76. Lustig, B. et al. Negative feedback loop of Wnt signaling through upregulation of conductin/axin2 in colorectal and liver tumors. Mol. Cell. Biol. 22, 1184–1193 (2002).

77. Jho, E. et al. Wnt/beta-catenin/Tcf signaling induces the transcription of Axin2, a negative regulator of the signaling pathway. Mol. Cell. Biol. 22, 1172–1183 (2002).

78. Kakugawa, S. et al. Notum deacylates Wnt proteins to suppress signalling activity. Nature 519, 187–192 (2015).

79. Jin, S. et al. Inference and analysis of cell-cell communication using CellChat. Nat. Commun. 12, 1088 (2021).

80. Price, A. J., Manjegowda, M. C., Kain, J., Anandh, S. & Bochkis, I. M. Hdac3, Setdb1, and Kap1 mark H3K9me3/H3K14ac bivalent regions in young and aged liver. Aging Cell 19, e13092 (2020).

81. Acosta, J. C. et al. A complex secretory program orchestrated by the inflammasome controls paracrine senescence. Nat. Cell Biol. 15, 978–990 (2013).

82. Fleming, S. J. et al. Unsupervised removal of systematic background noise from droplet-based single-cell experiments using CellBender. Nat. Methods. 20, 1323–1335 (2023).

83. McGinnis, C. S., Murrow, L. M. & Gartner, Z. J. DoubletFinder: doublet detection in single-cell RNA sequencing data using artificial nearest neighbors. Cell Syst. 8, 329–337.e4 (2019).

84. Hao, Y. et al. Dictionary learning for integrative, multimodal and scalable single-cell analysis. Nat. Biotechnol. 42, 293–304 (2024).

85. Stuart, T., Srivastava, A., Madad, S., Lareau, C. A. & Satija, R. Single-cell chromatin state analysis with Signac. Nat. Methods. 18, 1333–1341 (2021).

86. Hafemeister, C. & Satija, R. Normalization and variance stabilization of single-cell RNA-seq data using regularized negative binomial regression. Genome Biol. 20, 296 (2019).

87. Choudhary, S. & Satija, R. Comparison and evaluation of statistical error models for scRNA-seq. Genome Biol. 23, 27 (2022).

88. Korsunsky, I. et al. Fast, sensitive and accurate integration of single-cell data with Harmony. Nat. Methods. 16, 1289–1296 (2019).

89. Doherty, P. et al. MACS3: a peak-calling platform for bulk and single-cell regulatory genomics. Genomics Proteomics Bioinformatics qzag097 (2026) doi:10.1093/gpbjnl/qzag097.

90. Zhang, Y. et al. Model-based analysis of ChIP-Seq (MACS). Genome Biol. 9, R137 (2008).

91. Amemiya, H. M., Kundaje, A. & Boyle, A. P. The ENCODE blacklist: identification of problematic regions of the genome. Sci. Rep. 9, 9354 (2019).

92. Li, Y. E. et al. An atlas of gene regulatory elements in adult mouse cerebrum. Nature 598, 129–136 (2021).

93. Liu, H. et al. DNA methylation atlas of the mouse brain at single-cell resolution. Nature 598, 120–128 (2021).

94. Zemke, N. R. et al. Conserved and divergent gene regulatory programs of the mammalian neocortex. Nature 624, 390–402 (2023).

95. Krueger, F. & Andrews, S. R. Bismark: a flexible aligner and methylation caller for Bisulfite-Seq applications. Bioinformatics 27, 1571–1572 (2011).

96. Langmead, B., Trapnell, C., Pop, M. & Salzberg, S. L. Ultrafast and memory-efficient alignment of short DNA sequences to the human genome. Genome Biol. 10, R25 (2009).

97. Li, H. et al. The Sequence Alignment/Map format and SAMtools. Bioinformatics 25, 2078–2079 (2009).

98. Broad Institute. Picard tools. https://broadinstitute.github.io/picard/ (accessed 16 November 2026).

99. Cusanovich, D. A. et al. Multiplex single-cell profiling of chromatin accessibility by combinatorial cellular indexing. Science 348, 910–914 (2015).

100. Traag, V. A., Waltman, L. & Van Eck, N. J. From Louvain to Leiden: guaranteeing well-connected communities. Sci. Rep. 9, 5233 (2019).

101. McInnes, L., Healy, J., Saul, N. & Großberger, L. UMAP: uniform manifold approximation and projection. J. Open Source Softw. 3, 861 (2018).

102. Lee, D.-S. et al. Simultaneous profiling of 3D genome structure and DNA methylation in single human cells. Nat. Methods. 16, 999–1006 (2019).

103. Dong, W., Moses, C. & Li, K. Efficient k-nearest neighbor graph construction for generic similarity measures. in Proceedings of the 20th international conference on World wide web 577–586 (ACM, Hyderabad India, 2011). doi:10.1145/1963405.1963487.

104. Zhou, J., et al. Robust single-cell Hi-C clustering by convolution- and random-walk–based imputation. Proc. Natl. Acad. Sci. U.S.A. 116, 14011–14018 (2019).

105. Abdennur, N. & Mirny, L. A. Cooler: scalable storage for Hi-C data and other genomically labeled arrays. Bioinformatics 36, 311–316 (2020).

106. Imakaev, M. et al. Iterative correction of Hi-C data reveals hallmarks of chromosome organization. Nat. Methods 9, 999–1003 (2012).

107. Ramírez, F. et al. High-resolution TADs reveal DNA sequences underlying genome organization in flies. Nat. Commun. 9, 189 (2018).

108. Zhu, C. et al. Joint profiling of histone modifications and transcriptome in single cells from mouse brain. Nat. Methods. 18, 283–292 (2021).

109. Xie, Y. et al. Droplet-based single-cell joint profiling of histone modifications and transcriptomes. Nat. Struct. Mol. Biol. 30, 1428–1433 (2023).

110. Blondel, V. D., Guillaume, J.-L., Lambiotte, R. & Lefebvre, E. Fast unfolding of communities in large networks. J. Stat. Mech. 2008, P10008 (2008).

111. Quinlan, A. R. & Hall, I. M. BEDTools: a flexible suite of utilities for comparing genomic features. Bioinformatics 26, 841–842 (2010).

112. Heinz, S. et al. Simple combinations of lineage-determining transcription factors prime cis-regulatory elements required for macrophage and B cell identities. Mol. Cell 38, 576–589 (2010).

113. Wu, T. et al. clusterProfiler 4.0: A universal enrichment tool for interpreting omics data. The Innovation 2, 100141 (2021).

114. Finak, G. et al. MAST: a flexible statistical framework for assessing transcriptional changes and characterizing heterogeneity in single-cell RNA sequencing data. Genome Biol. 16, 278 (2015).

115. Ernst, J. & Kellis, M. ChromHMM: automating chromatin-state discovery and characterization. Nat. Methods. 9, 215–216 (2012).

116. Pedregosa, F. et al. Scikit-learn: machine learning in Python. J. Mach. Learn. Res. 12, 2825–2830 (2011).

117. Ester, M., Kriegel, H.-P., Sander, J. & Xu, X. A density-based algorithm for discovering clusters in large spatial databases with noise. In Proceedings of the Second International Conference on Knowledge Discovery and Data Mining 226–231 (AAAI Press, 1996).

118. Gschwind, A. R. et al. An encyclopedia of human enhancer–gene regulatory interactions. Nature 657, 179–190 (2026).

119. Stringer, C., Wang, T., Michaelos, M. & Pachitariu, M. Cellpose: a generalist algorithm for cellular segmentation. Nat. Methods 18, 100–106 (2021).

120. Stuart, T. et al. Comprehensive integration of single-cell data. Cell 177, 1888–1902.e21 (2019).

121. Zhang, M. et al. Molecularly defined and spatially resolved cell atlas of the whole mouse brain. Nature 624, 343–354 (2023).

122. Hildebrandt, F. et al. Spatial Transcriptomics to define transcriptional patterns of zonation and structural components in the mouse liver. Nat. Commun. 12, 7046 (2021).

123. Berndt, N. et al. Functional Consequences of Metabolic Zonation in Murine Livers: Insights for an Old Story. Hepatology 73, 795–810 (2021).

124. Chen, S., Zhou, Y., Chen, Y. & Gu, J. fastp: an ultra-fast all-in-one FASTQ preprocessor. Bioinformatics 34, i884–i890 (2018).

125. Dobin, A. et al. STAR: ultrafast universal RNA-seq aligner. Bioinformatics 29, 15–21 (2013).

126. Liu, D. Algorithms for efficiently collapsing reads with unique molecular identifiers. PeerJ 7, e8275 (2019).

127. Ewels, P., Magnusson, M., Lundin, S. & Käller, M. MultiQC: summarize analysis results for multiple tools and samples in a single report. Bioinformatics 32, 3047–3048 (2016).

128. Liao, Y., Smyth, G. K. & Shi, W. featureCounts: an efficient general purpose program for assigning sequence reads to genomic features. Bioinformatics 30, 923–930 (2014).

129. Love, M. I., Huber, W. & Anders, S. Moderated estimation of fold change and dispersion for RNA-seq data with DESeq2. Genome Biol. 15, 550 (2014).

130. Korotkevich, G. et al. Fast gene set enrichment analysis. Preprint at bioRxiv 10.1101/060012 (2021).

131. Castanza, A. S. et al. Extending support for mouse data in the Molecular Signatures Database (MSigDB). Nat. Methods 20, 1619–1620 (2023).

132. Schindelin, J. et al. Fiji: an open-source platform for biological-image analysis. Nat. Methods 9, 676–682 (2012).

133. Ramírez, F. et al. deepTools2: a next generation web server for deep-sequencing data analysis. Nucleic Acids Res 44, W160–W165 (2016).

